# Next-Gen sequencing of novel pandemic swine flu [A(H1N1)pdm09] virus in India revealed novel mutations across the genome

**DOI:** 10.1101/693358

**Authors:** Paban Kumar Dash, Chitra Pattabiraman, Kundan Tandel, Shashi Sharma, Jyoti S Kumar, Shilpa Siddappa, Malali Gowda, Sudhir Krishna, Man Mohan Parida

**Author notes:** **Corresponding author**, Dr M.M.Parida, Scientist -G, Head, Division of Virology, Defence R & D Establishment, Jhansi Road, Gwalior – 474002, India.

## Abstract

The Influenza A H1N1 virus of 2009 was the first pandemic flu virus of the 21^st^ century. Identifying the emergence of mutations in rapidly mutating Influenza viruses that allow increased transmission or confer resistance are invaluable to global outbreak response. Here we recovered 5 complete Influenza A genomes from 4 oropharygeal swabs and one cell culture isolate from a severe Indian outbreak of flu in early 2015. Multiple amino acids substitutions including those known to confer resistance to Oseltamivir and increased pathogenecity in mice were found in the Neuraminidase gene. Additional mutations both reported and novel were found throughout the genome compared to the vaccine strain (California/04/2009). All eight segments of the complete genomes were found to be genetically related to the 2009 pandemic strain, A(H1N1)pdm09 and belonging to the emerging genogroup 6B. This group was found to be of south East Asian origin by time scale phylogentic analysis. A phylogeographic analysis revealed 39 significant migration events among globally circulating viruses. This study is the first extensive complete genome and phylogeographic analysis of 2015 Indian A(H1N1) pdm09 viruses. We report several novel mutations in the 2015 Indian strains which need to be evaluated for effect on viral replication, transmission and resistance to therapy. The identification of mutant A(H1N1)pdm09 from India warrants continuous monitoring of viral evolution for implementation of suitable medical countermeasures.

## 1. Introduction

The last pandemic of influenza occurred when a quadrouple reassorted Influenza A virus which originated in Mexico spread across the human population in 2009. This novel pandemic virus was found to originate from multiple reassortments of avian, swine, and human viruses and probably circulated in the north American swine population for over a decade (Potdar et al., 2010).

The genome of IAV comprises of 8 segments consisting of 8 negative-sense singlestranded RNA genomes. The 13.5 kb genome encodes 12 proteins: hemagglutinin (HA), M proteins (M1 and M2), neuraminidase (NA), nucleocapsid protein (NP), nonstructural proteins (NS1 and NS2), and polymerase subunits (PA, PA-X, PB1-F1, PB1-F2, and PB2). IAVs are divided into multiple subtypes based on two surface glycoproteins hemagglutinin (H1 to H18) and neuraminidase (N1 to N11) (Matsuzaki et al., 2014; Garten et al., 2009)

The evolution of Influenza viruses, driven by antigenic drift and antigenic shift allows them to circumvent host immunity and leads to occurrence of regular epidemic and occasional pandemic. Work on multiple influenza viruses has demonstrated that mutations in the receptor binding domain of the influenza hemaglutinin protein are one of the key antigenic determinants together with the viral neuraminidase (Park et al., 2009). Characterization of NA further allows identification of drug resistant mutants particularly to widely used <u>zanamivir</u> (Relenza) and <u>oseltamivir</u> (Tamiflu). Novel variations in HA and NA will lead to changes in the antigenic profile and the consequent inadequateness of immune responses to the virus could give rise to more virulent strains (Kosoltanapiwat et al., 2014). In contrast, relatively little is known about mutations in the other 6 segments of the influenza genome. While more recent evidence does suggest a role of mutations in the matrix proteins in pathogenecity, the information on novel mutations in the full-length virus, especially in geographical niches remains poor.

In 2015, India saw a large number of human flu cases requiring hospitalization. During the pandemic period (2009-2010), more than 50000 cases with a case fatality of 6% were reported in India. Subsequently, the virus activity subsided during 2011-2014 (Parida et al., 2016; Takashita et al., 2015; Dakhave et al., 2013, Broor et al., 2012). Surprisingly, resurgence of A(H1N1)pdm09 was recorded with more severity than the previous pandemic period in early part of 2015. This nation–wide epidemic affected more than 39000 persons with a case fatality of more than 2,500. This outbreak was characterized by viruses which belong to the emerging groups of H1N1 – genogroup 6B. We have recently published HA gene sequence of some of the viruses circulating in 2015, which highlighted the need to characterize the full-genome of these viruses. Here we recovered complete genomes of viruses from this outbreak both from oral swabs and from an isolate and identified potential amino acid changes throughout the viral genome. A time scale phylogeny was constructed to understand their evolution particularly focussing on the emerging genogroup 6B, since the onset of pandemic in 2009.

## 2. Material and Methods

### 2.1 Clinical samples

Acute phase respiratory swab (nasopharyngeal/ throat/ nasal) samples from patients suspected for A(H1N1)pdm09 infection were collected by health administrators of different district of Madhya Pradesh, India. These were referred to DRDE, Gwalior, a nodal reference lab for pandemic swine flu H1N1 molecular diagnosis. WHO guidelines were followed for sample collection, transport and laboratory investigation in a BSL-3 laboratory (WHO, 2009). The detailed clinical history and epidemiological information of the five samples sequenced in this study were provided in Table 1. All these samples were collected at the height of epidemic during Feb-Mar 2015. Ethical clearance for this study was approved by DRDE Biosafety Committee. Informed consent was obtained by district health authorities as part of laboratory diagnosis and treatment.

**Table 1:** Detailed patient profile of pandemic swine flu H1N1 investigated in this study

| Virus ID | Sex | Age<br>(Years) | Location | Category | Treatment<br>taken | Ct<br>value | Date of<br>sample<br>collection | Day since<br>onset of<br>symptoms | X – Ray Findings |
| --- | --- | --- | --- | --- | --- | --- | --- | --- | --- |
| A/India/DRDE_GWL812/2015 | F | 55 | BHOPAL | C | Yes | 24 | 9 Mar 16 | 08 Days | B/L Homogenous<br>Opacities |
| A/India/DRDE_GWL84/2015 | M | 35 | BHOPAL | C | Yes | 26 | 17 Mar 16 | 08 Days | B/L Pneumonitis |
| A/India/DRDE_GWL672/2015 | F | 42 | BHOPAL | C | No | 24 | 05 Mar 16 | 04 Days | B/L Homogenous<br>Opacities |
| A/India/DRDE_GWL897/2015 | M | 56 | GWALIOR | A | Yes | 23 | 14 Mar 16 | 04 Days | Nil |
| A/India/DRDE_GWL989/2015 | F | 09 | SEHORE | B | Yes | 23 | 18 Mar 16 | 03 Days | B/L Hyperinflation<br>with Pneumonitis<br>changes |

### 2.2 Extraction of viral RNA

Viral RNA was extracted from the infected nasopharyngeal/throat/nasal samples using the QIAamp viral RNA mini kit (Qiagen, Germany), according to the manufacturer’s protocol, Briefly 560 μl of buffer AVL was added to 140 μl of infected material and vortexed for 15 sec and the mixture was incubated at room temperature (15^0^C-25^0^C) for 10 min. After washing twice finally RNA was eluted in 70 μl of elution buffer in collection tube and stored at −80 0C until use.

### 2.3 Metagenomic sequencing from cell culture supernatants and oropharngeal samples

RNA from 6 clinical samples, 3 isolates (passaged in MDCK cells) and 1 healthy control was subjected to deep sequencing on the Ion Proton/PGM sequencing platform. Extracted RNA was concentrated and used for library preparation using the Ion Proton Library preparation kit (Ion Total RNA-Seq kit V2), samples were multiplexed in set of 5-6 using IonXpress barcodes (Ion Xpress RNA Seq BC) and template preparation was carried out using Ion PI HI-Q template OT2 200 kit. A water control was included in one chip. Sequencing was carried out with 8 pM of pooled libaries using the Ion PI HIQ 200 kit and pre-primed chips for sequencing (Ion PI chip kit V3). The number of sequences from each sample are provided in Table 2.

**Table 2:**
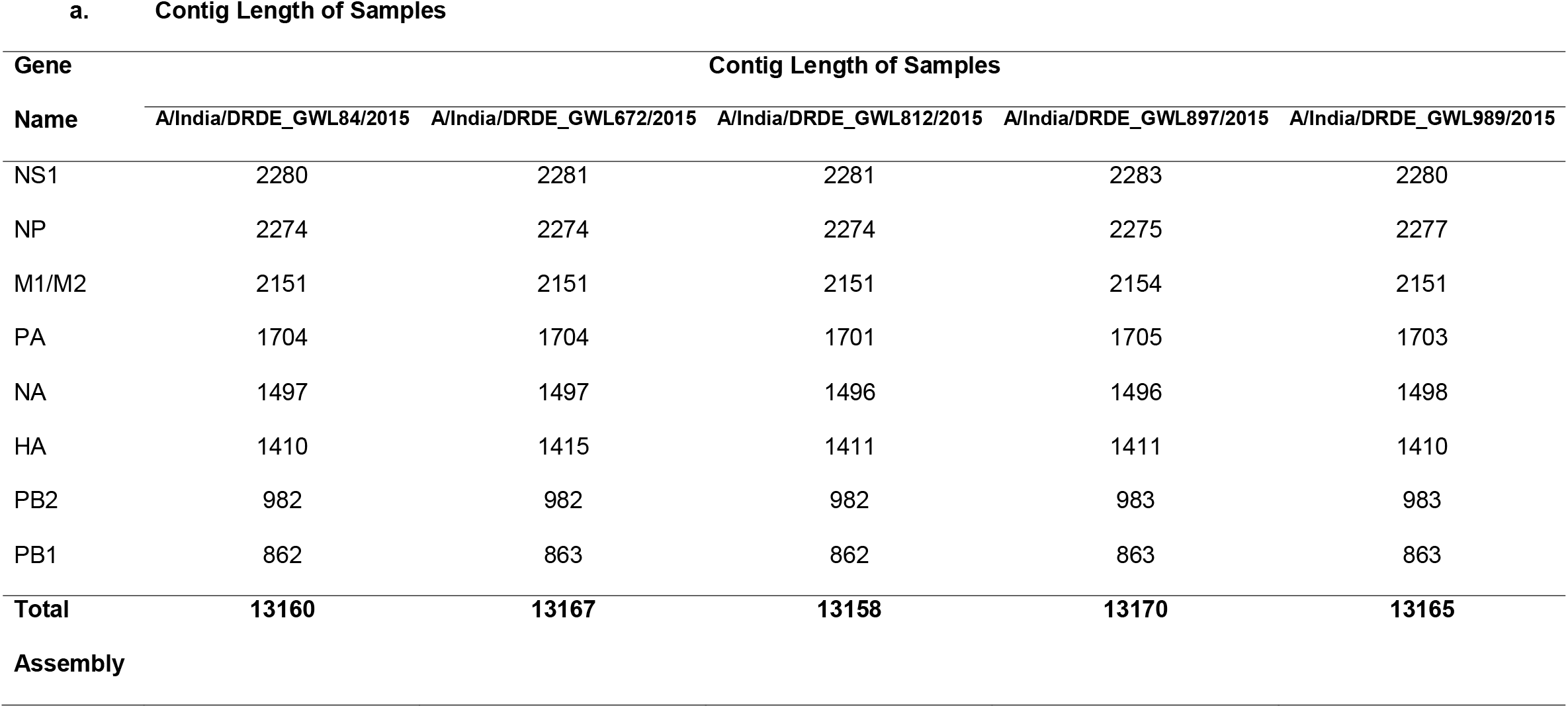
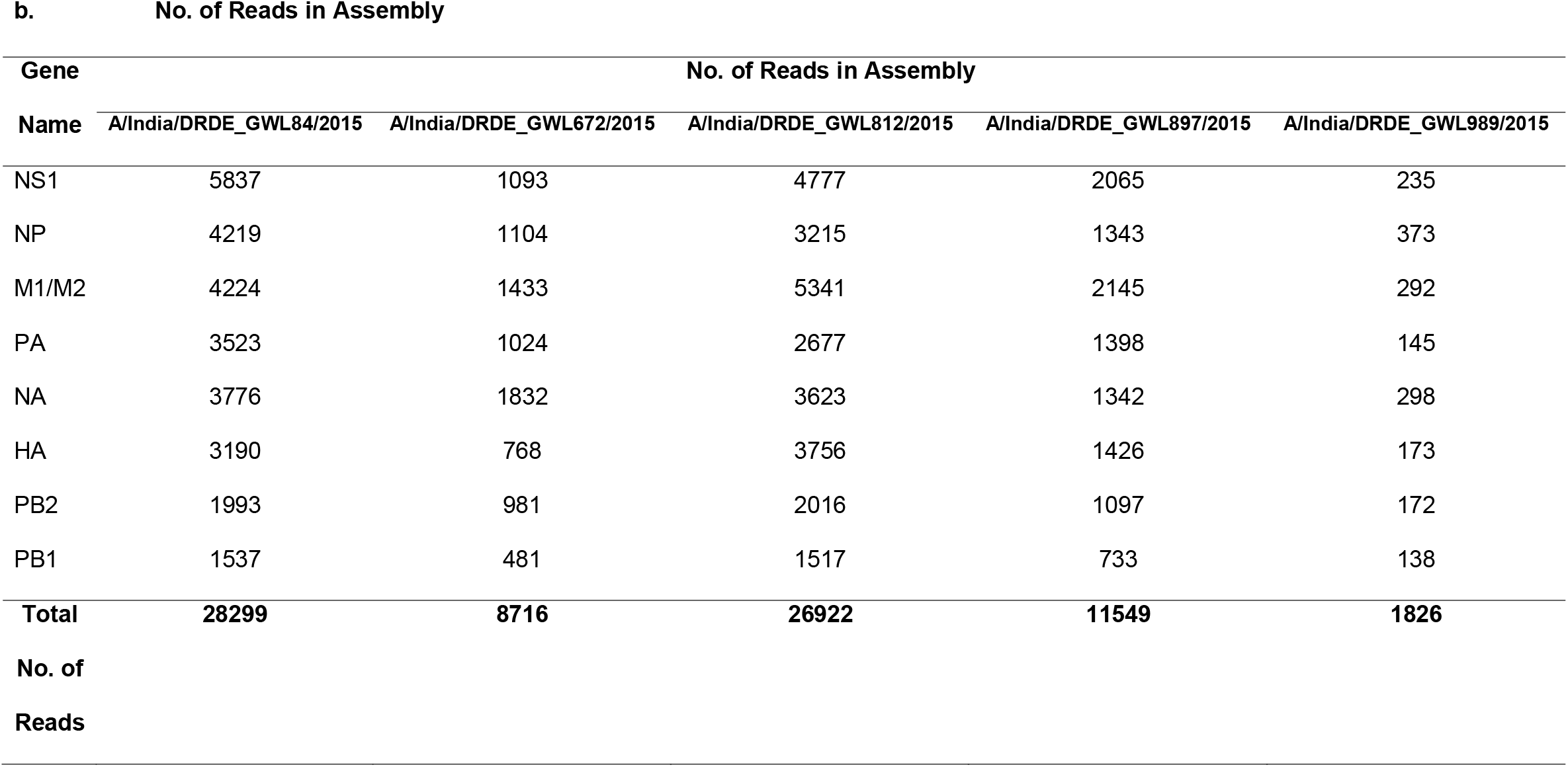
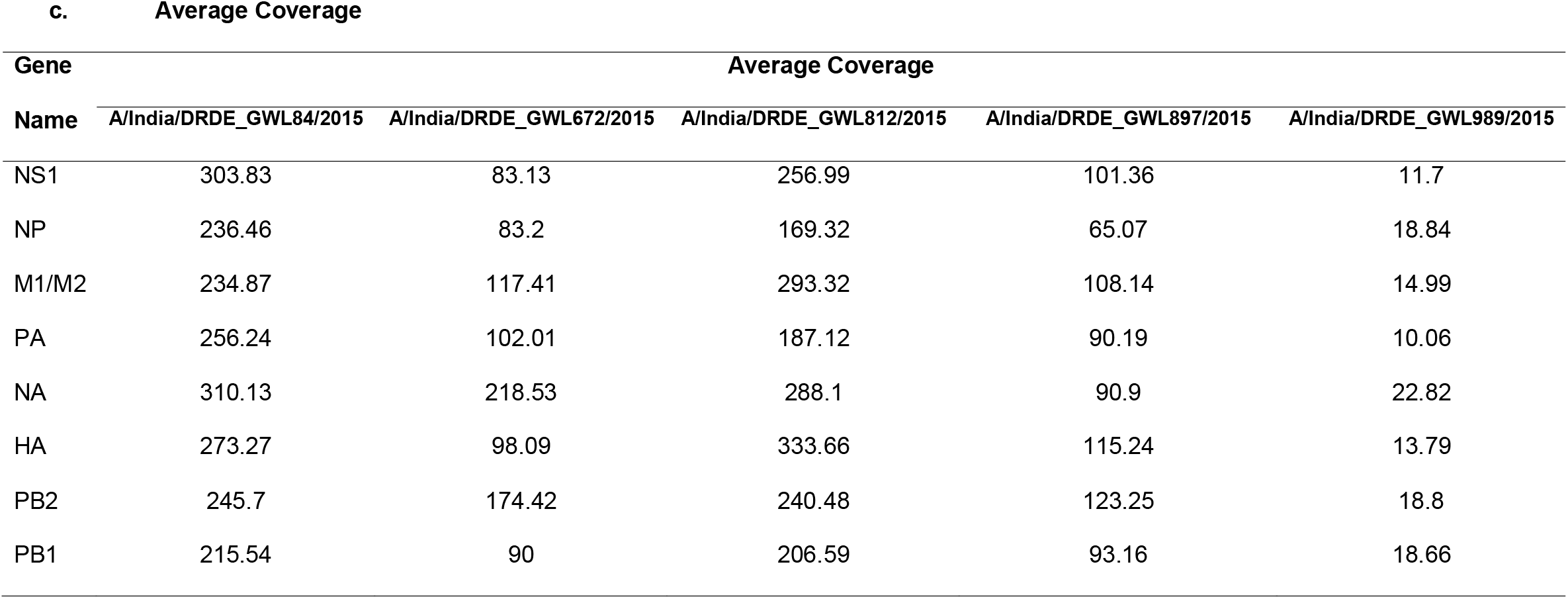
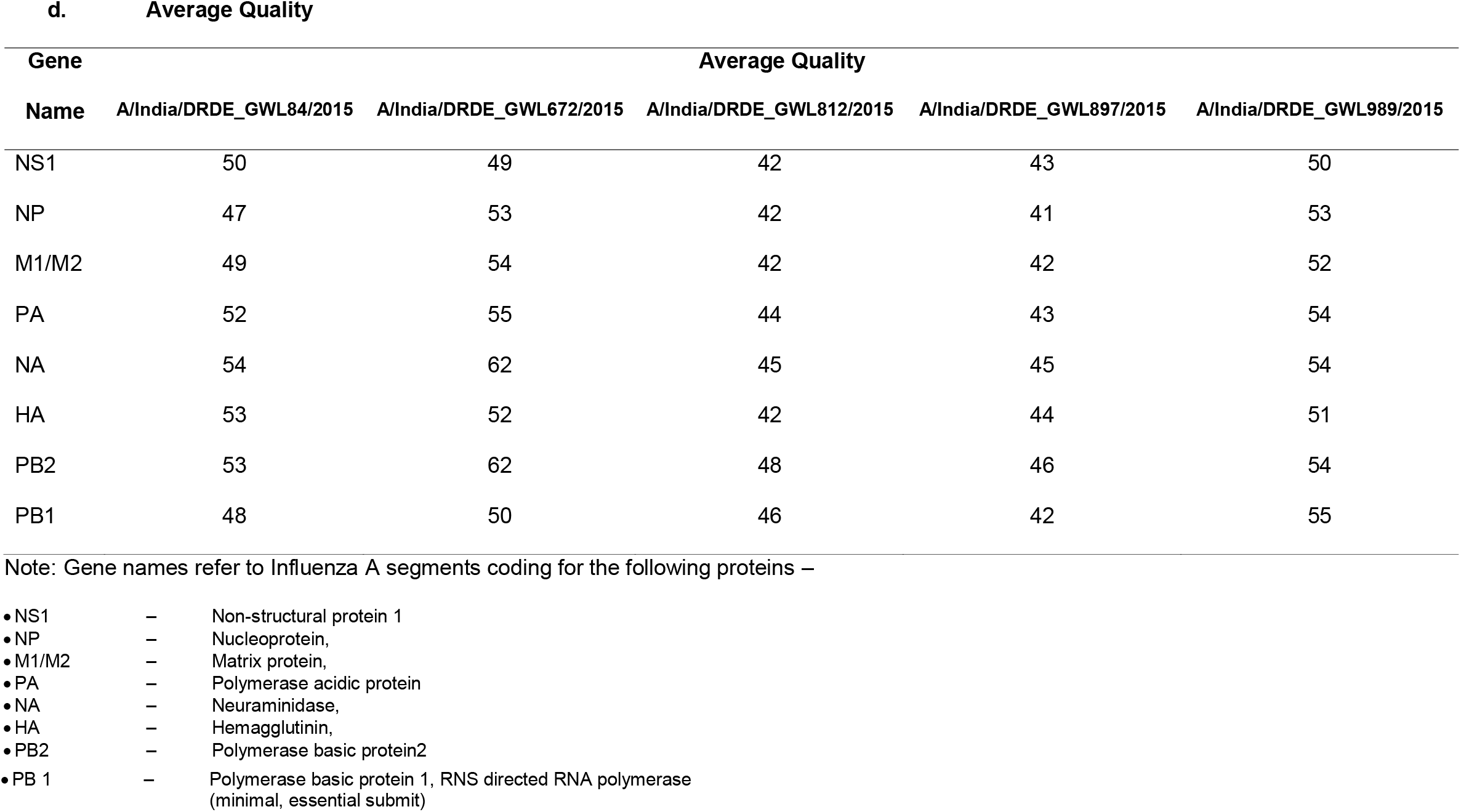
Contig statistics of mapping assemblies

### 2.4 Analysis of sequencing reads

The reads were retrieved after demultiplexing in fastq format. The SNAP alignment tool was used to remove host sequences (human genome and mRNA for samples and canine genome for isolates) and ribosomal RNA sequences (From the SILVA database). The remaining reads were aligned using blast to the viral RefSeq database to provide an unbiased spectrum of viruses. Mapping assembly was performed using the MIRA tool (Chevreux et al., 2000) with the A/California/07/2009(H1N1) strain as the reference strain. The resulting consensus sequence was used in all other analysis. The five viral genome sequences reported here were deposited in GenBank under the accession numbers KX078482 to KX078489, KX078490 to KX078497, KX078498 to KX078505, KX078506 to KX078513, and KX078514 to KX078521 (Dash et al., 2018).

Representative sequences of A(H1N1)pdm09 from geographically diverse region circulating between 2009 and 2015 were retrieved from GISAID EpiFlu database (Table S1). Multiple sequence alignment of the viruses was carried out employing MUSCLE (Edgar, 2004) hosted at the EMBL-EBI server. The deduced amino acid was determined from the nucleotide sequence using the EditSeq module of Lasergene 5 software package (DNASTAR Inc, USA). The deduced amino acid sequences of five A(H1N1)pdm09 sequenced in this study along with prototype vaccine strain (A/California/07/2009) and six Indian representative A(H1N1)pdm09 viruses from 2009 to 2014 were compared. The percent nucleotide identity and percent amino acid identity values were calculated as pairwise p-distances. Concatenated (n=96) and individual gene segment (n=120)-wise phylogenetic analysis was carried out using maximum likelihood algorithm employing GTR+G+I nucleotide substitution model in MEGA5 software programme (Tamura et al., 2011). The robustness of the tree topology was assessed by bootstrap analysis of 1000 pseudo-replicates.

### 2.5 Selection pressure analysis

Recombination analysis was carried out using GARD analysis available at open-source Datamonkey web server. Selection pressure analysis acting on the codons of hemagglutinin (HA), neuraminidase (NA), nucleoprotein (NP) and matrix protein (MP:M1,M2) of H1N1pdm virus was carried out using HyPhy open-source software package available under the datamonkey (http://www.datamonkey.org/) (Delport et al., 2010). Analysis was performed using reference sequences [n=115(HA); n=115(NA); n=115(NP) and n=115 (MP:M1, M2)] including globally circulating H1N1pdm virus. The ratio of non-synonnymous (dN) to synonymous (dS) substitutions per site (dN/dS) were estimated using four different approaches including: single likelihood ancestor counting (SLAC), fixed effects likelihood (FEL), mixed effects model of evolution (MEME) and fast unbiased bayesian approximation (FUBAR) (Muerell., 2012). Best nucleotide substitutions model for different data sets as determined through the available tool in Datamonkey server was adopted in the analysis.

### 2.6 Molecular clock analysis

Ninety six complete genome (13027 nt) sequences (including 86 from genogroup 6B) from eleven representative geographical locations were downloaded from the GISAID EpiFlu database were used for molecular clock analysis. The sampling dates ranged from 2009-2015 and sampling locations were Africa (n=8), Caribbean (n=3), Central Asia (n=1), Europe (n=14), Middle East (n=2), North America (n=16), North Asia (n=3), Oceania (n=7), South America (n=4), South Asia (n=20), and Southeast Asia (n=18). All sequences were aligned using Bioedit and MEGA softwares. Best fit model was selected for the analysis of this data set using MEGA version 6 software using model selection application. Time scaled phylogeny construction was performed using Bayesian MCMC approach (Beast v 1.7.4; http://beast.io.ed.ac.uk) (Drummond and Rambaut, 2007) implementing a HKY85 model using both strict and an uncorrelated lognormal relaxed clock model. Bayesian GMRF skyride population model was selected in order to perform discrete phylogeographic analysis (Lemey et al., 2009). MCMC analysis was conducted for 100 million generation and sampled at every 10,000 steps. Convergence was assessed on the basis of effective sampling size (ESS) after a 10% burn-in using Tracer software v1.5 (http://tree.bio.ed.ac.uk/software/tracer). All parameter estimates was indicated by 95% highest posterior density (95% HPD) intervals. The tree was summarized in another application of beast analysis tree annotator program included in the beast package. Final tree was analyzed in Figtree v 1.4.0 for tree visualization.

### 2.7 Phylogeographic analysis

The continuous time markov chain (CTMC) process over discrete sampling locations implemented in BEAST (Drummond and Rambaut, 2007) was used for the phylgeographical analysis, implementing BSSVS (Bayesian stochastic search variable selection) model which allows the diffusion rates to be zero with a positive posterior probability. Comparison of the posterior and prior probability of the individual rates being zero provided a formal BF (Bayes factor) for testing the significant linkage between locations. Rates with a BF of >3 were considered well supported and formed the migration pathway. SPREAD application (Bielejec et al., 2011, http://www.phylogeography.org/SPREAD.html) was used to visualize and convert the estimated divergence time on spatial estimates annotated in the MCC tree to a key hole markup language (KML) file.

## 3. Results

We have previously reported H1N1 testing and HA gene sequencing for a cohort of patients from India in 2015. Here we performed full genome sequencing for 9 of those and got 5 complete genomes (Table 1). We sequenced 6 oropharyngeal swabs confirmed to be H1N1 by RT-PCR, 3 cell culture (in MDCK cells) isolates at passage 1 and 1 healthy oropharngeal swab at the peak of the Flu outbreak of 2015 in India (January-March, 2015). Unbiased metagenomic sequencing was performed with the aim of capturing variant viral genomes as well as other potentially pathogenic viruses. The reads from IonProton sequencing were mapped using the A/California/07/2009 as the reference strain and complete genomes were recovered from 4 swabs and 1 isolate. The average genome length was 13133 bp, with good coverage across 8 segments/genes (Table 2). Contig statistics show an average read depth of about 100X except for the genome from the isolate which had an average read depth of 16X across all segments (Table 2). These five genomes (henceforth called 2015 genomes) were used for all subsequent analysis. The sequences had a similarity of 98.7 −100% similarity among themselves, with maximum sequence divergence in the Neuraminidase gene wiith the A/India/DRDE_GWL812/2015 showing maximum divergence. All 5 genomes were confirmed to be H1N1 and related to the 2009 pandemic strain.

We then compared the 2015 genomes to the prototype A/California/07/2009 and representative Indian isolates from 2009-2014. Amino acid sequence similarity among the set of 12 viruses was between 95.7-100% depending on protein. Many of the amino acid substitutions were found in residues on the surface of the protein (Fig 1.) 11 amino acid substitutions were consistently found in all 5 2015 genomes in the Hemagluttinin protein. 3 of the 11 substitutions were in the antigenic site of HA. (K180Q in Sa, S202T in Sb and S220T in Ca1) (Fig. 2). 13 mutations, common to all 5 2015 genomes were found in Neuraminidase. The characteristic H275Y Oseltamivir resistance mutation was found in one sample (with pre-exposure to the drug). 2, 7 4, 5, 4 and 5 amino acid substitution were identified in PB1, PB2, NP PA, MP and NS protein were consistently identified. Apart from these, multiple unique mutations were also noted in the different strains (Fig 1 and Table 3).

**Fig 1.**
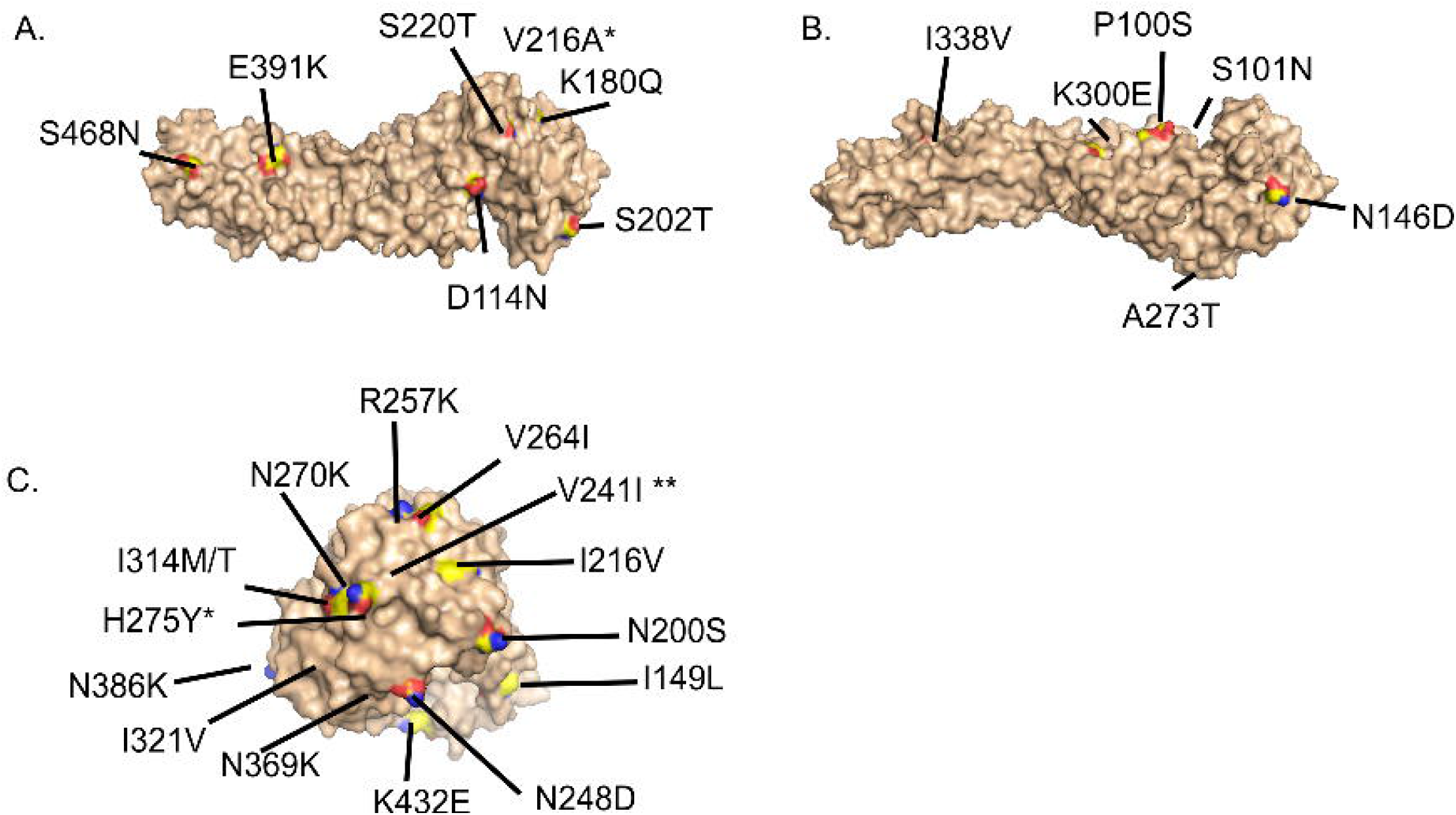
3D structure of prototype A/California/04/2009 strain showing the critical amino acid substitutions. Figure shows the location of amino acid substitutions for Influenza Hemagglutinin (A and B – different rotational views) and Neuraminidase (C). Homology modeling was performed using SWISS-MODEL and the best model for the California/04/2009 sequence was chosen for showing the substitutions. The protein surfaces as visualized by PyMOL are shown in light brown, amino acid that were found to be different in the Gwalior 2015 strains are colored by elements CHNOS. * represents amino acid not found on the surface. ** represent amino acid inside an accessible pocket. 13/16 substitutions for Hemagglutinin and 14/21 substitutions for Neuraminidase are shown, as the other residues were not present in the model.

**Fig 2.**
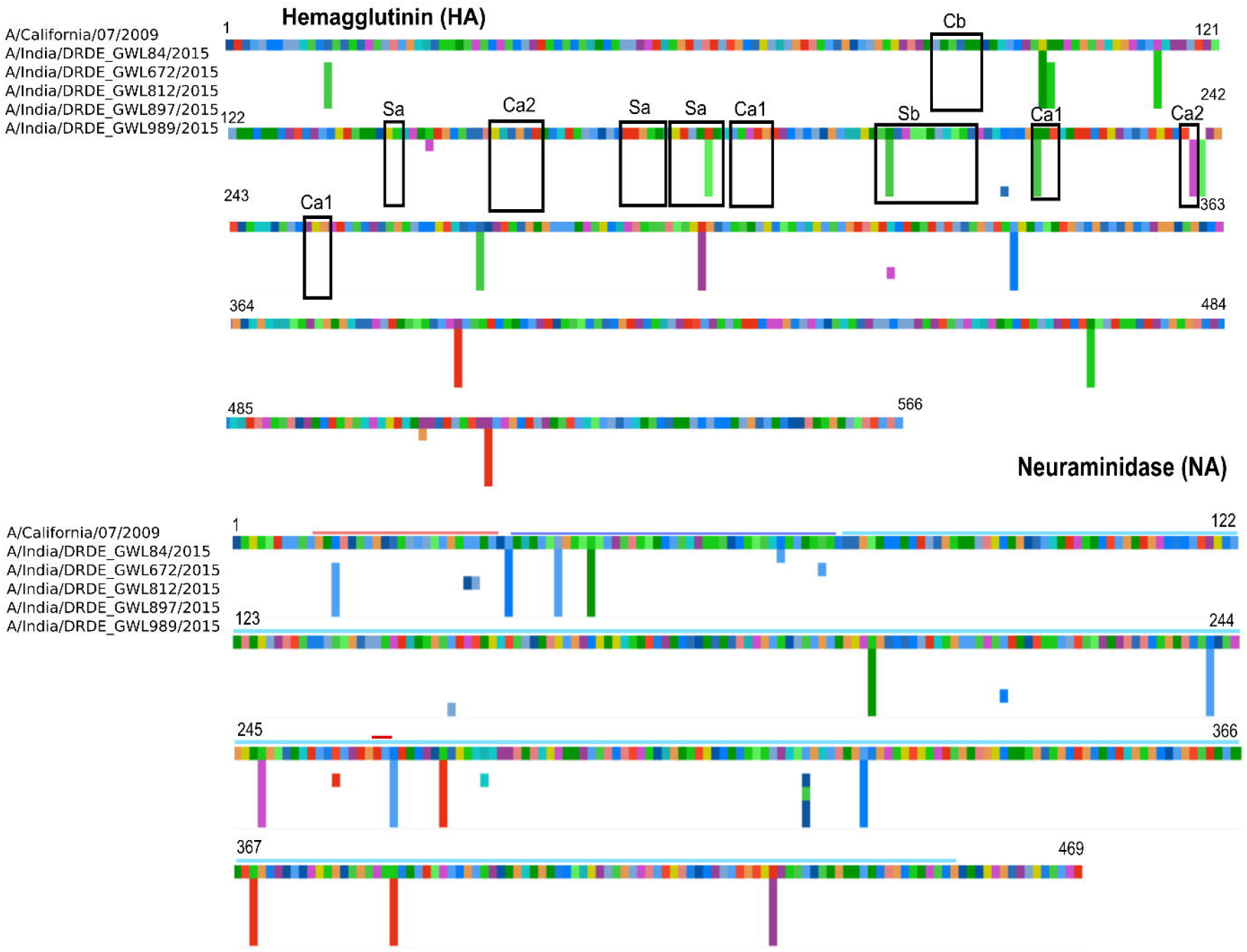
Figure showing amino acid substitutions in HA1 protein and Neuraminidase (NA) of 2015 Indian A(H1N1)pdm09 viruses in comparison to prototype A/California/07/2009 vaccine strain.

**Table 3:**
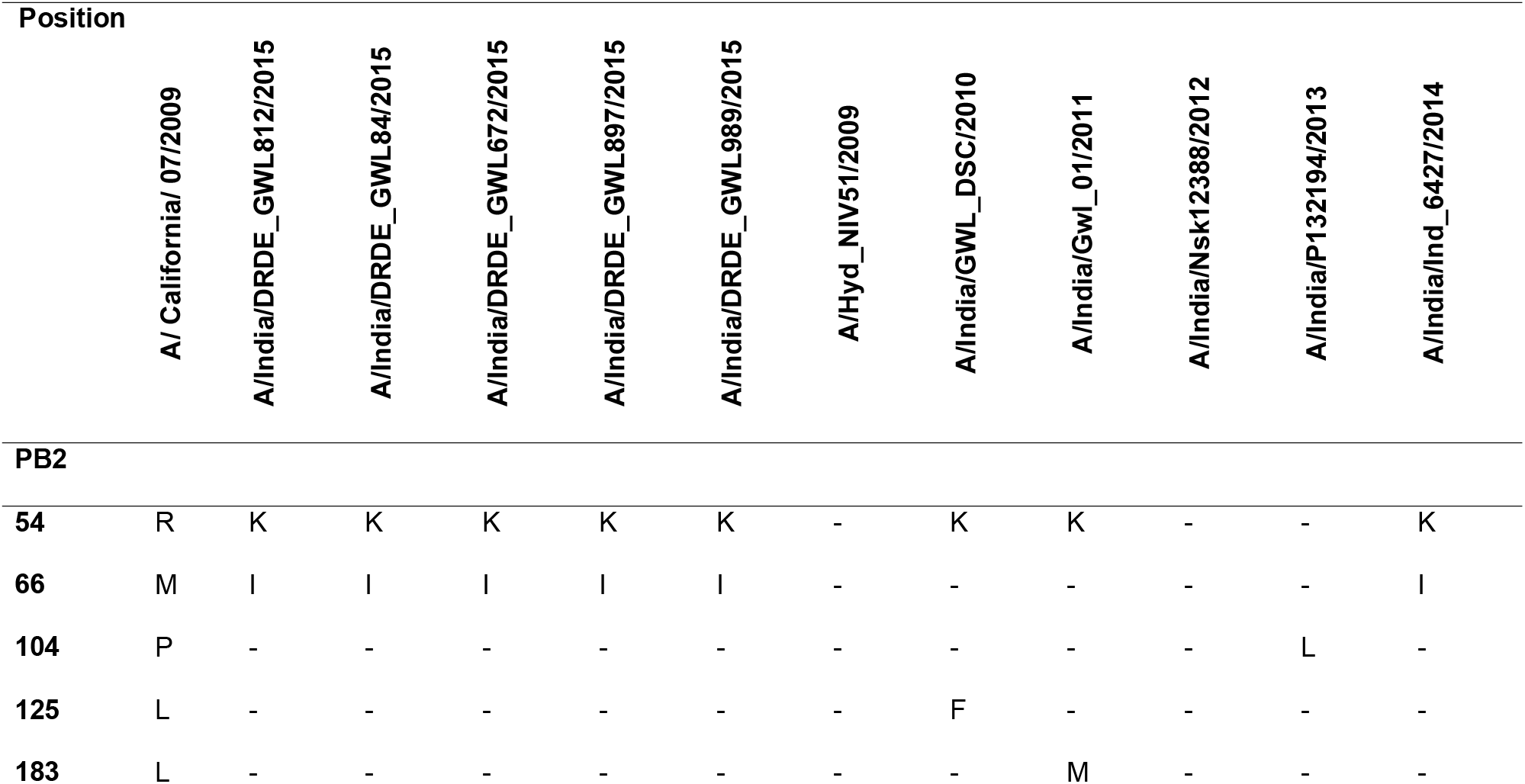
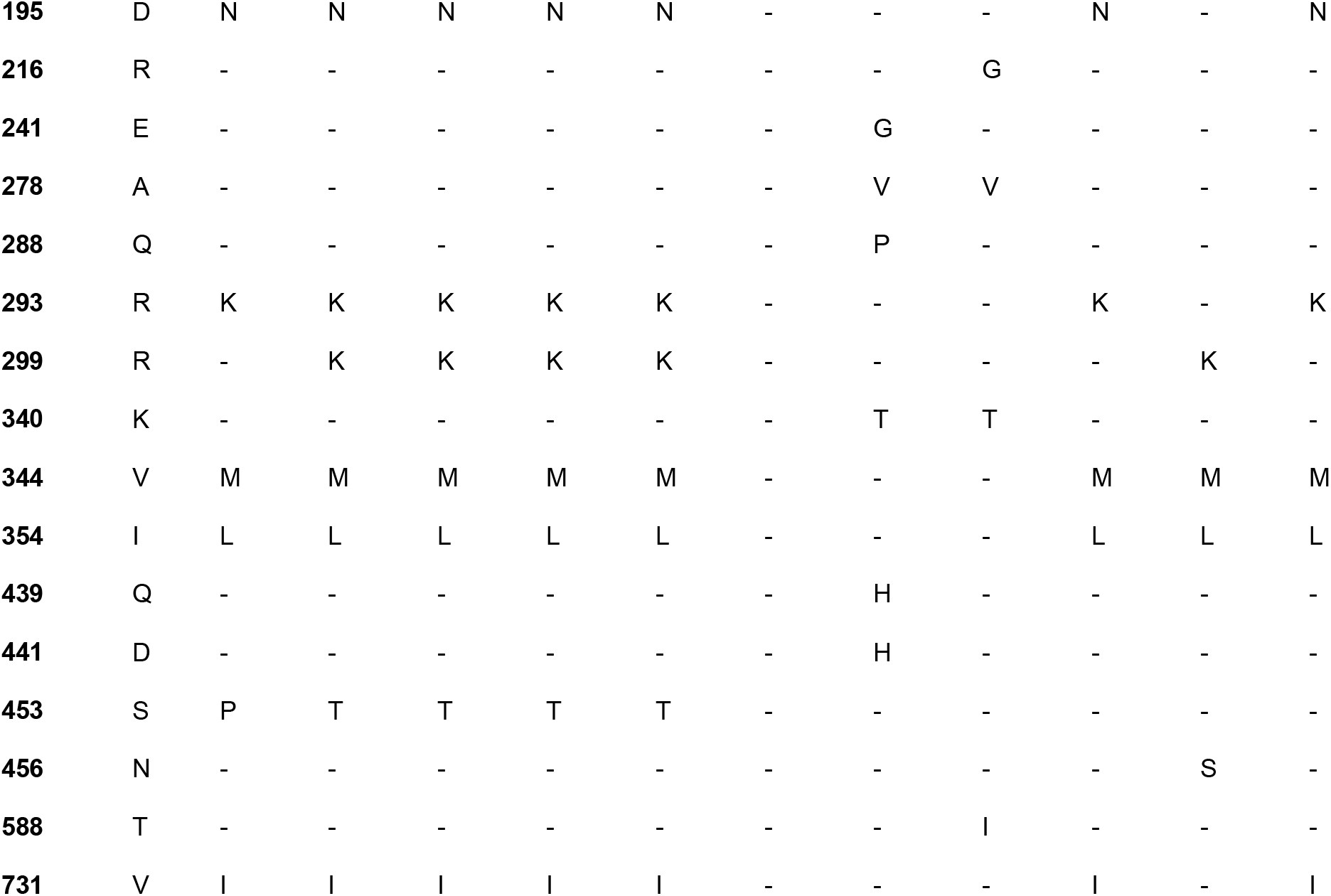
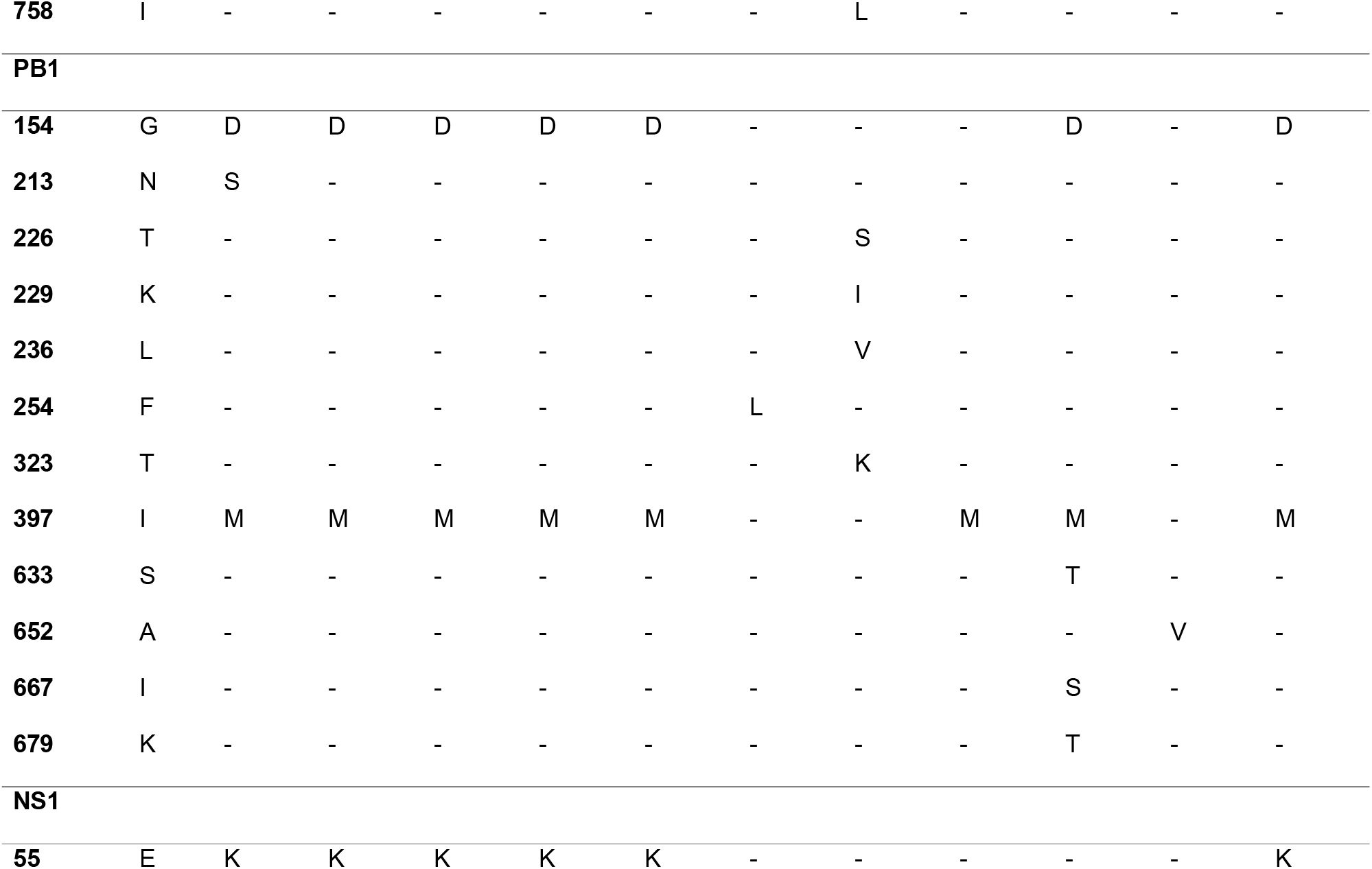
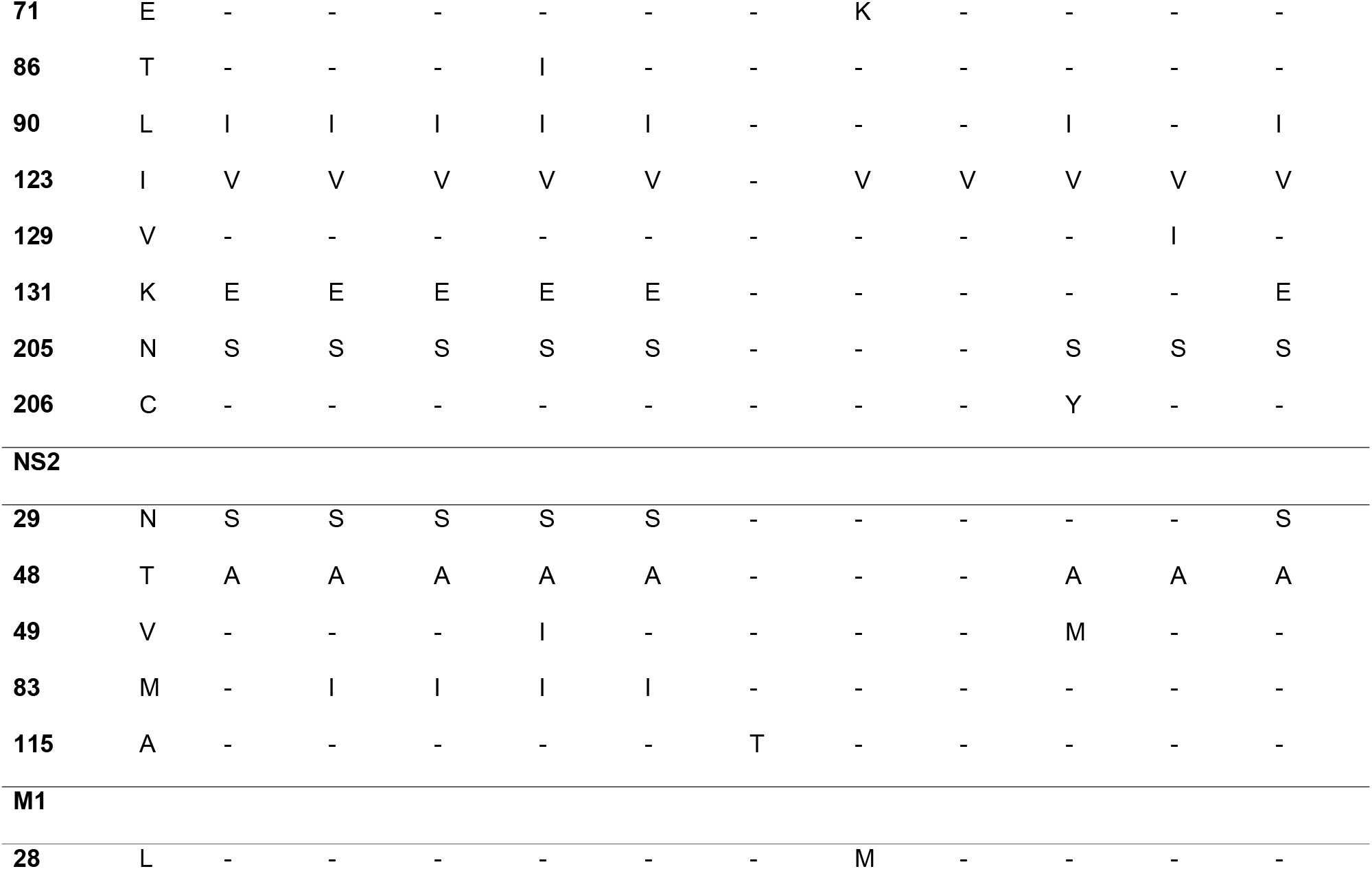
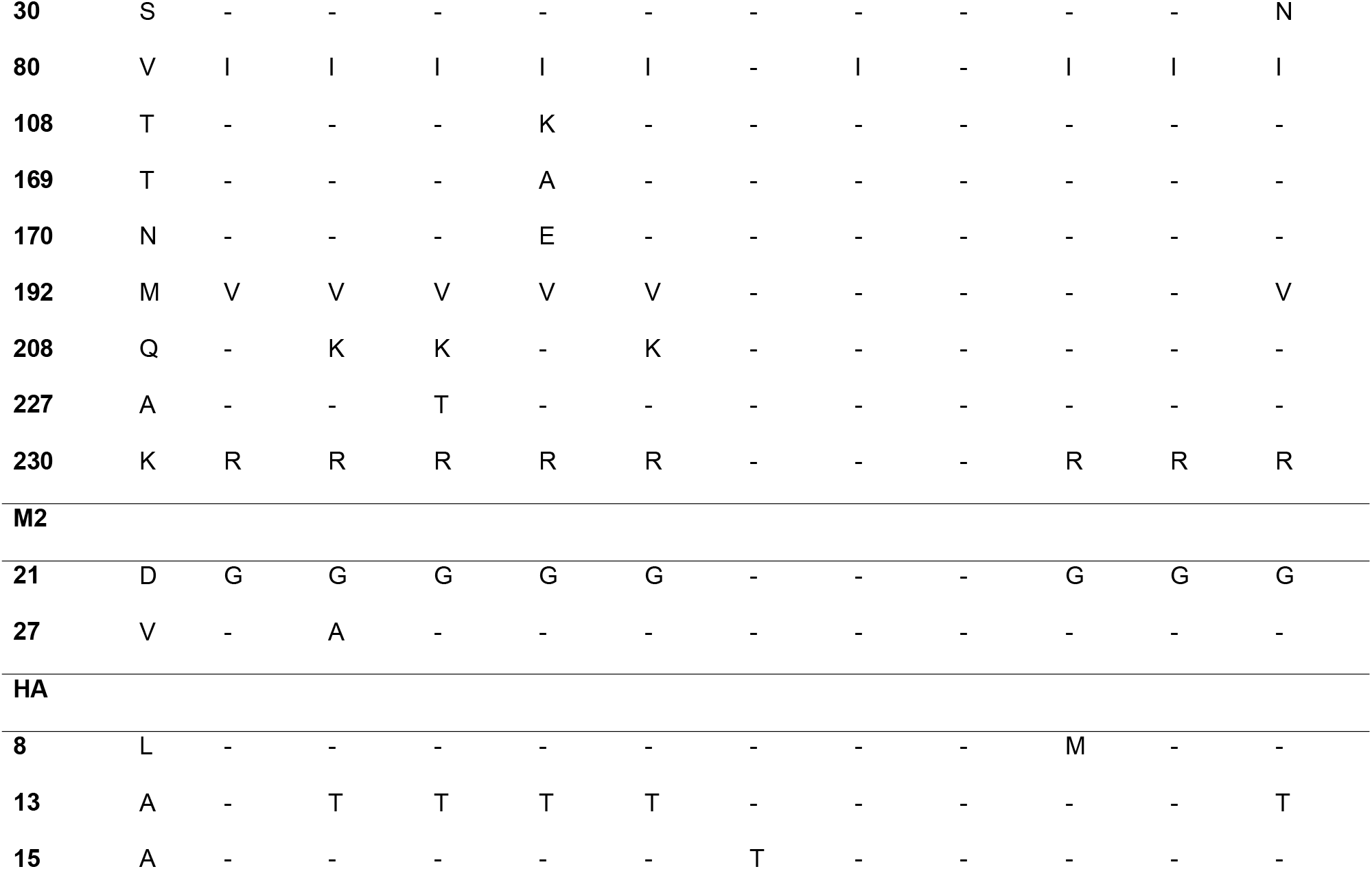
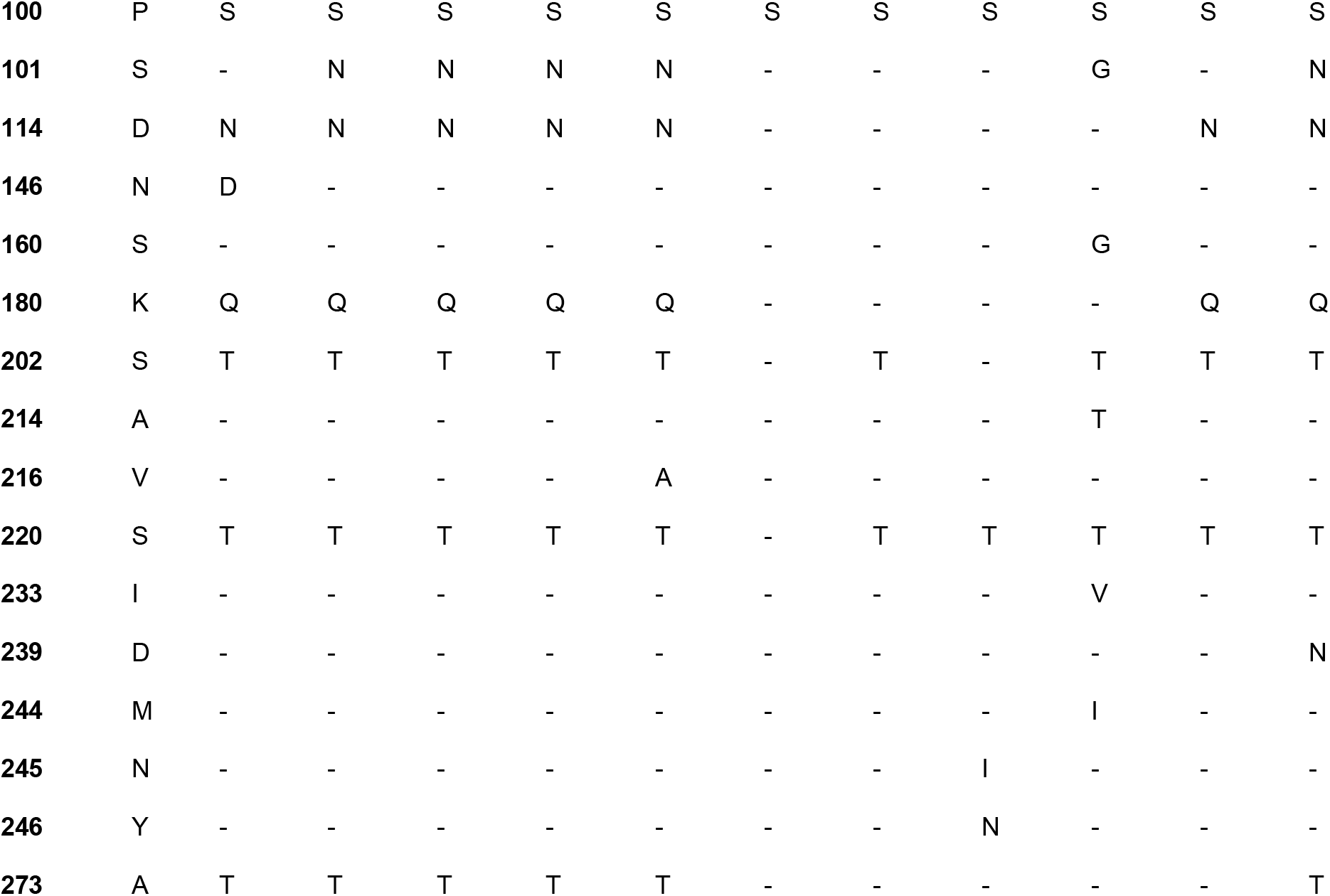
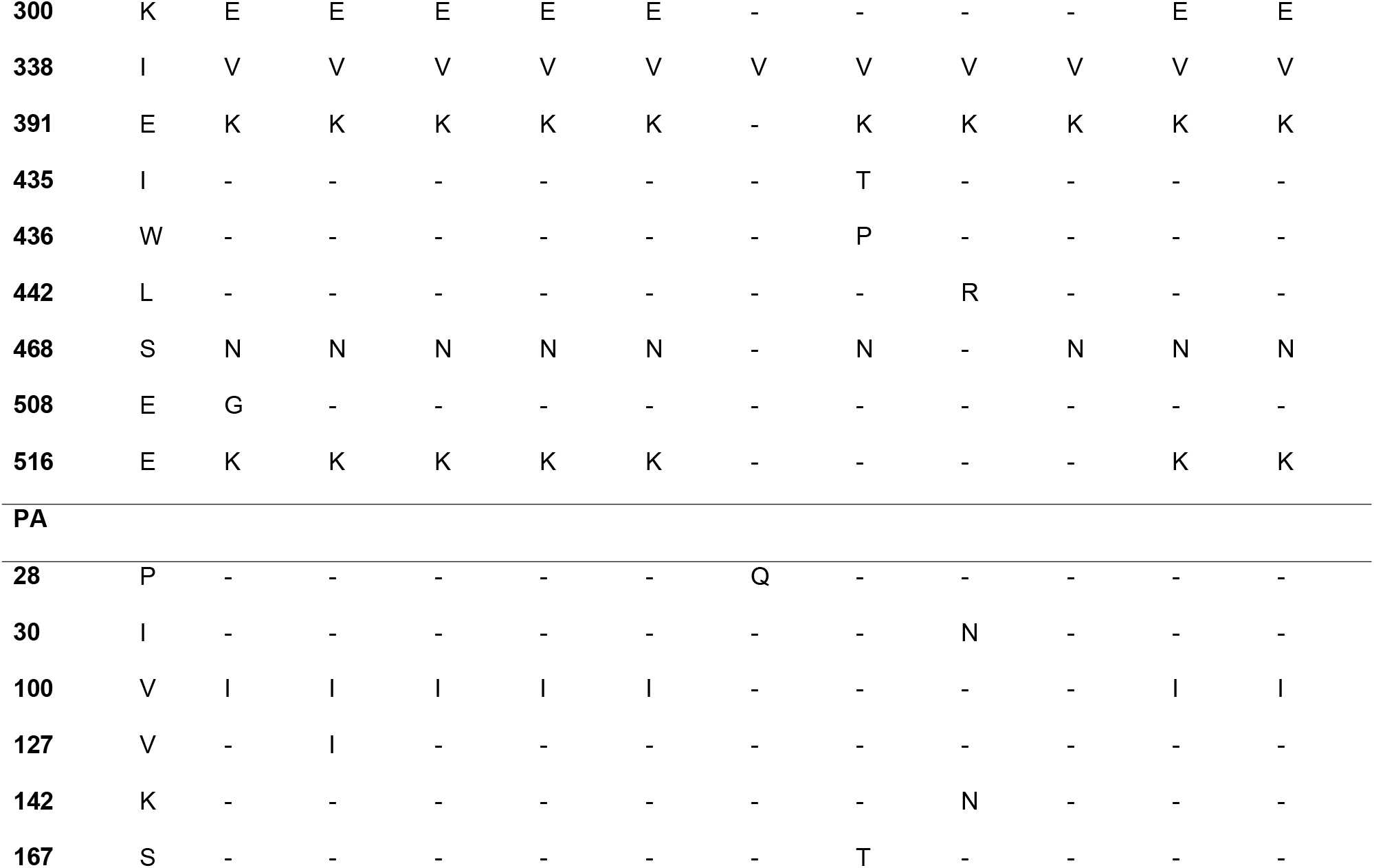
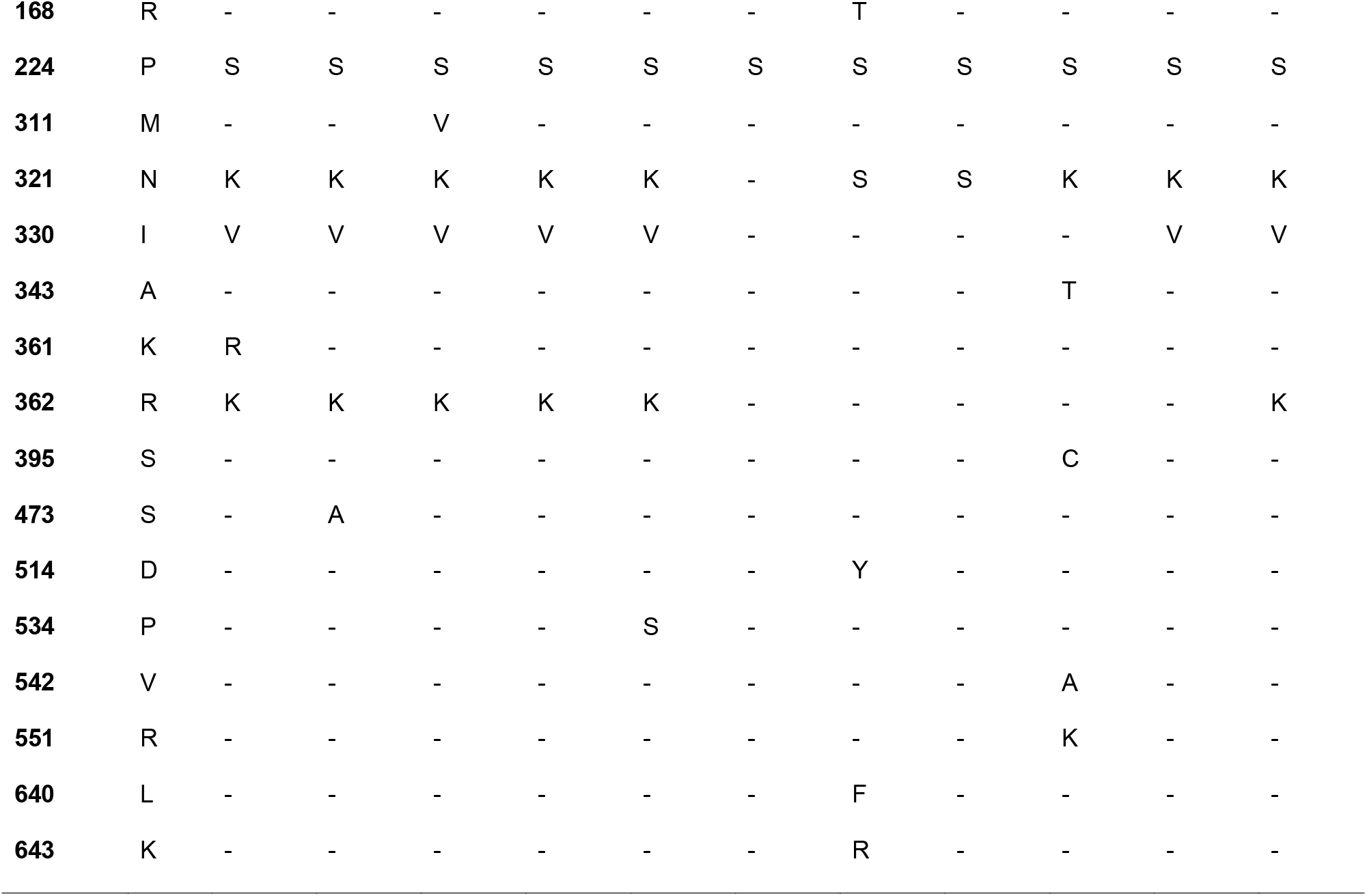
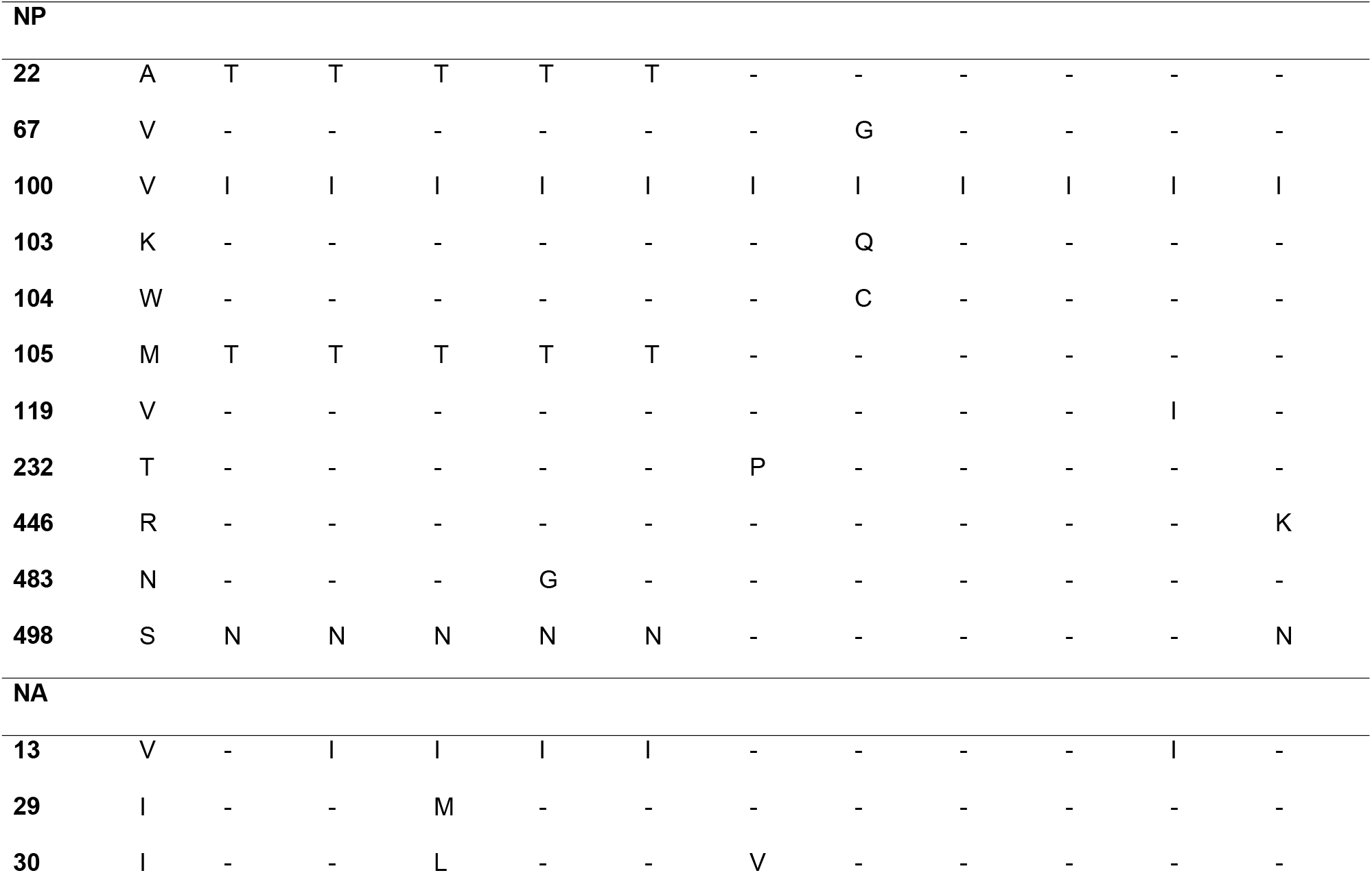
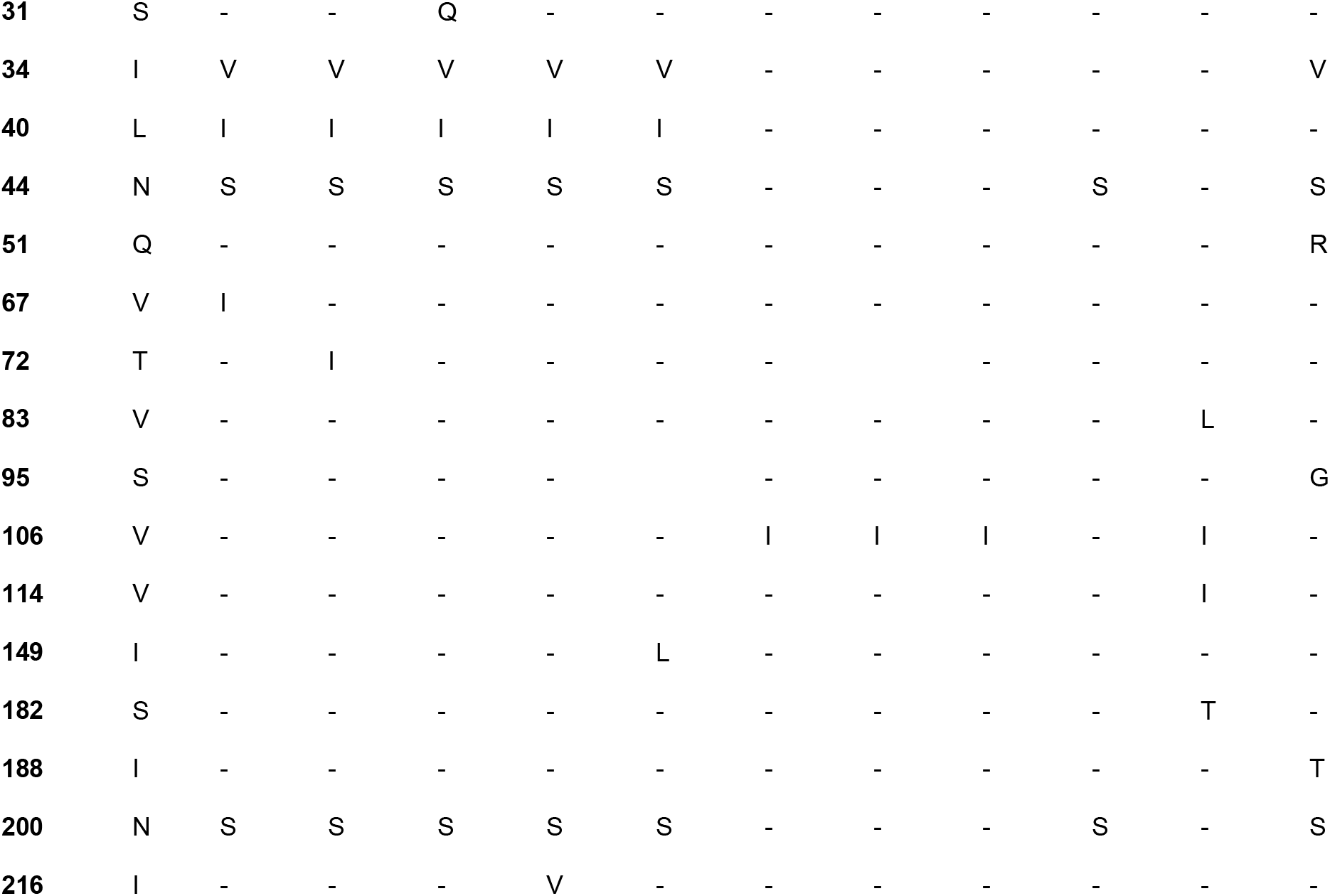
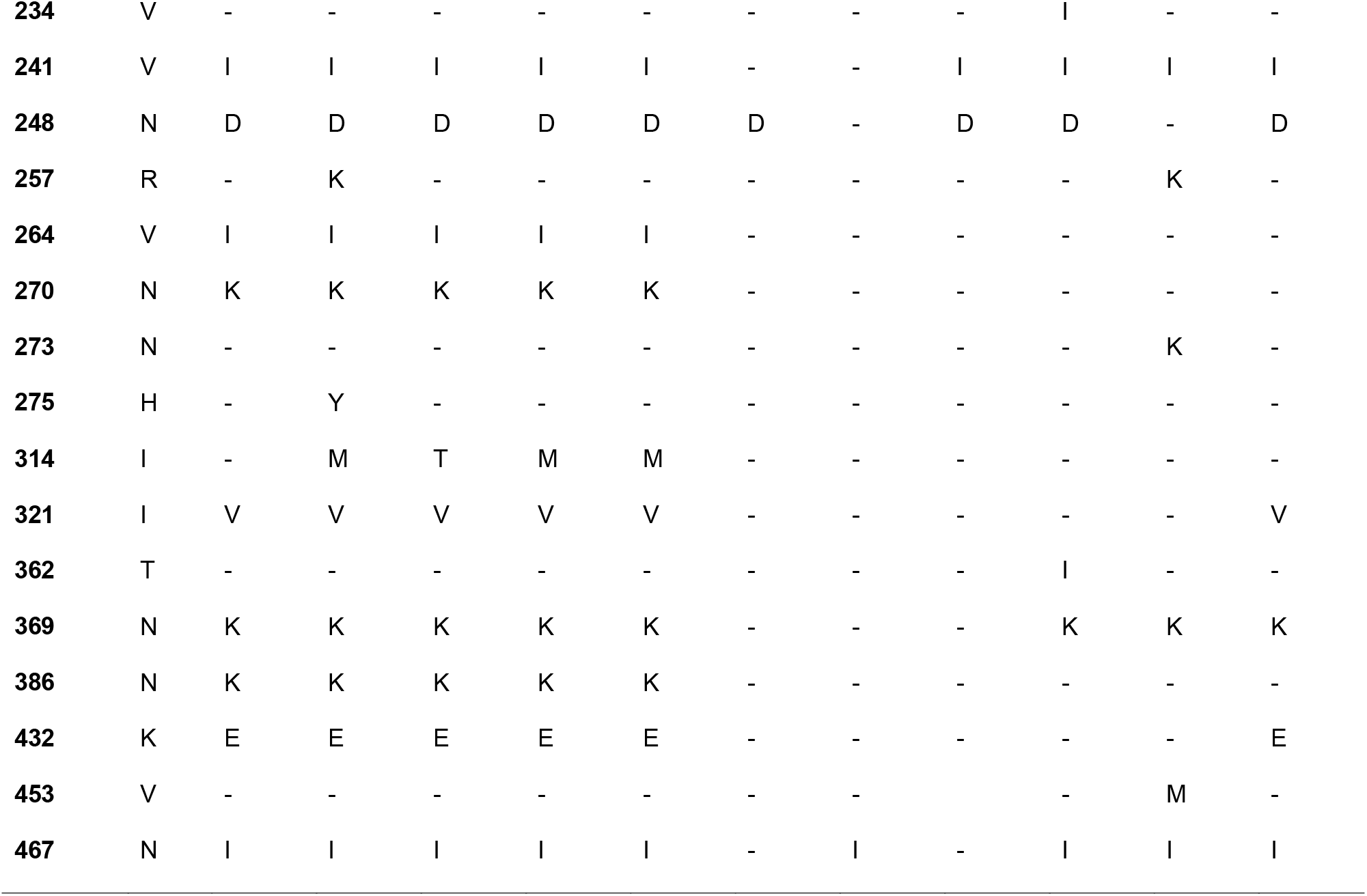
Amino acid substitutions among the five Indian H1N1pdm virus (sequenced in this study) compared to prototype H1N1pdm strain (A/California/07/2009) and other Indian viruses (2009-2014)

| Position | A/ California/ 07/2009 | A/India/DRDE_ GWL812/2015 | A/India/DRDE_ GWL84/2015 | A/India/DRDE_ GWL672/2015 | A/India/DRDE_ GWL897/2015 | A/India/DRDE_ GWL989/2015 | A/Hyd_ NIV51/2009 | A/India/GWL_ DSC/2010 | A/India/Gwl_ 01/2011 | A/India/Nsk12388/2012 | A/India/P132194/2013 | A/India/Ind_ 6427/2014 |
| --- | --- | --- | --- | --- | --- | --- | --- | --- | --- | --- | --- | --- |
| **PB2** |  |  |  |  |  |  |  |  |  |  |  |  |
| **54** | R | K | K | K | K | K | - | K | K | - | - | K |
| **66** | M | I | I | I | I | I | - | - | - | - | - | I |
| **104** | P | - | - | - | - | - | - | - | - | - | L | - |
| **125** | L | - | - | - | - | - | - | F | - | - | - | - |
| **183** | L | - | - | - | - | - | - | - | M | - | - | - |
| **195** | D | N | N | N | N | N | - | - | - | N | - | N |
| **216** | R | - | - | - | - | - | - | - | G | - | - | - |
| **241** | E | - | - | - | - | - | - | G | - | - | - | - |
| **278** | A | - | - | - | - | - | - | V | V | - | - | - |
| **288** | Q | - | - | - | - | - | - | P | - | - | - | - |
| **293** | R | K | K | K | K | K | - | - | - | K | - | K |
| **299** | R | - | K | K | K | K | - | - | - | - | K | - |
| **340** | K | - | - | - | - | - | - | T | T | - | - | - |
| **344** | V | M | M | M | M | M | - | - | - | M | M | M |
| **354** | I | L | L | L | L | L | - | - | - | L | L | L |
| **439** | Q | - | - | - | - | - | - | H | - | - | - | - |
| **441** | D | - | - | - | - | - | - | H | - | - | - | - |
| **453** | S | P | T | T | T | T | - | - | - | - | - | - |
| **456** | N | - | - | - | - | - | - | - | - | - | S | - |
| **588** | T | - | - | - | - | - | - | - | I | - | - | - |
| **731** | V | I | I | I | I | I | - | - | - | I | - | I |
| **758** | I | - | - | - | - | - | - | L | - | - | - | - |
| **PB1** |  |  |  |  |  |  |  |  |  |  |  |  |
| **154** | G | D | D | D | D | D | - | - | - | D | - | D |
| **213** | N | S | - | - | - | - | - | - | - | - | - | - |
| **226** | T | - | - | - | - | - | - | S | - | - | - | - |
| **229** | K | - | - | - | - | - | - | I | - | - | - | - |
| **236** | L | - | - | - | - | - | - | V | - | - | - | - |
| **254** | F | - | - | - | - | - | L | - | - | - | - | - |
| **323** | T | - | - | - | - | - | - | K | - | - | - | - |
| **397** | I | M | M | M | M | M | - | - | M | M | - | M |
| **633** | S | - | - | - | - | - | - | - | - | T | - | - |
| **652** | A | - | - | - | - | - | - | - | - | - | V | - |
| **667** | I | - | - | - | - | - | - | - | - | S | - | - |
| **679** | K | - | - | - | - | - | - | - | - | T | - | - |
| **NS1** |  |  |  |  |  |  |  |  |  |  |  |  |
| **55** | E | K | K | K | K | K | - | - | - | - | - | K |
| **71** | E | - | - | - | - | - | - | K | - | - | - | - |
| **86** | T | - | - | - | I | - | - | - | - | - | - | - |
| **90** | L | I | I | I | I | I | - | - | - | I | - | I |
| **123** | I | V | V | V | V | V | - | V | V | V | V | V |
| **129** | V | - | - | - | - | - | - | - | - | - | I | - |
| **131** | K | E | E | E | E | E | - | - | - | - | - | E |
| **205** | N | S | S | S | S | S | - | - | - | S | S | S |
| **206** | C | - | - | - | - | - | - | - | - | Y | - | - |
| **NS2** |  |  |  |  |  |  |  |  |  |  |  |  |
| **29** | N | S | S | S | S | S | - | - | - | - | - | S |
| **48** | T | A | A | A | A | A | - | - | - | A | A | A |
| **49** | V | - | - | - | I | - | - | - | - | M | - | - |
| **83** | M | - | I | I | I | I | - | - | - | - | - | - |
| **115** | A | - | - | - | - | - | T | - | - | - | - | - |
| **M1** |  |  |  |  |  |  |  |  |  |  |  |  |
| **28** | L | - | - | - | - | - | - | M | - | - | - | - |
| **30** | S | - | - | - | - | - | - | - | - | - | - | N |
| **80** | V | I | I | I | I | I | - | I | - | I | I | I |
| **108** | T | - | - | - | K | - | - | - | - | - | - | - |
| **169** | T | - | - | - | A | - | - | - | - | - | - | - |
| **170** | N | - | - | - | E | - | - | - | - | - | - | - |
| **192** | M | V | V | V | V | V | - | - | - | - | - | V |
| **208** | Q | - | K | K | - | K | - | - | - | - | - | - |
| **227** | A | - | - | T | - | - | - | - | - | - | - | - |
| **230** | K | R | R | R | R | R | - | - | - | R | R | R |
| M2 |  |  |  |  |  |  |  |  |  |  |  |  |
| **21** | D | G | G | G | G | G | - | - | - | G | G | G |
| **27** | V | - | A | - | - | - | - | - | - | - | - | - |
| HA |  |  |  |  |  |  |  |  |  |  |  |  |
| **8** | L | - | - | - | - | - | - | - | - | M | - | - |
| **13** | A | - | T | T | T | T | - | - | - | - | - | T |
| **15** | A | - | - | - | - | - | T | - | - | - | - | - |
| **100** | P | S | S | S | S | S | S | S | S | S | S | S |
| **101** | S | - | N | N | N | N | - | - | - | G | - | N |
| **114** | D | N | N | N | N | N | - | - | - | - | N | N |
| **146** | N | D | - | - | - | - | - | - | - | - | - | - |
| **160** | S | - | - | - | - | - | - | - | - | G | - | - |
| **180** | K | Q | Q | Q | Q | Q | - | - | - | - | Q | Q |
| **202** | S | T | T | T | T | T | - | T | - | T | T | T |
| **214** | A | - | - | - | - | - | - | - | - | T | - | - |
| **216** | V | - | - | - | - | A | - | - | - | - | - | - |
| **220** | S | T | T | T | T | T | - | T | T | T | T | T |
| **233** | I | - | - | - | - | - | - | - | - | V | - | - |
| **239** | D | - | - | - | - | - | - | - | - | - | - | N |
| **244** | M | - | - | - | - | - | - | - | - | I | - | - |
| **245** | N | - | - | - | - | - | - | - | I | - | - | - |
| **246** | Y | - | - | - | - | - | - | - | N | - | - | - |
| **273** | A | T | T | T | T | T | - | - | - | - | - | T |
| **300** | K | E | E | E | E | E | - | - | - | - | E | E |
| **338** | I | V | V | V | V | V | V | V | V | V | V | V |
| **391** | E | K | K | K | K | K | - | K | K | K | K | K |
| **435** | I | - | - | - | - | - | - | T | - | - | - | - |
| **436** | W | - | - | - | - | - | - | P | - | - | - | - |
| **442** | L | - | - | - | - | - | - | - | R | - | - | - |
| **468** | S | N | N | N | N | N | - | N | - | N | N | N |
| **508** | E | G | - | - | - | - | - | - | - | - | - | - |
| **516** | E | K | K | K | K | K | - | - | - | - | K | K |
| **PA** |  |  |  |  |  |  |  |  |  |  |  |  |
| **28** | P | - | - | - | - | - | Q | - | - | - | - | - |
| **30** | I | - | - | - | - | - | - | - | N | - | - | - |
| **100** | V | I | I | I | I | I | - | - | - | - | I | I |
| **127** | V | - | I | - | - | - | - | - | - | - | - | - |
| **142** | K | - | - | - | - | - | - | - | N | - | - | - |
| **167** | S | - | - | - | - | - | - | T | - | - | - | - |
| **168** | R | - | - | - | - | - | - | T | - | - | - | - |
| **224** | P | S | S | S | S | S | S | S | S | S | S | S |
| **311** | M | - | - | V | - | - | - | - | - | - | - | - |
| **321** | N | K | K | K | K | K | - | S | S | K | K | K |
| **330** | I | V | V | V | V | V | - | - | - | - | V | V |
| **343** | A | - | - | - | - | - | - | - | - | T | - | - |
| **361** | K | R | - | - | - | - | - | - | - | - | - | - |
| **362** | R | K | K | K | K | K | - | - | - | - | - | K |
| **395** | S | - | - | - | - | - | - | - | - | C | - | - |
| **473** | S | - | A | - | - | - | - | - | - | - | - | - |
| **514** | D | - | - | - | - | - | - | Y | - | - | - | - |
| **534** | P | - | - | - | - | S | - | - | - | - | - | - |
| **542** | V | - | - | - | - | - | - | - | - | A | - | - |
| **551** | R | - | - | - | - | - | - | - | - | K | - | - |
| **640** | L | - | - | - | - | - | - | F | - | - | - | - |
| **643** | K | - | - | - | - | - | - | R | - | - | - | - |

| NP |  |  |  |  |  |  |  |  |  |  |  |  |
| --- | --- | --- | --- | --- | --- | --- | --- | --- | --- | --- | --- | --- |
| 22 | A | T | T | T | T | T | - | - | - | - | - | - |
| 67 | V | - | - | - | - | - | - | G | - | - | - | - |
| 100 | V | I | I | I | I | I | I | I | I | I | I | I |
| 103 | K | - | - | - | - | - | - | Q | - | - | - | - |
| 104 | W | - | - | - | - | - | - | C | - | - | - | - |
| 105 | M | T | T | T | T | T | - | - | - | - | - | - |
| 119 | V | - | - | - | - | - | - | - | - | - | I | - |
| 232 | T | - | - | - | - | - | P | - | - | - | - | - |
| 446 | R | - | - | - | - | - | - | - | - | - | - | K |
| 483 | N | - | - | - | G | - | - | - | - | - | - | - |
| 498 | S | N | N | N | N | N | - | - | - | - | - | N |
| NA |  |  |  |  |  |  |  |  |  |  |  |  |
| 13 | V | - | I | I | I | I | - | - | - | - | I | - |
| 29 | I | - | - | M | - | - | - | - | - | - | - | - |
| 30 | I | - | - | L | - | - | V | - | - | - | - | - |
| **31** | S | - | - | Q | - | - | - | - | - | - | - | - |
| **34** | I | V | V | V | V | V | - | - | - | - | - | V |
| **40** | L | I | I | I | I | I | - | - | - | - | - | - |
| **44** | N | S | S | S | S | S | - | - | - | S | - | S |
| **51** | Q | - | - | - | - | - | - | - | - | - | - | R |
| **67** | V | I | - | - | - | - | - | - | - | - | - | - |
| **72** | T | - | I | - | - | - | - | - | - | - | - | - |
| **83** | V | - | - | - | - | - | - | - | - | - | L | - |
| **95** | S | - | - | - | - | - | - | - | - | - | - | G |
| **106** | V | - | - | - | - | - | I | I | I | - | I | - |
| **114** | V | - | - | - | - | - | - | - | - | - | I | - |
| **149** | I | - | - | - | - | L | - | - | - | - | - | - |
| **182** | S | - | - | - | - | - | - | - | - | - | T | - |
| **188** | I | - | - | - | - | - | - | - | - | - | - | T |
| **200** | N | S | S | S | S | S | - | - | - | S | - | S |
| **216** | I | - | - | - | V | - | - | - | - | - | - | - |
| **234** | V | - | - | - | - | - | - | - | - | I | - | - |
| **241** | V | I | I | I | I | I | - | - | I | I | I | I |
| **248** | N | D | D | D | D | D | D | - | D | D | - | D |
| **257** | R | - | K | - | - | - | - | - | - | - | K | - |
| **264** | V | I | I | I | I | I | - | - | - | - | - | - |
| **270** | N | K | K | K | K | K | - | - | - | - | - | - |
| **273** | N | - | - | - | - | - | - | - | - | - | K | - |
| **275** | H | - | Y | - | - | - | - | - | - | - | - | - |
| **314** | I | - | M | T | M | M | - | - | - | - | - | - |
| **321** | I | V | V | V | V | V | - | - | - | - | - | V |
| **362** | T | - | - | - | - | - | - | - | - | I | - | - |
| **369** | N | K | K | K | K | K | - | - | - | K | K | K |
| **386** | N | K | K | K | K | K | - | - | - | - | - | - |
| **432** | K | E | E | E | E | E | - | - | - | - | - | E |
| **453** | V | - | - | - | - | - | - | - | - | - | M | - |
| **467** | N | I | I | I | I | I | - | I | - | I | I | I |

Next, we performed recombination and selection pressure analysis to understand the evolution of different parts of the genome. Selection pressure analysis showed that a large number of codons were under strong purifying selection. 23 sites were identified to be under positive selection in HA, NA, NP and MP (Table 4), with 7 sites in HA, 6 in NA and NP and 4 in MP. Of these position 239 in HA and positions 13, 248, 275 and 386 in NA were selected by all the different approaches used in this analysis.

**Table 4:**
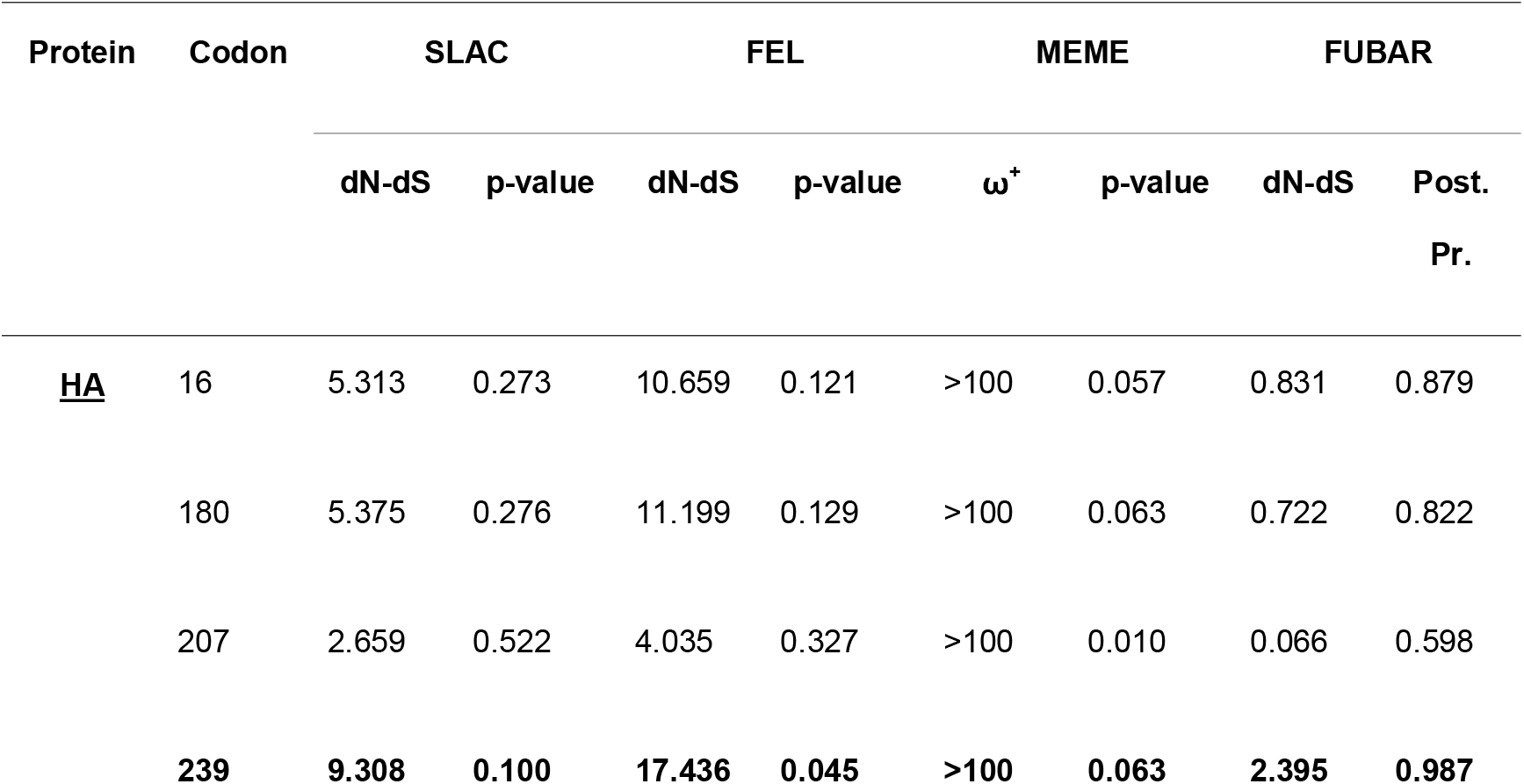
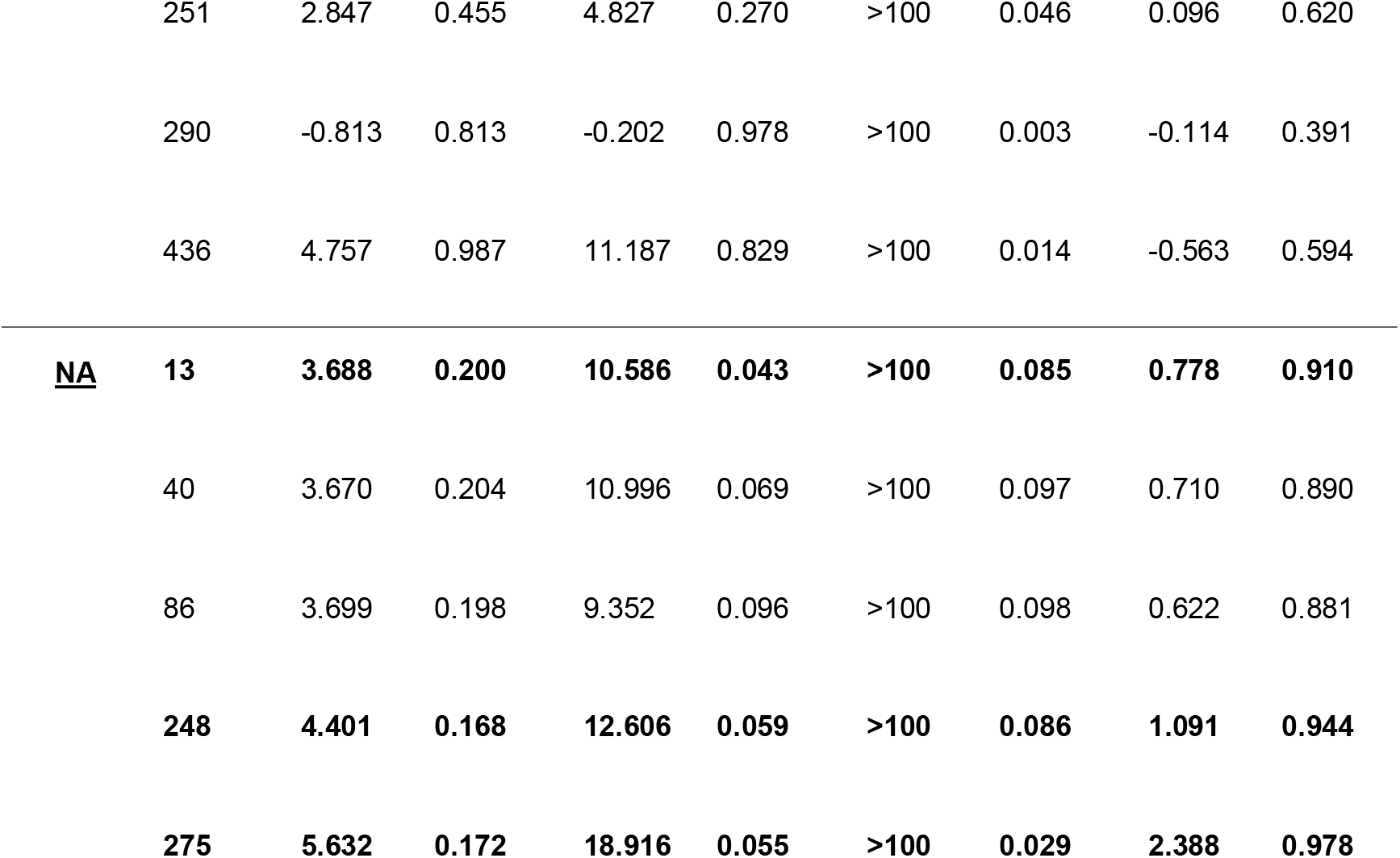
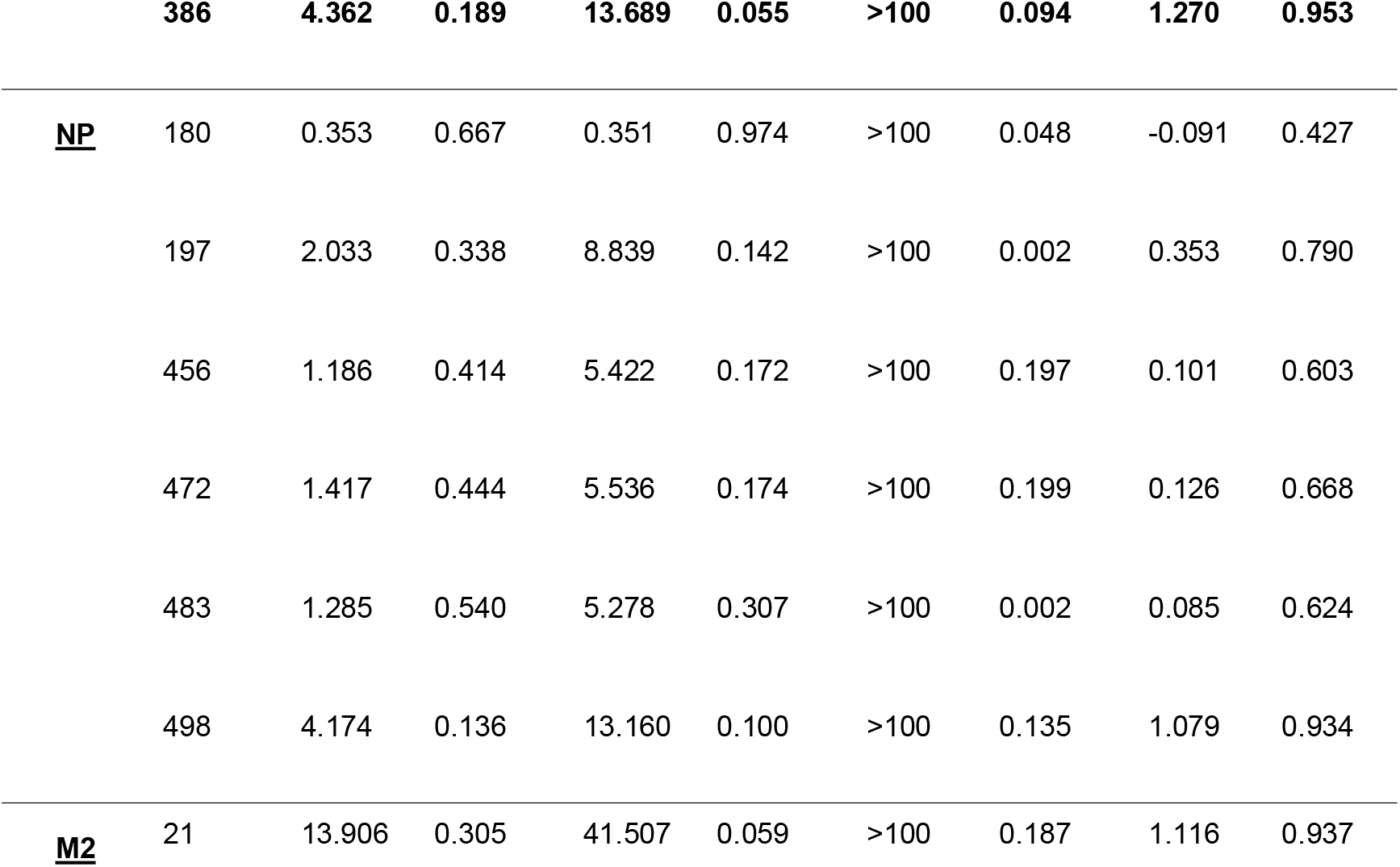
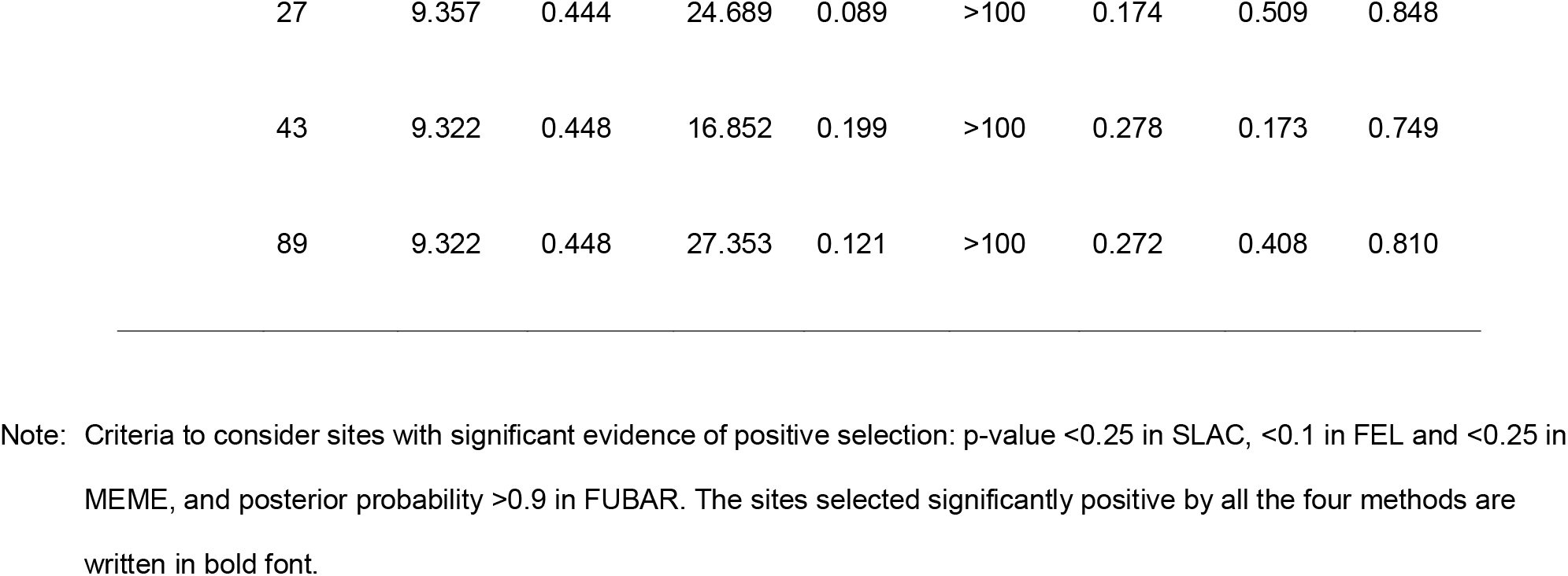
Selection pressure analysis of HA protein (566 codons); NA protein (469 codons), NP Protein (498 codons) and M2 Protein (97 codons) of H1N1pdm virus using SLAC, FEL, MEME and FUBAR methods.

In order to assess the relatedness of the Indian 2015 genomes to circulating Influenza A H1N1 genomes worldwide (2009-2015), we performed phylogenetic analysis of representative sequences of A(H1N1)pdm09 from distinct geographical regions. A maximum likelihood based phylogenetic tree revealed 10 distinct genogroups (1-5, 6A, 6B, 6C, 7 and 8) (Fig 3). While the prototype strain belongs to genogroup 1, the 2015 Indian genomes, grouped into genogroup 6B along with a large number of viruses circulating across the globe since 2013. Older isolates from India grouped into genogroup 1, 3, 4, 7, 6A and 6C. As predicted from the differences in amino acids, A/India/DRDE_GWL812/2015 grouped into a separate cluster (I) within genogroup 6B compared to the other 4 2015 genomes (cluster II). The individual gene segment wise phylogeny also revealed similar topology (results not shown).

**Fig 3.**
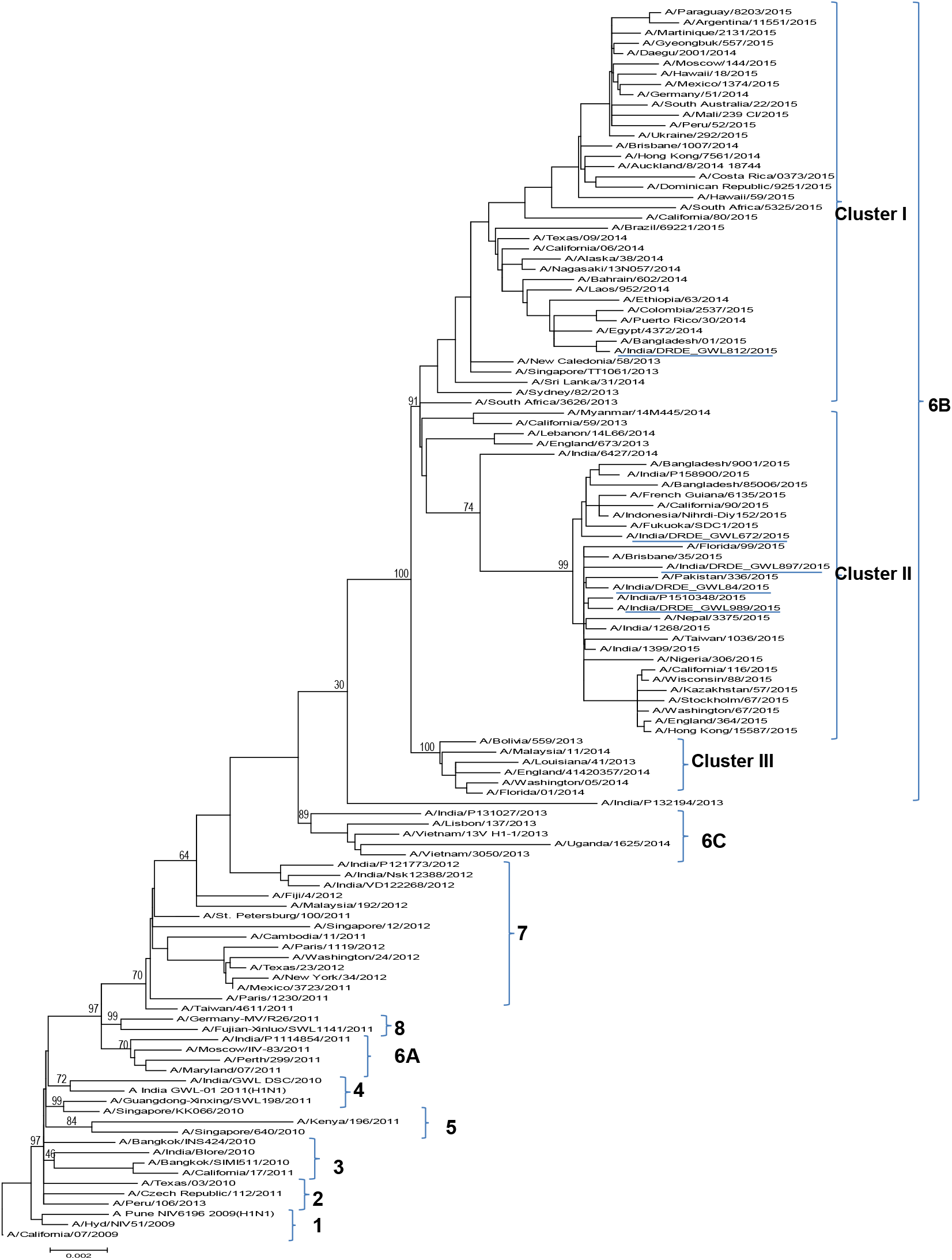
Phylogenetic tree among A(H1N1)pdm09 viruses generated by maximum likelihood method based on the nucleotide sequence of concatenated whole genome of geographically diverse viruses.

The molecular clock analysis revealed the mean rate of nucleotide substitution among all the A(H1N1)pdm09 sequnces analysed in this study is 2.543×10^-3^ substitutions/site/year (95% HPD, 2.181×10^-3^ to 2.266×10^-3^).Using a time to most recent common ancestor (TMRCA) approach, we found that the genogroup 6B emerged around 26 Dec 2011 (,with respect to the most recent isolate of 2015. The maximum clade credibility tree showed multiple clades, all supported by high posterior probabilities (>0.95) (Figure 4). This analysis predicted south east Asia as the ancestral state of genogroup 6B and classified it into 3 distinct clades which diverged at 28 Jan 2012, 01 Apr 2012 and 29 Apr 2012 3 (A/India/DRDE_GWL84/2015, A/India/DRDE_GWL897/2015, and A/India/DRDE_GWL989/2015) out of 5 Indian 2015 genomes clustered into clade 2a, 1 (A/India/DRDE_GWL672/2015) in clade 2b and A/India/DRDE_GWL812/2015 grouped into Clade 3 together with an isolate from Bangladesh (A/Bangladesh/01/2015) and a large number of global isolates.

**Fig 4.**
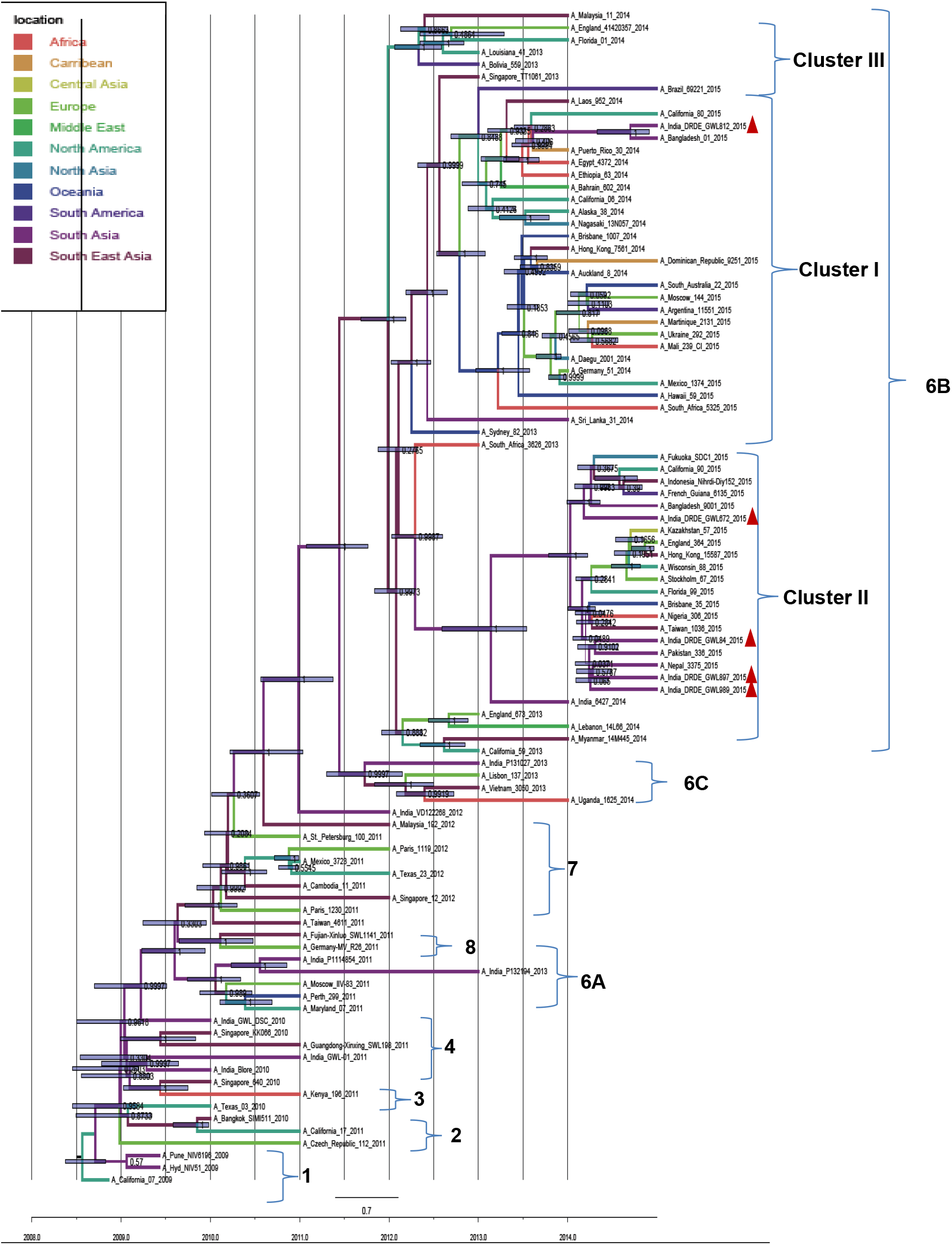
Maximum clade credibility (MCC) tree of A(H1N1)pdm09 viruses constructed using BEAST version 1.7.4. Colors of branches indicate geographic locations as per the color key. Branch lengths correspond to lengths of time. The letters at the nodes correspond to posterior probability values and divergence date.

Phylogeographic analysis of the genogroup 6B viruses of A(H1N1)pdm09 using SPREAD showed the spatial diffusion of these viruses (Figure 5). The geograpahical diffusion pattern revealed 39 well supported transitions of viruses (BF>3) under the Bayesian stochastic search variable selection (BSSVS) model. In particular, migration events for the virus were noted between Africa – Europe (BF=36555), Central Asia – middle East (BF=1635), south America – south Asia (BF=1466) with highest credibility. The migration between Africa-North America and Caribbean-north America Europe-south Asia, central Asia-south America were also characterized by high BF values (>250) (Table S2).

**Fig 5.**
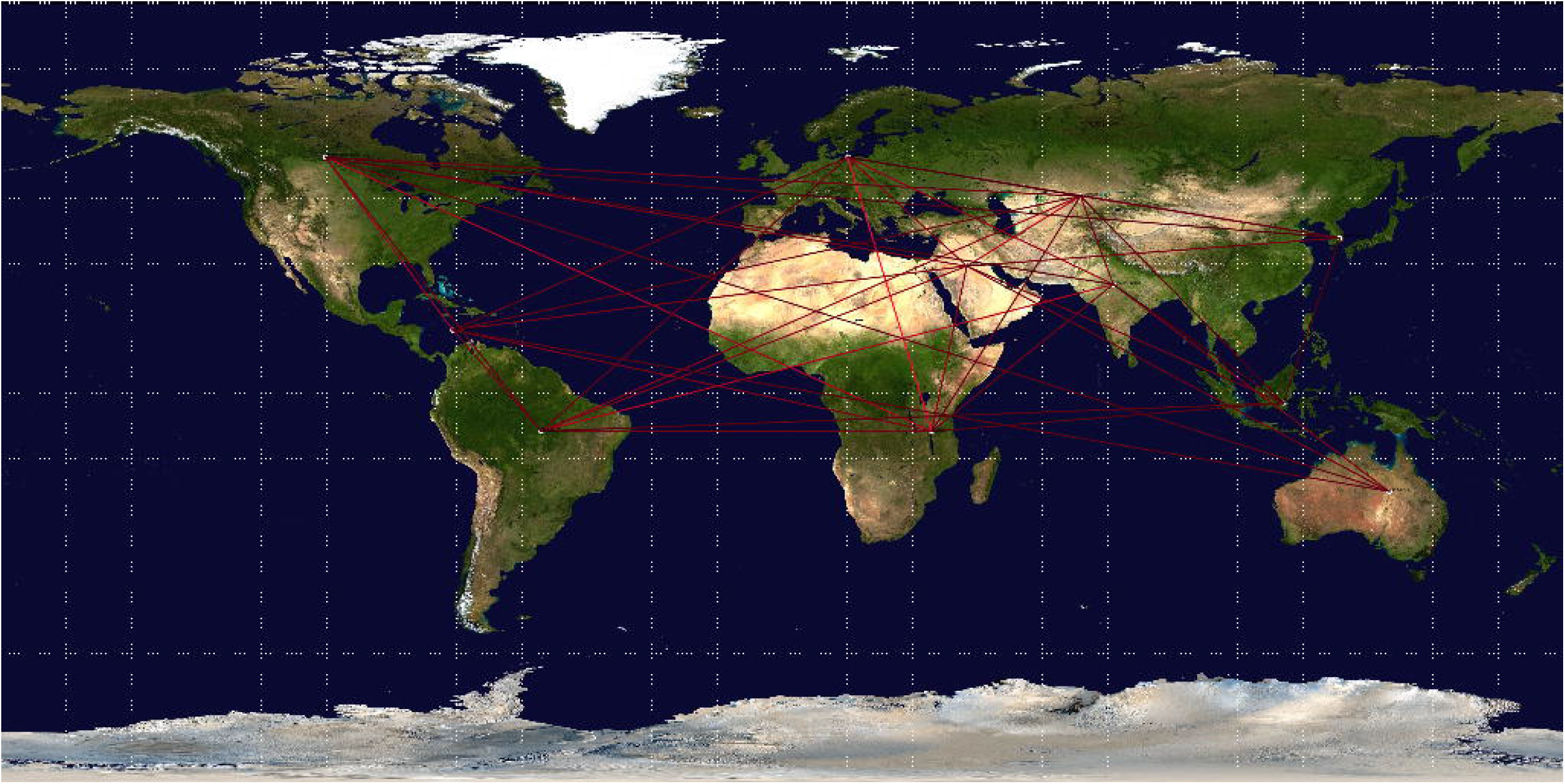
Map showing the global dispersal of cosmopolitan genotype of A(H1N1)pdm09 viruses, belonging to genogroup 6B. The migration route is inferred from branching pattern of maximum clade credibility (MCC) tree, as prepared using SPREAD software. The migration route supported by BF values > 3 is considered significant and is shown in this figure.

## 4. Discussion

Influenza virus A(H1N1)pdm09, which causes swine flu in human, is a rapidly changing virus requiring a seasonal review of circulating strains and evaluation of vaccine effectiveness. The emergence of the quadruple assortment strain in 2009 and the resulting pandemic attest to the serious consequences of emergence of new strains of H1N1. In 2015, an unprecedented number of Influenza cases with increased severity were reported in India. In this study we obtained full length influenza virus sequences from 5 individuals during this outbreak. Analysis of the Hemaglutinin (HA) gene alone revealed 11 substitutions common to all 5 strains when compared to the prototype vaccine strain (A/California/07/2009), 3 of which were in the antigenic site. This is a matter of great concern as the probability of occurrence of a new flu epidemic has been linked either to appearance of >10 amino acid substitutions in the HA protein or > 4 amino acid substitutions in its antigenic sites (Park et al., 2009). Further, HA: K180Q in the Sa antigenic site was also found to be under strong positive selection. 2 of the substitutions that we have identified in the HA gene (HA1: S220T; HA2 E391K) have been reported to increase transmission of the virus.

Surprisingly, the Neuraminidase (NA) gene was found to be most divergent both from analysis of nucleotides and translated amino acids and also harboured the maximum number of sites under positive selection. The presence of the characteristic Histidine to Tyrosine change in the drug binding pocket-NA: H275Y in A/India/DRDE_GWL84/2015 confirms the circulation of oseltamivir resistant virus in this outbreak. We also identified two novel mutations T72I and R257K in NA protein of this virus which needs further investigation. Additionally, we found the N248D substitution in the drug binding pocket, linked with reduction in sensitivity to Oseltamivir in all 5 strains. In addition to this, 3 substitutions (V241I, N369K, and N386K) identified in all the five Indian viruses were also consistently recorded in oseltamivir - resistant viruses isolated from China and Japan during 2013-14. These mutations were linked to offset the destabilizing effect of H275Y and enhance the virus transmission (Escuret et al., 2008, Huang et al., 2015; Koel et al., 2013; Koel et al., 2015). The NA: N386K in particular, leading to the loss of a glycosylation site, was found to be under strong selection pressure.

Codon specific selection pressures revealed strong purifying selection, as observed in other influenza viruses. The NA: V13I (in 4 of the 2015 strains) and NP: S498N (observed in Indian isolates of 2014 and 2015) were found to be under positive and strong positive selection respectively.

Molecular phylogeny analysis of the concatenated genome divides the strains from the H1N1 pandemic 2009 strains into 10 genogroups. Of these groups, 6B is the most diverse and contains within it most of the viruses isolated since 2013 including the 5 2015 Indian strains. Our analysis estimates that the common ancestor for genogroup 6B arose in end of 2011 with south east Asia as the ancestral state. Consistent with previous results we also dated the emergence of the pandemic 2009 strain to mid 2008. (Fraser et al., 2009, Smith et al., 2009).

Unlike many other pathogens which cluster geographically, influenza viruses generally follow temporal clustering pattern. Maximum clade credibility (MCC) phylogenetic analysis demonstrates the existence of multiple clusters among genogroup 6B (clusters are confirmed on the basis of more than 90% posterior probability values). The existence of multiple clusters confirms the idea that the virus is evolving continuously, raising the possibility of pandemic strain emergence.

This study is the first extensive complete genome analysis of 2015 Indian A(H1N1)pdm09 viruses. The identification of mutant A(H1N1)pdm09 viruses, particularly the presence of so many mutations in HA and as well as in NA are quite important both from surveillance, prediction of outbreak and therapeutic intervention.

## Supporting information

Details of sequences retrieved from the EpiFlu Database for phylogenetic analysis in this study

Spacial diffusion of A(H1N1)pdm09 viruses between geographical territories with bayes factor

Bayesian test statistics

## Acknowledgements

This work was funded by the DRDO. The authors are thankful to Dr D.K. Dubey, Director, Defence Research and Development Establishment, Ministry of Defence, Government of India, for his support, constant inspiration and providing the necessary facilities for this study. The authors are also thankful to the Commissioner, Health Services, Govt of MP, IDSP network and CMHO, respective districts of MP for providing the clinical samples. The authors are thankful to GISAID and all the contributors of A(H1N1)pdm09 sequences, which were utilised in this study.

