## Supplementary material for "Next-Gen sequencing of novel pandemic swine flu [A(H1N1)pdm09] virus in India revealed novel mutations across the genome": Details of sequences retrieved from the EpiFlu Database for phylogenetic analysis in this study

**Table S1:	Details of the A(H1N1)pdm09 sequences retrieved from the Global Initiative on Sharing Avian Influenza Data	(GISAID)’s EpiFlu Database for phylogenetic analysis in this study**

We acknowledge the authors, originating and submitting laboratories of the sequences from GISAID’s EpiFlu™ Database on which this research is based. The list is detailed below.

All submitters of data may be contacted directly via the GISAID website www.gisaid.org

| **ID** | **Segment** | **Country** | **Collection date** | **Isolate name** | **Originating laboratory** | **Submitting laboratory** | **Authors** |
| --- | --- | --- | --- | --- | --- | --- | --- |
| EPI695457 | PA | Brazil | 2015-Sep-04 | A/Brazil/69221/2015 | Instituto Adolfo Lutz | Centers for Disease Control and Prevention | NA |
| EPI695458 | PB2 | Brazil | 2015-Sep-04 | A/Brazil/69221/2015 | Instituto Adolfo Lutz | Centers for Disease Control and Prevention | NA |
| EPI695459 | PB1 | Brazil | 2015-Sep-04 | A/Brazil/69221/2015 | Instituto Adolfo Lutz | Centers for Disease Control and Prevention | NA |
| EPI695454 | NP | Brazil | 2015-Sep-04 | A/Brazil/69221/2015 | Instituto Adolfo Lutz | Centers for Disease Control and Prevention | NA |
| EPI695455 | NS | Brazil | 2015-Sep-04 | A/Brazil/69221/2015 | Instituto Adolfo Lutz | Centers for Disease Control and Prevention | NA |
| EPI695456 | MP | Brazil | 2015-Sep-04 | A/Brazil/69221/2015 | Instituto Adolfo Lutz | Centers for Disease Control and Prevention | NA |
| EPI695460 | NA | Brazil | 2015-Sep-04 | A/Brazil/69221/2015 | Instituto Adolfo Lutz | Centers for Disease Control and Prevention | NA |
| EPI695461 | HA | Brazil | 2015-Sep-04 | A/Brazil/69221/2015 | Instituto Adolfo Lutz | Centers for Disease Control and Prevention | NA |
| EPI680459 | PB2 | Pakistan | 2015-Nov-14 | A/Pakistan/336/2015 | National Institute of Health | Centers for Disease Control and Prevention | NA |
| EPI680460 | PB1 | Pakistan | 2015-Nov-14 | A/Pakistan/336/2015 | National Institute of Health | Centers for Disease Control and Prevention | NA |
| EPI680455 | NP | Pakistan | 2015-Nov-14 | A/Pakistan/336/2015 | National Institute of Health | Centers for Disease Control and Prevention | NA |
| EPI680456 | NS | Pakistan | 2015-Nov-14 | A/Pakistan/336/2015 | National Institute of Health | Centers for Disease Control and Prevention | NA |
| EPI680457 | MP | Pakistan | 2015-Nov-14 | A/Pakistan/336/2015 | National Institute of Health | Centers for Disease Control and Prevention | NA |
| EPI680458 | PA | Pakistan | 2015-Nov-14 | A/Pakistan/336/2015 | National Institute of Health | Centers for Disease Control and Prevention | NA |
| EPI680461 | NA | Pakistan | 2015-Nov-14 | A/Pakistan/336/2015 | National Institute of Health | Centers for Disease Control and Prevention | NA |
| EPI680462 | HA | Pakistan | 2015-Nov-14 | A/Pakistan/336/2015 | National Institute of Health | Centers for Disease Control and Prevention | NA |
| EPI691359 | NP | Taiwan | 2015-Sep-05 | A/Taiwan/1036/2015 | ADImmune Corporation | Centers for Disease Control and Prevention | NA |
| EPI691360 | NS | Taiwan | 2015-Sep-05 | A/Taiwan/1036/2015 | ADImmune Corporation | Centers for Disease Control and Prevention | NA |
| EPI691361 | MP | Taiwan | 2015-Sep-05 | A/Taiwan/1036/2015 | ADImmune Corporation | Centers for Disease Control and Prevention | NA |
| EPI691362 | PA | Taiwan | 2015-Sep-05 | A/Taiwan/1036/2015 | ADImmune Corporation | Centers for Disease Control and Prevention | NA |
| EPI691363 | PB2 | Taiwan | 2015-Sep-05 | A/Taiwan/1036/2015 | ADImmune Corporation | Centers for Disease Control and Prevention | NA |
| EPI691364 | PB1 | Taiwan | 2015-Sep-05 | A/Taiwan/1036/2015 | ADImmune Corporation | Centers for Disease Control and Prevention | NA |
| EPI691365 | NA | Taiwan | 2015-Sep-05 | A/Taiwan/1036/2015 | ADImmune Corporation | Centers for Disease Control and Prevention | NA |
| EPI691366 | HA | Taiwan | 2015-Sep-05 | A/Taiwan/1036/2015 | ADImmune Corporation | Centers for Disease Control and Prevention | NA |
| EPI672206 | PB2 | United States | 2015-Sep-12 | A/California/90/2015 | California Department of Health Services | Centers for Disease Control and Prevention | NA |
| EPI672207 | PB1 | United States | 2015-Sep-12 | A/California/90/2015 | California Department of Health Services | Centers for Disease Control and Prevention | NA |
| EPI672202 | NP | United States | 2015-Sep-12 | A/California/90/2015 | California Department of Health Services | Centers for Disease Control and Prevention | NA |
| EPI672203 | NS | United States | 2015-Sep-12 | A/California/90/2015 | California Department of Health Services | Centers for Disease Control and Prevention | NA |
| EPI672204 | MP | United States | 2015-Sep-12 | A/California/90/2015 | California Department of Health Services | Centers for Disease Control and Prevention | NA |
| EPI672205 | PA | United States | 2015-Sep-12 | A/California/90/2015 | California Department of Health Services | Centers for Disease Control and Prevention | NA |
| EPI672208 | NA | United States | 2015-Sep-12 | A/California/90/2015 | California Department of Health Services | Centers for Disease Control and Prevention | NA |
| EPI672209 | HA | United States | 2015-Sep-12 | A/California/90/2015 | California Department of Health Services | Centers for Disease Control and Prevention | NA |
| EPI691214 | NP | United States | 2015-Dec-19 | A/Florida/99/2015 | Florida Department of Health-Tampa | Centers for Disease Control and Prevention | NA |
| EPI691215 | NS | United States | 2015-Dec-19 | A/Florida/99/2015 | Florida Department of Health-Tampa | Centers for Disease Control and Prevention | NA |
| EPI691216 | MP | United States | 2015-Dec-19 | A/Florida/99/2015 | Florida Department of Health-Tampa | Centers for Disease Control and Prevention | NA |
| EPI691217 | PA | United States | 2015-Dec-19 | A/Florida/99/2015 | Florida Department of Health-Tampa | Centers for Disease Control and Prevention | NA |
| EPI691218 | PB2 | United States | 2015-Dec-19 | A/Florida/99/2015 | Florida Department of Health-Tampa | Centers for Disease Control and Prevention | NA |
| EPI691219 | PB1 | United States | 2015-Dec-19 | A/Florida/99/2015 | Florida Department of Health-Tampa | Centers for Disease Control and Prevention | NA |
| EPI691220 | NA | United States | 2015-Dec-19 | A/Florida/99/2015 | Florida Department of Health-Tampa | Centers for Disease Control and Prevention | NA |
| EPI691221 | HA | United States | 2015-Dec-19 | A/Florida/99/2015 | Florida Department of Health-Tampa | Centers for Disease Control and Prevention | NA |
| EPI693486 | PA | Nepal | 2015-Sep-09 | A/Nepal/3375/2015 | National Public Health Laboratory | Centers for Disease Control and Prevention | NA |
| EPI693483 | NP | Nepal | 2015-Sep-09 | A/Nepal/3375/2015 | National Public Health Laboratory | Centers for Disease Control and Prevention | NA |
| EPI693484 | NS | Nepal | 2015-Sep-09 | A/Nepal/3375/2015 | National Public Health Laboratory | Centers for Disease Control and Prevention | NA |
| EPI693485 | MP | Nepal | 2015-Sep-09 | A/Nepal/3375/2015 | National Public Health Laboratory | Centers for Disease Control and Prevention | NA |
| EPI693487 | PB2 | Nepal | 2015-Sep-09 | A/Nepal/3375/2015 | National Public Health Laboratory | Centers for Disease Control and Prevention | NA |
| EPI693488 | PB1 | Nepal | 2015-Sep-09 | A/Nepal/3375/2015 | National Public Health Laboratory | Centers for Disease Control and Prevention | NA |
| EPI693489 | NA | Nepal | 2015-Sep-09 | A/Nepal/3375/2015 | National Public Health Laboratory | Centers for Disease Control and Prevention | NA |
| EPI693490 | HA | Nepal | 2015-Sep-09 | A/Nepal/3375/2015 | National Public Health Laboratory | Centers for Disease Control and Prevention | NA |
| EPI687126 | NP | Japan | 2015-Mar-16 | A/Fukuoka/SDC1/2015 | Virus Research Center, Sendai Medical Center | National Institute of Infectious Diseases (NIID) | Takashita,Emi; Fujisaki,Seiichiro; Shirakura,Masayuki; Watanabe,Shinji; Odagiri,Takato |
| EPI687127 | NS | Japan | 2015-Mar-16 | A/Fukuoka/SDC1/2015 | Virus Research Center, Sendai Medical Center | National Institute of Infectious Diseases (NIID) | Takashita,Emi; Fujisaki,Seiichiro; Shirakura,Masayuki; Watanabe,Shinji; Odagiri,Takato |
| EPI687128 | MP | Japan | 2015-Mar-16 | A/Fukuoka/SDC1/2015 | Virus Research Center, Sendai Medical Center | National Institute of Infectious Diseases (NIID) | Takashita,Emi; Fujisaki,Seiichiro; Shirakura,Masayuki; Watanabe,Shinji; Odagiri,Takato |
| EPI687129 | PA | Japan | 2015-Mar-16 | A/Fukuoka/SDC1/2015 | Virus Research Center, Sendai Medical Center | National Institute of Infectious Diseases (NIID) | Takashita,Emi; Fujisaki,Seiichiro; Shirakura,Masayuki; Watanabe,Shinji; Odagiri,Takato |
| EPI687130 | PB2 | Japan | 2015-Mar-16 | A/Fukuoka/SDC1/2015 | Virus Research Center, Sendai Medical Center | National Institute of Infectious Diseases (NIID) | Takashita,Emi; Fujisaki,Seiichiro; Shirakura,Masayuki; Watanabe,Shinji; Odagiri,Takato |
| EPI687131 | PB1 | Japan | 2015-Mar-16 | A/Fukuoka/SDC1/2015 | Virus Research Center, Sendai Medical Center | National Institute of Infectious Diseases (NIID) | Takashita,Emi; Fujisaki,Seiichiro; Shirakura,Masayuki; Watanabe,Shinji; Odagiri,Takato |
| EPI586589 | NA | Japan | 2015-Mar-16 | A/Fukuoka/SDC1/2015 | Virus Research Center, Sendai Medical Center | National Institute of Infectious Diseases (NIID) | Takashita,Emi; Fujisaki,Seiichiro; Shirakura,Masayuki; Watanabe,Shinji; Odagiri,Takato |
| EPI586590 | HA | Japan | 2015-Mar-16 | A/Fukuoka/SDC1/2015 | Virus Research Center, Sendai Medical Center | National Institute of Infectious Diseases (NIID) | Takashita,Emi; Fujisaki,Seiichiro; Shirakura,Masayuki; Watanabe,Shinji; Odagiri,Takato |
| EPI584985 | NP | India | 2015-Jan-27 | A/India/1268/2015 | National Institute of Virology | Centers for Disease Control and Prevention | NA |
| EPI584987 | NS | India | 2015-Jan-27 | A/India/1268/2015 | National Institute of Virology | Centers for Disease Control and Prevention | NA |
| EPI584988 | MP | India | 2015-Jan-27 | A/India/1268/2015 | National Institute of Virology | Centers for Disease Control and Prevention | NA |
| EPI584989 | PA | India | 2015-Jan-27 | A/India/1268/2015 | National Institute of Virology | Centers for Disease Control and Prevention | NA |
| EPI584990 | PB2 | India | 2015-Jan-27 | A/India/1268/2015 | National Institute of Virology | Centers for Disease Control and Prevention | NA |
| EPI584991 | PB1 | India | 2015-Jan-27 | A/India/1268/2015 | National Institute of Virology | Centers for Disease Control and Prevention | NA |
| EPI584992 | NA | India | 2015-Jan-27 | A/India/1268/2015 | National Institute of Virology | Centers for Disease Control and Prevention | NA |
| EPI584993 | HA | India | 2015-Jan-27 | A/India/1268/2015 | National Institute of Virology | Centers for Disease Control and Prevention | NA |
| EPI630416 | NA | India | 2015-Mar-30 | A/India/P158900/2015 | National Institute of Virology | National Institute of Infectious Diseases (NIID) | Takashita,Emi; Fujisaki,Seiichiro; Shirakura,Masayuki; Watanabe,Shinji; Odagiri,Takato |
| EPI630417 | HA | India | 2015-Mar-30 | A/India/P158900/2015 | National Institute of Virology | National Institute of Infectious Diseases (NIID) | Takashita,Emi; Fujisaki,Seiichiro; Shirakura,Masayuki; Watanabe,Shinji; Odagiri,Takato |
| EPI635511 | NP | India | 2015-Mar-30 | A/India/P158900/2015 | National Institute of Virology | National Institute of Infectious Diseases (NIID) | Takashita,Emi; Fujisaki,Seiichiro; Shirakura,Masayuki; Watanabe,Shinji; Odagiri,Takato |
| EPI635512 | NS | India | 2015-Mar-30 | A/India/P158900/2015 | National Institute of Virology | National Institute of Infectious Diseases (NIID) | Takashita,Emi; Fujisaki,Seiichiro; Shirakura,Masayuki; Watanabe,Shinji; Odagiri,Takato |
| EPI635513 | MP | India | 2015-Mar-30 | A/India/P158900/2015 | National Institute of Virology | National Institute of Infectious Diseases (NIID) | Takashita,Emi; Fujisaki,Seiichiro; Shirakura,Masayuki; Watanabe,Shinji; Odagiri,Takato |
| EPI635514 | PA | India | 2015-Mar-30 | A/India/P158900/2015 | National Institute of Virology | National Institute of Infectious Diseases (NIID) | Takashita,Emi; Fujisaki,Seiichiro; Shirakura,Masayuki; Watanabe,Shinji; Odagiri,Takato |
| EPI635515 | PB2 | India | 2015-Mar-30 | A/India/P158900/2015 | National Institute of Virology | National Institute of Infectious Diseases (NIID) | Takashita,Emi; Fujisaki,Seiichiro; Shirakura,Masayuki; Watanabe,Shinji; Odagiri,Takato |
| EPI635516 | PB1 | India | 2015-Mar-30 | A/India/P158900/2015 | National Institute of Virology | National Institute of Infectious Diseases (NIID) | Takashita,Emi; Fujisaki,Seiichiro; Shirakura,Masayuki; Watanabe,Shinji; Odagiri,Takato |
| EPI668468 | NP | Indonesia | 2015-Apr-29 | A/Indonesia/Nihrdi-Diy152/2015 | Ministry of Health, NIHRD | Centers for Disease Control and Prevention | NA |
| EPI668469 | NS | Indonesia | 2015-Apr-29 | A/Indonesia/Nihrdi-Diy152/2015 | Ministry of Health, NIHRD | Centers for Disease Control and Prevention | NA |
| EPI668470 | MP | Indonesia | 2015-Apr-29 | A/Indonesia/Nihrdi-Diy152/2015 | Ministry of Health, NIHRD | Centers for Disease Control and Prevention | NA |
| EPI668471 | PA | Indonesia | 2015-Apr-29 | A/Indonesia/Nihrdi-Diy152/2015 | Ministry of Health, NIHRD | Centers for Disease Control and Prevention | NA |
| EPI668472 | PB1 | Indonesia | 2015-Apr-29 | A/Indonesia/Nihrdi-Diy152/2015 | Ministry of Health, NIHRD | Centers for Disease Control and Prevention | NA |
| EPI668473 | PB2 | Indonesia | 2015-Apr-29 | A/Indonesia/Nihrdi-Diy152/2015 | Ministry of Health, NIHRD | Centers for Disease Control and Prevention | NA |
| EPI668474 | NA | Indonesia | 2015-Apr-29 | A/Indonesia/Nihrdi-Diy152/2015 | Ministry of Health, NIHRD | Centers for Disease Control and Prevention | NA |
| EPI668475 | HA | Indonesia | 2015-Apr-29 | A/Indonesia/Nihrdi-Diy152/2015 | Ministry of Health, NIHRD | Centers for Disease Control and Prevention | NA |
| EPI674922 | HA | Bangladesh | 2015-Aug-11 | A/Bangladesh/85006/2015 | Institute of Epidemiology Disease Control and Research (IEDCR) & Bangladesh National Influenza Centre (NIC) | Centers for Disease Control and Prevention | NA |
| EPI674915 | NP | Bangladesh | 2015-Aug-11 | A/Bangladesh/85006/2015 | Institute of Epidemiology Disease Control and Research (IEDCR) & Bangladesh National Influenza Centre (NIC) | Centers for Disease Control and Prevention | NA |
| EPI674916 | NS | Bangladesh | 2015-Aug-11 | A/Bangladesh/85006/2015 | Institute of Epidemiology Disease Control and Research (IEDCR) & Bangladesh National Influenza Centre (NIC) | Centers for Disease Control and Prevention | NA |
| EPI674917 | MP | Bangladesh | 2015-Aug-11 | A/Bangladesh/85006/2015 | Institute of Epidemiology Disease Control and Research (IEDCR) & Bangladesh National Influenza Centre (NIC) | Centers for Disease Control and Prevention | NA |
| EPI674918 | PA | Bangladesh | 2015-Aug-11 | A/Bangladesh/85006/2015 | Institute of Epidemiology Disease Control and Research (IEDCR) & Bangladesh National Influenza Centre (NIC) | Centers for Disease Control and Prevention | NA |
| EPI674919 | PB2 | Bangladesh | 2015-Aug-11 | A/Bangladesh/85006/2015 | Institute of Epidemiology Disease Control and Research (IEDCR) & Bangladesh National Influenza Centre (NIC) | Centers for Disease Control and Prevention | NA |
| EPI674920 | PB1 | Bangladesh | 2015-Aug-11 | A/Bangladesh/85006/2015 | Institute of Epidemiology Disease Control and Research (IEDCR) & Bangladesh National Influenza Centre (NIC) | Centers for Disease Control and Prevention | NA |
| EPI674921 | NA | Bangladesh | 2015-Aug-11 | A/Bangladesh/85006/2015 | Institute of Epidemiology Disease Control and Research (IEDCR) & Bangladesh National Influenza Centre (NIC) | Centers for Disease Control and Prevention | NA |
| EPI690993 | PA | Bangladesh | 2015-Oct-03 | A/Bangladesh/9001/2015 | Institute of Epidemiology Disease Control and Research (IEDCR) & Bangladesh National Influenza Centre (NIC) | Centers for Disease Control and Prevention | NA |
| EPI690994 | PB2 | Bangladesh | 2015-Oct-03 | A/Bangladesh/9001/2015 | Institute of Epidemiology Disease Control and Research (IEDCR) & Bangladesh National Influenza Centre (NIC) | Centers for Disease Control and Prevention | NA |
| EPI690995 | PB1 | Bangladesh | 2015-Oct-03 | A/Bangladesh/9001/2015 | Institute of Epidemiology Disease Control and Research (IEDCR) & Bangladesh National Influenza Centre (NIC) | Centers for Disease Control and Prevention | NA |
| EPI690990 | NP | Bangladesh | 2015-Oct-03 | A/Bangladesh/9001/2015 | Institute of Epidemiology Disease Control and Research (IEDCR) & Bangladesh National Influenza Centre (NIC) | Centers for Disease Control and Prevention | NA |
| EPI690991 | NS | Bangladesh | 2015-Oct-03 | A/Bangladesh/9001/2015 | Institute of Epidemiology Disease Control and Research (IEDCR) & Bangladesh National Influenza Centre (NIC) | Centers for Disease Control and Prevention | NA |
| EPI690992 | MP | Bangladesh | 2015-Oct-03 | A/Bangladesh/9001/2015 | Institute of Epidemiology Disease Control and Research (IEDCR) & Bangladesh National Influenza Centre (NIC) | Centers for Disease Control and Prevention | NA |
| EPI690996 | NA | Bangladesh | 2015-Oct-03 | A/Bangladesh/9001/2015 | Institute of Epidemiology Disease Control and Research (IEDCR) & Bangladesh National Influenza Centre (NIC) | Centers for Disease Control and Prevention | NA |
| EPI690997 | HA | Bangladesh | 2015-Oct-03 | A/Bangladesh/9001/2015 | Institute of Epidemiology Disease Control and Research (IEDCR) & Bangladesh National Influenza Centre (NIC) | Centers for Disease Control and Prevention | NA |
| EPI630420 | NA | India | 2015-Mar-04 | A/India/P1510348/2015 | National Institute of Virology | National Institute of Infectious Diseases (NIID) | Takashita,Emi; Fujisaki,Seiichiro; Shirakura,Masayuki; Watanabe,Shinji; Odagiri,Takato |
| EPI630421 | HA | India | 2015-Mar-04 | A/India/P1510348/2015 | National Institute of Virology | National Institute of Infectious Diseases (NIID) | Takashita,Emi; Fujisaki,Seiichiro; Shirakura,Masayuki; Watanabe,Shinji; Odagiri,Takato |
| EPI635523 | NP | India | 2015-Mar-04 | A/India/P1510348/2015 | National Institute of Virology | National Institute of Infectious Diseases (NIID) | Takashita,Emi; Fujisaki,Seiichiro; Shirakura,Masayuki; Watanabe,Shinji; Odagiri,Takato |
| EPI635524 | NS | India | 2015-Mar-04 | A/India/P1510348/2015 | National Institute of Virology | National Institute of Infectious Diseases (NIID) | Takashita,Emi; Fujisaki,Seiichiro; Shirakura,Masayuki; Watanabe,Shinji; Odagiri,Takato |
| EPI635525 | MP | India | 2015-Mar-04 | A/India/P1510348/2015 | National Institute of Virology | National Institute of Infectious Diseases (NIID) | Takashita,Emi; Fujisaki,Seiichiro; Shirakura,Masayuki; Watanabe,Shinji; Odagiri,Takato |
| EPI635526 | PA | India | 2015-Mar-04 | A/India/P1510348/2015 | National Institute of Virology | National Institute of Infectious Diseases (NIID) | Takashita,Emi; Fujisaki,Seiichiro; Shirakura,Masayuki; Watanabe,Shinji; Odagiri,Takato |
| EPI635527 | PB2 | India | 2015-Mar-04 | A/India/P1510348/2015 | National Institute of Virology | National Institute of Infectious Diseases (NIID) | Takashita,Emi; Fujisaki,Seiichiro; Shirakura,Masayuki; Watanabe,Shinji; Odagiri,Takato |
| EPI635528 | PB1 | India | 2015-Mar-04 | A/India/P1510348/2015 | National Institute of Virology | National Institute of Infectious Diseases (NIID) | Takashita,Emi; Fujisaki,Seiichiro; Shirakura,Masayuki; Watanabe,Shinji; Odagiri,Takato |
| EPI679298 | PB1 | United Kingdom | 2015-Nov-17 | A/England/364/2015 | Microbiology Services Colindale, Public Health England | Microbiology Services Colindale, Public Health England | Galiano,M. |
| EPI679293 | NP | United Kingdom | 2015-Nov-17 | A/England/364/2015 | Microbiology Services Colindale, Public Health England | Microbiology Services Colindale, Public Health England | Galiano,M. |
| EPI679294 | NS | United Kingdom | 2015-Nov-17 | A/England/364/2015 | Microbiology Services Colindale, Public Health England | Microbiology Services Colindale, Public Health England | Galiano,M. |
| EPI679295 | MP | United Kingdom | 2015-Nov-17 | A/England/364/2015 | Microbiology Services Colindale, Public Health England | Microbiology Services Colindale, Public Health England | Galiano,M. |
| EPI679296 | PA | United Kingdom | 2015-Nov-17 | A/England/364/2015 | Microbiology Services Colindale, Public Health England | Microbiology Services Colindale, Public Health England | Galiano,M. |
| EPI679297 | PB2 | United Kingdom | 2015-Nov-17 | A/England/364/2015 | Microbiology Services Colindale, Public Health England | Microbiology Services Colindale, Public Health England | Galiano,M. |
| EPI679299 | NA | United Kingdom | 2015-Nov-17 | A/England/364/2015 | Microbiology Services Colindale, Public Health England | Microbiology Services Colindale, Public Health England | Galiano,M. |
| EPI679300 | HA | United Kingdom | 2015-Nov-17 | A/England/364/2015 | Microbiology Services Colindale, Public Health England | Microbiology Services Colindale, Public Health England | Galiano,M. |
| EPI691174 | PA | United States | 2015-Nov-30 | A/Washington/67/2015 | Washington State Public Health Laboratory | Centers for Disease Control and Prevention | NA |
| EPI691175 | PB2 | United States | 2015-Nov-30 | A/Washington/67/2015 | Washington State Public Health Laboratory | Centers for Disease Control and Prevention | NA |
| EPI691176 | PB1 | United States | 2015-Nov-30 | A/Washington/67/2015 | Washington State Public Health Laboratory | Centers for Disease Control and Prevention | NA |
| EPI691171 | NP | United States | 2015-Nov-30 | A/Washington/67/2015 | Washington State Public Health Laboratory | Centers for Disease Control and Prevention | NA |
| EPI691172 | NS | United States | 2015-Nov-30 | A/Washington/67/2015 | Washington State Public Health Laboratory | Centers for Disease Control and Prevention | NA |
| EPI691173 | MP | United States | 2015-Nov-30 | A/Washington/67/2015 | Washington State Public Health Laboratory | Centers for Disease Control and Prevention | NA |
| EPI691177 | NA | United States | 2015-Nov-30 | A/Washington/67/2015 | Washington State Public Health Laboratory | Centers for Disease Control and Prevention | NA |
| EPI691178 | HA | United States | 2015-Nov-30 | A/Washington/67/2015 | Washington State Public Health Laboratory | Centers for Disease Control and Prevention | NA |
| EPI614487 | NP | India | 2014-Mar-06 | A/India/6427/2014 | National Institute of Virology | Centers for Disease Control and Prevention | NA |
| EPI614488 | NS | India | 2014-Mar-06 | A/India/6427/2014 | National Institute of Virology | Centers for Disease Control and Prevention | NA |
| EPI614489 | PA | India | 2014-Mar-06 | A/India/6427/2014 | National Institute of Virology | Centers for Disease Control and Prevention | NA |
| EPI614490 | PB2 | India | 2014-Mar-06 | A/India/6427/2014 | National Institute of Virology | Centers for Disease Control and Prevention | NA |
| EPI614491 | PB1 | India | 2014-Mar-06 | A/India/6427/2014 | National Institute of Virology | Centers for Disease Control and Prevention | NA |
| EPI537949 | MP | India | 2014-Mar-06 | A/India/6427/2014 | National Institute of Virology | Centers for Disease Control and Prevention | NA |
| EPI537950 | NA | India | 2014-Mar-06 | A/India/6427/2014 | National Institute of Virology | Centers for Disease Control and Prevention | NA |
| EPI537951 | HA | India | 2014-Mar-06 | A/India/6427/2014 | National Institute of Virology | Centers for Disease Control and Prevention | NA |
| EPI584968 | NP | India | 2015-Feb-01 | A/India/1399/2015 | National Institute of Virology | Centers for Disease Control and Prevention | NA |
| EPI584969 | NS | India | 2015-Feb-01 | A/India/1399/2015 | National Institute of Virology | Centers for Disease Control and Prevention | NA |
| EPI584970 | MP | India | 2015-Feb-01 | A/India/1399/2015 | National Institute of Virology | Centers for Disease Control and Prevention | NA |
| EPI584971 | PA | India | 2015-Feb-01 | A/India/1399/2015 | National Institute of Virology | Centers for Disease Control and Prevention | NA |
| EPI584972 | PB2 | India | 2015-Feb-01 | A/India/1399/2015 | National Institute of Virology | Centers for Disease Control and Prevention | NA |
| EPI584974 | PB1 | India | 2015-Feb-01 | A/India/1399/2015 | National Institute of Virology | Centers for Disease Control and Prevention | NA |
| EPI584975 | NA | India | 2015-Feb-01 | A/India/1399/2015 | National Institute of Virology | Centers for Disease Control and Prevention | NA |
| EPI584976 | HA | India | 2015-Feb-01 | A/India/1399/2015 | National Institute of Virology | Centers for Disease Control and Prevention | NA |
| EPI643444 | PB2 | Australia | 2015-Mar-12 | A/Brisbane/35/2015 | WHO Collaborating Centre for Reference and Research on Influenza | Centers for Disease Control and Prevention | NA |
| EPI643445 | PB1 | Australia | 2015-Mar-12 | A/Brisbane/35/2015 | WHO Collaborating Centre for Reference and Research on Influenza | Centers for Disease Control and Prevention | NA |
| EPI643440 | NP | Australia | 2015-Mar-12 | A/Brisbane/35/2015 | WHO Collaborating Centre for Reference and Research on Influenza | Centers for Disease Control and Prevention | NA |
| EPI643441 | NS | Australia | 2015-Mar-12 | A/Brisbane/35/2015 | WHO Collaborating Centre for Reference and Research on Influenza | Centers for Disease Control and Prevention | NA |
| EPI643442 | MP | Australia | 2015-Mar-12 | A/Brisbane/35/2015 | WHO Collaborating Centre for Reference and Research on Influenza | Centers for Disease Control and Prevention | NA |
| EPI643443 | PA | Australia | 2015-Mar-12 | A/Brisbane/35/2015 | WHO Collaborating Centre for Reference and Research on Influenza | Centers for Disease Control and Prevention | NA |
| EPI643446 | NA | Australia | 2015-Mar-12 | A/Brisbane/35/2015 | WHO Collaborating Centre for Reference and Research on Influenza | Centers for Disease Control and Prevention | NA |
| EPI643447 | HA | Australia | 2015-Mar-12 | A/Brisbane/35/2015 | WHO Collaborating Centre for Reference and Research on Influenza | Centers for Disease Control and Prevention | NA |
| EPI679001 | NP | Nigeria | 2015-Sep-21 | A/Nigeria/306/2015 | National Influenza Reference Laboratory | Centers for Disease Control and Prevention | NA |
| EPI679008 | HA | Nigeria | 2015-Sep-21 | A/Nigeria/306/2015 | National Influenza Reference Laboratory | Centers for Disease Control and Prevention | NA |
| EPI679002 | NS | Nigeria | 2015-Sep-21 | A/Nigeria/306/2015 | National Influenza Reference Laboratory | Centers for Disease Control and Prevention | NA |
| EPI679003 | MP | Nigeria | 2015-Sep-21 | A/Nigeria/306/2015 | National Influenza Reference Laboratory | Centers for Disease Control and Prevention | NA |
| EPI679004 | PA | Nigeria | 2015-Sep-21 | A/Nigeria/306/2015 | National Influenza Reference Laboratory | Centers for Disease Control and Prevention | NA |
| EPI679005 | PB2 | Nigeria | 2015-Sep-21 | A/Nigeria/306/2015 | National Influenza Reference Laboratory | Centers for Disease Control and Prevention | NA |
| EPI679006 | PB1 | Nigeria | 2015-Sep-21 | A/Nigeria/306/2015 | National Influenza Reference Laboratory | Centers for Disease Control and Prevention | NA |
| EPI679007 | NA | Nigeria | 2015-Sep-21 | A/Nigeria/306/2015 | National Influenza Reference Laboratory | Centers for Disease Control and Prevention | NA |
| EPI684368 | PB2 | United States | 2015-Nov-25 | A/Wisconsin/88/2015 | Wisconsin State Laboratory of Hygiene | Centers for Disease Control and Prevention | NA |
| EPI684364 | NP | United States | 2015-Nov-25 | A/Wisconsin/88/2015 | Wisconsin State Laboratory of Hygiene | Centers for Disease Control and Prevention | NA |
| EPI684365 | NS | United States | 2015-Nov-25 | A/Wisconsin/88/2015 | Wisconsin State Laboratory of Hygiene | Centers for Disease Control and Prevention | NA |
| EPI684366 | MP | United States | 2015-Nov-25 | A/Wisconsin/88/2015 | Wisconsin State Laboratory of Hygiene | Centers for Disease Control and Prevention | NA |
| EPI684367 | PA | United States | 2015-Nov-25 | A/Wisconsin/88/2015 | Wisconsin State Laboratory of Hygiene | Centers for Disease Control and Prevention | NA |
| EPI684369 | NA | United States | 2015-Nov-25 | A/Wisconsin/88/2015 | Wisconsin State Laboratory of Hygiene | Centers for Disease Control and Prevention | NA |
| EPI684370 | HA | United States | 2015-Nov-25 | A/Wisconsin/88/2015 | Wisconsin State Laboratory of Hygiene | Centers for Disease Control and Prevention | NA |
| EPI687193 | NP | Sweden | 2015-Nov-21 | A/Stockholm/67/2015 |  | Swedish Institute for Infectious Disease Control | NA |
| EPI687194 | PB1 | Sweden | 2015-Nov-21 | A/Stockholm/67/2015 |  | Swedish Institute for Infectious Disease Control | NA |
| EPI687195 | PB2 | Sweden | 2015-Nov-21 | A/Stockholm/67/2015 |  | Swedish Institute for Infectious Disease Control | NA |
| EPI687197 | MP | Sweden | 2015-Nov-21 | A/Stockholm/67/2015 |  | Swedish Institute for Infectious Disease Control | NA |
| EPI687198 | NA | Sweden | 2015-Nov-21 | A/Stockholm/67/2015 |  | Swedish Institute for Infectious Disease Control | NA |
| EPI687196 | NS | Sweden | 2015-Nov-21 | A/Stockholm/67/2015 |  | Swedish Institute for Infectious Disease Control | NA |
| EPI687199 | HA | Sweden | 2015-Nov-21 | A/Stockholm/67/2015 |  | Swedish Institute for Infectious Disease Control | NA |
| EPI687192 | PA | Sweden | 2015-Nov-21 | A/Stockholm/67/2015 |  | Swedish Institute for Infectious Disease Control | NA |
| EPI691166 | PA | United States | 2015-Dec-02 | A/California/116/2015 | California Department of Health Services | Centers for Disease Control and Prevention | NA |
| EPI691167 | PB2 | United States | 2015-Dec-02 | A/California/116/2015 | California Department of Health Services | Centers for Disease Control and Prevention | NA |
| EPI691168 | PB1 | United States | 2015-Dec-02 | A/California/116/2015 | California Department of Health Services | Centers for Disease Control and Prevention | NA |
| EPI691163 | NP | United States | 2015-Dec-02 | A/California/116/2015 | California Department of Health Services | Centers for Disease Control and Prevention | NA |
| EPI691164 | NS | United States | 2015-Dec-02 | A/California/116/2015 | California Department of Health Services | Centers for Disease Control and Prevention | NA |
| EPI691165 | MP | United States | 2015-Dec-02 | A/California/116/2015 | California Department of Health Services | Centers for Disease Control and Prevention | NA |
| EPI691169 | NA | United States | 2015-Dec-02 | A/California/116/2015 | California Department of Health Services | Centers for Disease Control and Prevention | NA |
| EPI691170 | HA | United States | 2015-Dec-02 | A/California/116/2015 | California Department of Health Services | Centers for Disease Control and Prevention | NA |
| EPI691307 | NP | Kazakhstan | 2015-Nov-23 | A/Kazakhstan/57/2015 | CSEE | Centers for Disease Control and Prevention | NA |
| EPI691308 | NS | Kazakhstan | 2015-Nov-23 | A/Kazakhstan/57/2015 | CSEE | Centers for Disease Control and Prevention | NA |
| EPI691309 | MP | Kazakhstan | 2015-Nov-23 | A/Kazakhstan/57/2015 | CSEE | Centers for Disease Control and Prevention | NA |
| EPI691310 | PA | Kazakhstan | 2015-Nov-23 | A/Kazakhstan/57/2015 | CSEE | Centers for Disease Control and Prevention | NA |
| EPI691311 | PB2 | Kazakhstan | 2015-Nov-23 | A/Kazakhstan/57/2015 | CSEE | Centers for Disease Control and Prevention | NA |
| EPI691312 | PB1 | Kazakhstan | 2015-Nov-23 | A/Kazakhstan/57/2015 | CSEE | Centers for Disease Control and Prevention | NA |
| EPI691313 | NA | Kazakhstan | 2015-Nov-23 | A/Kazakhstan/57/2015 | CSEE | Centers for Disease Control and Prevention | NA |
| EPI691314 | HA | Kazakhstan | 2015-Nov-23 | A/Kazakhstan/57/2015 | CSEE | Centers for Disease Control and Prevention | NA |
| EPI580473 | PB2 | Lebanon | 2014-Feb-07 | A/Lebanon/14L66/2014 |  | Niigata University | Saito, Reiko; Takemae, Nobuhiro; Saito, Takehiko; Shobugawa, Yugo; Kondo, Hiroki; Hibino, Akinobu |
| EPI580475 | PA | Lebanon | 2014-Feb-07 | A/Lebanon/14L66/2014 |  | Niigata University | Saito, Reiko; Takemae, Nobuhiro; Saito, Takehiko; Shobugawa, Yugo; Kondo, Hiroki; Hibino, Akinobu |
| EPI580480 | NA | Lebanon | 2014-Feb-07 | A/Lebanon/14L66/2014 |  | Niigata University | Saito, Reiko; Takemae, Nobuhiro; Saito, Takehiko; Shobugawa, Yugo; Kondo, Hiroki; Hibino, Akinobu |
| EPI580478 | NP | Lebanon | 2014-Feb-07 | A/Lebanon/14L66/2014 |  | Niigata University | Saito, Reiko; Takemae, Nobuhiro; Saito, Takehiko; Shobugawa, Yugo; Kondo, Hiroki; Hibino, Akinobu |
| EPI580476 | HA | Lebanon | 2014-Feb-07 | A/Lebanon/14L66/2014 |  | Niigata University | Saito, Reiko; Takemae, Nobuhiro; Saito, Takehiko; Shobugawa, Yugo; Kondo, Hiroki; Hibino, Akinobu |
| EPI580479 | NS | Lebanon | 2014-Feb-07 | A/Lebanon/14L66/2014 |  | Niigata University | Saito, Reiko; Takemae, Nobuhiro; Saito, Takehiko; Shobugawa, Yugo; Kondo, Hiroki; Hibino, Akinobu |
| EPI580474 | PB1 | Lebanon | 2014-Feb-07 | A/Lebanon/14L66/2014 |  | Niigata University | Saito, Reiko; Takemae, Nobuhiro; Saito, Takehiko; Shobugawa, Yugo; Kondo, Hiroki; Hibino, Akinobu |
| EPI580477 | MP | Lebanon | 2014-Feb-07 | A/Lebanon/14L66/2014 |  | Niigata University | Saito, Reiko; Takemae, Nobuhiro; Saito, Takehiko; Shobugawa, Yugo; Kondo, Hiroki; Hibino, Akinobu |
| EPI632423 | PB2 | Russian Federation | 2015-Apr-14 | A/Moscow/144/2015 | Ivanovsky Research Institute of Virology RAMS | Centers for Disease Control and Prevention | NA |
| EPI632424 | PB1 | Russian Federation | 2015-Apr-14 | A/Moscow/144/2015 | Ivanovsky Research Institute of Virology RAMS | Centers for Disease Control and Prevention | NA |
| EPI632419 | NP | Russian Federation | 2015-Apr-14 | A/Moscow/144/2015 | Ivanovsky Research Institute of Virology RAMS | Centers for Disease Control and Prevention | NA |
| EPI632420 | NS | Russian Federation | 2015-Apr-14 | A/Moscow/144/2015 | Ivanovsky Research Institute of Virology RAMS | Centers for Disease Control and Prevention | NA |
| EPI632421 | MP | Russian Federation | 2015-Apr-14 | A/Moscow/144/2015 | Ivanovsky Research Institute of Virology RAMS | Centers for Disease Control and Prevention | NA |
| EPI632422 | PA | Russian Federation | 2015-Apr-14 | A/Moscow/144/2015 | Ivanovsky Research Institute of Virology RAMS | Centers for Disease Control and Prevention | NA |
| EPI632425 | NA | Russian Federation | 2015-Apr-14 | A/Moscow/144/2015 | Ivanovsky Research Institute of Virology RAMS | Centers for Disease Control and Prevention | NA |
| EPI632426 | HA | Russian Federation | 2015-Apr-14 | A/Moscow/144/2015 | Ivanovsky Research Institute of Virology RAMS | Centers for Disease Control and Prevention | NA |
| EPI643362 | NP | Martinique | 2015-Mar-20 | A/Martinique/2131/2015 | National Influenza Center French Guiana and French Indies | Centers for Disease Control and Prevention | NA |
| EPI643363 | NS | Martinique | 2015-Mar-20 | A/Martinique/2131/2015 | National Influenza Center French Guiana and French Indies | Centers for Disease Control and Prevention | NA |
| EPI643364 | MP | Martinique | 2015-Mar-20 | A/Martinique/2131/2015 | National Influenza Center French Guiana and French Indies | Centers for Disease Control and Prevention | NA |
| EPI643365 | PA | Martinique | 2015-Mar-20 | A/Martinique/2131/2015 | National Influenza Center French Guiana and French Indies | Centers for Disease Control and Prevention | NA |
| EPI643366 | PB2 | Martinique | 2015-Mar-20 | A/Martinique/2131/2015 | National Influenza Center French Guiana and French Indies | Centers for Disease Control and Prevention | NA |
| EPI643367 | PB1 | Martinique | 2015-Mar-20 | A/Martinique/2131/2015 | National Influenza Center French Guiana and French Indies | Centers for Disease Control and Prevention | NA |
| EPI643368 | NA | Martinique | 2015-Mar-20 | A/Martinique/2131/2015 | National Influenza Center French Guiana and French Indies | Centers for Disease Control and Prevention | NA |
| EPI643369 | HA | Martinique | 2015-Mar-20 | A/Martinique/2131/2015 | National Influenza Center French Guiana and French Indies | Centers for Disease Control and Prevention | NA |
| EPI643472 | NP | Paraguay | 2015-Jun-19 | A/Paraguay/8203/2015 | Central Laboratory of Public Health | Centers for Disease Control and Prevention | NA |
| EPI643473 | NS | Paraguay | 2015-Jun-19 | A/Paraguay/8203/2015 | Central Laboratory of Public Health | Centers for Disease Control and Prevention | NA |
| EPI643474 | MP | Paraguay | 2015-Jun-19 | A/Paraguay/8203/2015 | Central Laboratory of Public Health | Centers for Disease Control and Prevention | NA |
| EPI643475 | PA | Paraguay | 2015-Jun-19 | A/Paraguay/8203/2015 | Central Laboratory of Public Health | Centers for Disease Control and Prevention | NA |
| EPI643478 | NA | Paraguay | 2015-Jun-19 | A/Paraguay/8203/2015 | Central Laboratory of Public Health | Centers for Disease Control and Prevention | NA |
| EPI643479 | HA | Paraguay | 2015-Jun-19 | A/Paraguay/8203/2015 | Central Laboratory of Public Health | Centers for Disease Control and Prevention | NA |
| EPI643476 | PB2 | Paraguay | 2015-Jun-19 | A/Paraguay/8203/2015 | Central Laboratory of Public Health | Centers for Disease Control and Prevention | NA |
| EPI643477 | PB1 | Paraguay | 2015-Jun-19 | A/Paraguay/8203/2015 | Central Laboratory of Public Health | Centers for Disease Control and Prevention | NA |
| EPI668444 | NP | United States | 2015-Sep-18 | A/Hawaii/59/2015 | State of Hawaii Department of Health | Centers for Disease Control and Prevention | NA |
| EPI668445 | NS | United States | 2015-Sep-18 | A/Hawaii/59/2015 | State of Hawaii Department of Health | Centers for Disease Control and Prevention | NA |
| EPI668446 | MP | United States | 2015-Sep-18 | A/Hawaii/59/2015 | State of Hawaii Department of Health | Centers for Disease Control and Prevention | NA |
| EPI668447 | PA | United States | 2015-Sep-18 | A/Hawaii/59/2015 | State of Hawaii Department of Health | Centers for Disease Control and Prevention | NA |
| EPI668448 | PB2 | United States | 2015-Sep-18 | A/Hawaii/59/2015 | State of Hawaii Department of Health | Centers for Disease Control and Prevention | NA |
| EPI668449 | PB1 | United States | 2015-Sep-18 | A/Hawaii/59/2015 | State of Hawaii Department of Health | Centers for Disease Control and Prevention | NA |
| EPI668450 | NA | United States | 2015-Sep-18 | A/Hawaii/59/2015 | State of Hawaii Department of Health | Centers for Disease Control and Prevention | NA |
| EPI668451 | HA | United States | 2015-Sep-18 | A/Hawaii/59/2015 | State of Hawaii Department of Health | Centers for Disease Control and Prevention | NA |
| EPI669739 | PB2 | Mexico | 2015-Mar-23 | A/Mexico/1374/2015 | Laboratorio de Virus Respiratorio | Centers for Disease Control and Prevention | NA |
| EPI669740 | PB1 | Mexico | 2015-Mar-23 | A/Mexico/1374/2015 | Laboratorio de Virus Respiratorio | Centers for Disease Control and Prevention | NA |
| EPI669735 | NP | Mexico | 2015-Mar-23 | A/Mexico/1374/2015 | Laboratorio de Virus Respiratorio | Centers for Disease Control and Prevention | NA |
| EPI669736 | NS | Mexico | 2015-Mar-23 | A/Mexico/1374/2015 | Laboratorio de Virus Respiratorio | Centers for Disease Control and Prevention | NA |
| EPI669737 | MP | Mexico | 2015-Mar-23 | A/Mexico/1374/2015 | Laboratorio de Virus Respiratorio | Centers for Disease Control and Prevention | NA |
| EPI669738 | PA | Mexico | 2015-Mar-23 | A/Mexico/1374/2015 | Laboratorio de Virus Respiratorio | Centers for Disease Control and Prevention | NA |
| EPI669741 | NA | Mexico | 2015-Mar-23 | A/Mexico/1374/2015 | Laboratorio de Virus Respiratorio | Centers for Disease Control and Prevention | NA |
| EPI669742 | HA | Mexico | 2015-Mar-23 | A/Mexico/1374/2015 | Laboratorio de Virus Respiratorio | Centers for Disease Control and Prevention | NA |
| EPI676383 | NS | Argentina | 2015-Jul-08 | A/Argentina/11551/2015 | CEMIC University Hospital | Centers for Disease Control and Prevention | NA |
| EPI669743 | NP | Argentina | 2015-Jul-08 | A/Argentina/11551/2015 | CEMIC University Hospital | Centers for Disease Control and Prevention | NA |
| EPI669744 | MP | Argentina | 2015-Jul-08 | A/Argentina/11551/2015 | CEMIC University Hospital | Centers for Disease Control and Prevention | NA |
| EPI669745 | PA | Argentina | 2015-Jul-08 | A/Argentina/11551/2015 | CEMIC University Hospital | Centers for Disease Control and Prevention | NA |
| EPI669746 | PB2 | Argentina | 2015-Jul-08 | A/Argentina/11551/2015 | CEMIC University Hospital | Centers for Disease Control and Prevention | NA |
| EPI669747 | PB1 | Argentina | 2015-Jul-08 | A/Argentina/11551/2015 | CEMIC University Hospital | Centers for Disease Control and Prevention | NA |
| EPI669748 | NA | Argentina | 2015-Jul-08 | A/Argentina/11551/2015 | CEMIC University Hospital | Centers for Disease Control and Prevention | NA |
| EPI669749 | HA | Argentina | 2015-Jul-08 | A/Argentina/11551/2015 | CEMIC University Hospital | Centers for Disease Control and Prevention | NA |
| EPI677726 | NP | United Kingdom | 2013-Dec-22 | A/England/673/2013 | Microbiology Services Colindale, Public Health England | Microbiology Services Colindale, Public Health England | Galiano,M. |
| EPI677727 | NS | United Kingdom | 2013-Dec-22 | A/England/673/2013 | Microbiology Services Colindale, Public Health England | Microbiology Services Colindale, Public Health England | Galiano,M. |
| EPI677728 | MP | United Kingdom | 2013-Dec-22 | A/England/673/2013 | Microbiology Services Colindale, Public Health England | Microbiology Services Colindale, Public Health England | Galiano,M. |
| EPI677729 | PA | United Kingdom | 2013-Dec-22 | A/England/673/2013 | Microbiology Services Colindale, Public Health England | Microbiology Services Colindale, Public Health England | Galiano,M. |
| EPI677730 | PB2 | United Kingdom | 2013-Dec-22 | A/England/673/2013 | Microbiology Services Colindale, Public Health England | Microbiology Services Colindale, Public Health England | Galiano,M. |
| EPI677731 | PB1 | United Kingdom | 2013-Dec-22 | A/England/673/2013 | Microbiology Services Colindale, Public Health England | Microbiology Services Colindale, Public Health England | Galiano,M. |
| EPI677732 | NA | United Kingdom | 2013-Dec-22 | A/England/673/2013 | Microbiology Services Colindale, Public Health England | Microbiology Services Colindale, Public Health England | Galiano,M. |
| EPI677733 | HA | United Kingdom | 2013-Dec-22 | A/England/673/2013 | Microbiology Services Colindale, Public Health England | Microbiology Services Colindale, Public Health England | Galiano,M. |
| EPI680477 | NA | Peru | 2015-Jul-21 | A/Peru/52/2015 | US NAMRU-6 | Centers for Disease Control and Prevention | NA |
| EPI680478 | HA | Peru | 2015-Jul-21 | A/Peru/52/2015 | US NAMRU-6 | Centers for Disease Control and Prevention | NA |
| EPI680475 | PB2 | Peru | 2015-Jul-21 | A/Peru/52/2015 | US NAMRU-6 | Centers for Disease Control and Prevention | NA |
| EPI680476 | PB1 | Peru | 2015-Jul-21 | A/Peru/52/2015 | US NAMRU-6 | Centers for Disease Control and Prevention | NA |
| EPI680471 | NP | Peru | 2015-Jul-21 | A/Peru/52/2015 | US NAMRU-6 | Centers for Disease Control and Prevention | NA |
| EPI680472 | NS | Peru | 2015-Jul-21 | A/Peru/52/2015 | US NAMRU-6 | Centers for Disease Control and Prevention | NA |
| EPI680473 | MP | Peru | 2015-Jul-21 | A/Peru/52/2015 | US NAMRU-6 | Centers for Disease Control and Prevention | NA |
| EPI680474 | PA | Peru | 2015-Jul-21 | A/Peru/52/2015 | US NAMRU-6 | Centers for Disease Control and Prevention | NA |
| EPI551351 | MP | New Zealand | 2014-Sep-10 | A/AUCKLAND/8/2014 | Auckland Hospital | WHO Collaborating Centre for Reference and Research on Influenza | Deng,Y-M.; Iannello,P.; Spirason,N.; Jelley,L.; Lau,H.; Komadina,N. |
| EPI551352 | NA | New Zealand | 2014-Sep-10 | A/AUCKLAND/8/2014 | Auckland Hospital | WHO Collaborating Centre for Reference and Research on Influenza | Deng,Y-M.; Iannello,P.; Spirason,N.; Jelley,L.; Lau,H.; Komadina,N. |
| EPI551353 | HA | New Zealand | 2014-Sep-10 | A/AUCKLAND/8/2014 | Auckland Hospital | WHO Collaborating Centre for Reference and Research on Influenza | Deng,Y-M.; Iannello,P.; Spirason,N.; Jelley,L.; Lau,H.; Komadina,N. |
| EPI561925 | NP | New Zealand | 2014-Sep-10 | A/AUCKLAND/8/2014 | Auckland Hospital | WHO Collaborating Centre for Reference and Research on Influenza | Deng,Y-M.; Iannello,P.; Spirason,N.; Jelley,L.; Lau,H.; Komadina,N. |
| EPI561926 | NS | New Zealand | 2014-Sep-10 | A/AUCKLAND/8/2014 | Auckland Hospital | WHO Collaborating Centre for Reference and Research on Influenza | Deng,Y-M.; Iannello,P.; Spirason,N.; Jelley,L.; Lau,H.; Komadina,N. |
| EPI561927 | PA | New Zealand | 2014-Sep-10 | A/AUCKLAND/8/2014 | Auckland Hospital | WHO Collaborating Centre for Reference and Research on Influenza | Deng,Y-M.; Iannello,P.; Spirason,N.; Jelley,L.; Lau,H.; Komadina,N. |
| EPI561928 | PB2 | New Zealand | 2014-Sep-10 | A/AUCKLAND/8/2014 | Auckland Hospital | WHO Collaborating Centre for Reference and Research on Influenza | Deng,Y-M.; Iannello,P.; Spirason,N.; Jelley,L.; Lau,H.; Komadina,N. |
| EPI561929 | PB1 | New Zealand | 2014-Sep-10 | A/AUCKLAND/8/2014 | Auckland Hospital | WHO Collaborating Centre for Reference and Research on Influenza | Deng,Y-M.; Iannello,P.; Spirason,N.; Jelley,L.; Lau,H.; Komadina,N. |
| EPI564945 | NP | Korea, Republic of | 2014-Dec-22 | A/Daegu/2001/2014 | National Institute of Health | Centers for Disease Control and Prevention | NA |
| EPI564946 | NS | Korea, Republic of | 2014-Dec-22 | A/Daegu/2001/2014 | National Institute of Health | Centers for Disease Control and Prevention | NA |
| EPI564947 | MP | Korea, Republic of | 2014-Dec-22 | A/Daegu/2001/2014 | National Institute of Health | Centers for Disease Control and Prevention | NA |
| EPI564948 | PA | Korea, Republic of | 2014-Dec-22 | A/Daegu/2001/2014 | National Institute of Health | Centers for Disease Control and Prevention | NA |
| EPI564949 | PB2 | Korea, Republic of | 2014-Dec-22 | A/Daegu/2001/2014 | National Institute of Health | Centers for Disease Control and Prevention | NA |
| EPI564950 | PB1 | Korea, Republic of | 2014-Dec-22 | A/Daegu/2001/2014 | National Institute of Health | Centers for Disease Control and Prevention | NA |
| EPI564951 | NA | Korea, Republic of | 2014-Dec-22 | A/Daegu/2001/2014 | National Institute of Health | Centers for Disease Control and Prevention | NA |
| EPI564952 | HA | Korea, Republic of | 2014-Dec-22 | A/Daegu/2001/2014 | National Institute of Health | Centers for Disease Control and Prevention | NA |
| EPI567729 | NP | Germany | 2014-Dec-19 | A/Germany/51/2014 | U.S. Air Force School of Aerospace Medicine | Centers for Disease Control and Prevention | NA |
| EPI567731 | MP | Germany | 2014-Dec-19 | A/Germany/51/2014 | U.S. Air Force School of Aerospace Medicine | Centers for Disease Control and Prevention | NA |
| EPI567730 | NS | Germany | 2014-Dec-19 | A/Germany/51/2014 | U.S. Air Force School of Aerospace Medicine | Centers for Disease Control and Prevention | NA |
| EPI567732 | PA | Germany | 2014-Dec-19 | A/Germany/51/2014 | U.S. Air Force School of Aerospace Medicine | Centers for Disease Control and Prevention | NA |
| EPI567733 | PB2 | Germany | 2014-Dec-19 | A/Germany/51/2014 | U.S. Air Force School of Aerospace Medicine | Centers for Disease Control and Prevention | NA |
| EPI567734 | PB1 | Germany | 2014-Dec-19 | A/Germany/51/2014 | U.S. Air Force School of Aerospace Medicine | Centers for Disease Control and Prevention | NA |
| EPI567735 | NA | Germany | 2014-Dec-19 | A/Germany/51/2014 | U.S. Air Force School of Aerospace Medicine | Centers for Disease Control and Prevention | NA |
| EPI567736 | HA | Germany | 2014-Dec-19 | A/Germany/51/2014 | U.S. Air Force School of Aerospace Medicine | Centers for Disease Control and Prevention | NA |
| EPI611074 | NP | United States | 2015-Apr-02 | A/Hawaii/18/2015 | State of Hawaii Department of Health | Centers for Disease Control and Prevention | NA |
| EPI611075 | NS | United States | 2015-Apr-02 | A/Hawaii/18/2015 | State of Hawaii Department of Health | Centers for Disease Control and Prevention | NA |
| EPI611076 | MP | United States | 2015-Apr-02 | A/Hawaii/18/2015 | State of Hawaii Department of Health | Centers for Disease Control and Prevention | NA |
| EPI611077 | PA | United States | 2015-Apr-02 | A/Hawaii/18/2015 | State of Hawaii Department of Health | Centers for Disease Control and Prevention | NA |
| EPI611080 | NA | United States | 2015-Apr-02 | A/Hawaii/18/2015 | State of Hawaii Department of Health | Centers for Disease Control and Prevention | NA |
| EPI611081 | HA | United States | 2015-Apr-02 | A/Hawaii/18/2015 | State of Hawaii Department of Health | Centers for Disease Control and Prevention | NA |
| EPI611078 | PB2 | United States | 2015-Apr-02 | A/Hawaii/18/2015 | State of Hawaii Department of Health | Centers for Disease Control and Prevention | NA |
| EPI611079 | PB1 | United States | 2015-Apr-02 | A/Hawaii/18/2015 | State of Hawaii Department of Health | Centers for Disease Control and Prevention | NA |
| EPI625627 | NA | Korea, Republic of | 2015-Feb-16 | A/Gyeongbuk/557/2015 | National Institute of Health | Centers for Disease Control and Prevention | NA |
| EPI625628 | HA | Korea, Republic of | 2015-Feb-16 | A/Gyeongbuk/557/2015 | National Institute of Health | Centers for Disease Control and Prevention | NA |
| EPI625625 | PB2 | Korea, Republic of | 2015-Feb-16 | A/Gyeongbuk/557/2015 | National Institute of Health | Centers for Disease Control and Prevention | NA |
| EPI625626 | PB1 | Korea, Republic of | 2015-Feb-16 | A/Gyeongbuk/557/2015 | National Institute of Health | Centers for Disease Control and Prevention | NA |
| EPI625621 | NP | Korea, Republic of | 2015-Feb-16 | A/Gyeongbuk/557/2015 | National Institute of Health | Centers for Disease Control and Prevention | NA |
| EPI625622 | NS | Korea, Republic of | 2015-Feb-16 | A/Gyeongbuk/557/2015 | National Institute of Health | Centers for Disease Control and Prevention | NA |
| EPI625623 | MP | Korea, Republic of | 2015-Feb-16 | A/Gyeongbuk/557/2015 | National Institute of Health | Centers for Disease Control and Prevention | NA |
| EPI625624 | PA | Korea, Republic of | 2015-Feb-16 | A/Gyeongbuk/557/2015 | National Institute of Health | Centers for Disease Control and Prevention | NA |
| EPI626296 | PB2 | Ukraine | 2015-Mar-08 | A/Ukraine/292/2015 | Institute of Epidemiology and Infectious Diseases AMS of Ukraine | Centers for Disease Control and Prevention | NA |
| EPI626297 | PB1 | Ukraine | 2015-Mar-08 | A/Ukraine/292/2015 | Institute of Epidemiology and Infectious Diseases AMS of Ukraine | Centers for Disease Control and Prevention | NA |
| EPI626292 | NP | Ukraine | 2015-Mar-08 | A/Ukraine/292/2015 | Institute of Epidemiology and Infectious Diseases AMS of Ukraine | Centers for Disease Control and Prevention | NA |
| EPI626293 | NS | Ukraine | 2015-Mar-08 | A/Ukraine/292/2015 | Institute of Epidemiology and Infectious Diseases AMS of Ukraine | Centers for Disease Control and Prevention | NA |
| EPI626294 | MP | Ukraine | 2015-Mar-08 | A/Ukraine/292/2015 | Institute of Epidemiology and Infectious Diseases AMS of Ukraine | Centers for Disease Control and Prevention | NA |
| EPI626295 | PA | Ukraine | 2015-Mar-08 | A/Ukraine/292/2015 | Institute of Epidemiology and Infectious Diseases AMS of Ukraine | Centers for Disease Control and Prevention | NA |
| EPI626298 | NA | Ukraine | 2015-Mar-08 | A/Ukraine/292/2015 | Institute of Epidemiology and Infectious Diseases AMS of Ukraine | Centers for Disease Control and Prevention | NA |
| EPI626299 | HA | Ukraine | 2015-Mar-08 | A/Ukraine/292/2015 | Institute of Epidemiology and Infectious Diseases AMS of Ukraine | Centers for Disease Control and Prevention | NA |
| EPI511882 | NP | United States | 2014-Feb-03 | A/California/06/2014 | California Department of Health Services | Centers for Disease Control and Prevention | NA |
| EPI511883 | NS | United States | 2014-Feb-03 | A/California/06/2014 | California Department of Health Services | Centers for Disease Control and Prevention | NA |
| EPI511884 | MP | United States | 2014-Feb-03 | A/California/06/2014 | California Department of Health Services | Centers for Disease Control and Prevention | NA |
| EPI511885 | PA | United States | 2014-Feb-03 | A/California/06/2014 | California Department of Health Services | Centers for Disease Control and Prevention | NA |
| EPI511886 | PB2 | United States | 2014-Feb-03 | A/California/06/2014 | California Department of Health Services | Centers for Disease Control and Prevention | NA |
| EPI511887 | PB1 | United States | 2014-Feb-03 | A/California/06/2014 | California Department of Health Services | Centers for Disease Control and Prevention | NA |
| EPI511888 | NA | United States | 2014-Feb-03 | A/California/06/2014 | California Department of Health Services | Centers for Disease Control and Prevention | NA |
| EPI511889 | HA | United States | 2014-Feb-03 | A/California/06/2014 | California Department of Health Services | Centers for Disease Control and Prevention | NA |
| EPI544135 | PB2 | Singapore | 2013-Dec-30 | A/Singapore/TT1061/2013 | Ministry of Health, Singapore | Ministry of Health, Singapore | Chen,B.B.; Phuah,S.P.; Poh,M.K.; Chen,S.L.; Cui,L. |
| EPI544136 | PB1 | Singapore | 2013-Dec-30 | A/Singapore/TT1061/2013 | Ministry of Health, Singapore | Ministry of Health, Singapore | Chen,B.B.; Phuah,S.P.; Poh,M.K.; Chen,S.L.; Cui,L. |
| EPI544131 | NP | Singapore | 2013-Dec-30 | A/Singapore/TT1061/2013 | Ministry of Health, Singapore | Ministry of Health, Singapore | Chen,B.B.; Phuah,S.P.; Poh,M.K.; Chen,S.L.; Cui,L. |
| EPI544132 | NS | Singapore | 2013-Dec-30 | A/Singapore/TT1061/2013 | Ministry of Health, Singapore | Ministry of Health, Singapore | Chen,B.B.; Phuah,S.P.; Poh,M.K.; Chen,S.L.; Cui,L. |
| EPI544133 | MP | Singapore | 2013-Dec-30 | A/Singapore/TT1061/2013 | Ministry of Health, Singapore | Ministry of Health, Singapore | Chen,B.B.; Phuah,S.P.; Poh,M.K.; Chen,S.L.; Cui,L. |
| EPI544134 | PA | Singapore | 2013-Dec-30 | A/Singapore/TT1061/2013 | Ministry of Health, Singapore | Ministry of Health, Singapore | Chen,B.B.; Phuah,S.P.; Poh,M.K.; Chen,S.L.; Cui,L. |
| EPI544137 | NA | Singapore | 2013-Dec-30 | A/Singapore/TT1061/2013 | Ministry of Health, Singapore | Ministry of Health, Singapore | Chen,B.B.; Phuah,S.P.; Poh,M.K.; Chen,S.L.; Cui,L. |
| EPI544138 | HA | Singapore | 2013-Dec-30 | A/Singapore/TT1061/2013 | Ministry of Health, Singapore | Ministry of Health, Singapore | Chen,B.B.; Phuah,S.P.; Poh,M.K.; Chen,S.L.; Cui,L. |
| EPI551823 | HA | Australia | 2014-Aug-15 | A/BRISBANE/1007/2014 | Institute of Medical and Veterinary Science (IMVS) | WHO Collaborating Centre for Reference and Research on Influenza | Deng,Y-M.; Iannello,P.; Spirason,N.; Jelley,L.; Lau,H.; Komadina,N. |
| EPI561930 | NP | Australia | 2014-Aug-15 | A/BRISBANE/1007/2014 | Institute of Medical and Veterinary Science (IMVS) | WHO Collaborating Centre for Reference and Research on Influenza | Deng,Y-M.; Iannello,P.; Spirason,N.; Jelley,L.; Lau,H.; Komadina,N. |
| EPI561931 | NS | Australia | 2014-Aug-15 | A/BRISBANE/1007/2014 | Institute of Medical and Veterinary Science (IMVS) | WHO Collaborating Centre for Reference and Research on Influenza | Deng,Y-M.; Iannello,P.; Spirason,N.; Jelley,L.; Lau,H.; Komadina,N. |
| EPI561932 | PA | Australia | 2014-Aug-15 | A/BRISBANE/1007/2014 | Institute of Medical and Veterinary Science (IMVS) | WHO Collaborating Centre for Reference and Research on Influenza | Deng,Y-M.; Iannello,P.; Spirason,N.; Jelley,L.; Lau,H.; Komadina,N. |
| EPI561933 | PB2 | Australia | 2014-Aug-15 | A/BRISBANE/1007/2014 | Institute of Medical and Veterinary Science (IMVS) | WHO Collaborating Centre for Reference and Research on Influenza | Deng,Y-M.; Iannello,P.; Spirason,N.; Jelley,L.; Lau,H.; Komadina,N. |
| EPI561934 | PB1 | Australia | 2014-Aug-15 | A/BRISBANE/1007/2014 | Institute of Medical and Veterinary Science (IMVS) | WHO Collaborating Centre for Reference and Research on Influenza | Deng,Y-M.; Iannello,P.; Spirason,N.; Jelley,L.; Lau,H.; Komadina,N. |
| EPI551821 | MP | Australia | 2014-Aug-15 | A/BRISBANE/1007/2014 | Institute of Medical and Veterinary Science (IMVS) | WHO Collaborating Centre for Reference and Research on Influenza | Deng,Y-M.; Iannello,P.; Spirason,N.; Jelley,L.; Lau,H.; Komadina,N. |
| EPI551822 | NA | Australia | 2014-Aug-15 | A/BRISBANE/1007/2014 | Institute of Medical and Veterinary Science (IMVS) | WHO Collaborating Centre for Reference and Research on Influenza | Deng,Y-M.; Iannello,P.; Spirason,N.; Jelley,L.; Lau,H.; Komadina,N. |
| EPI567848 | NP | Japan | 2014-Jan-25 | A/Nagasaki/13N057/2014 |  | Niigata University | Saito, Reiko; Takemae, Nobuhiro; Saito, Takehiko; Shobugawa, Yugo; Kondo, Hiroki; Hibino, Akinobu; Masaki, Hironori |
| EPI567841 | PA | Japan | 2014-Jan-25 | A/Nagasaki/13N057/2014 |  | Niigata University | Saito, Reiko; Takemae, Nobuhiro; Saito, Takehiko; Shobugawa, Yugo; Kondo, Hiroki; Hibino, Akinobu; Masaki, Hironori |
| EPI567845 | PB2 | Japan | 2014-Jan-25 | A/Nagasaki/13N057/2014 |  | Niigata University | Saito, Reiko; Takemae, Nobuhiro; Saito, Takehiko; Shobugawa, Yugo; Kondo, Hiroki; Hibino, Akinobu; Masaki, Hironori |
| EPI567842 | PB1 | Japan | 2014-Jan-25 | A/Nagasaki/13N057/2014 |  | Niigata University | Saito, Reiko; Takemae, Nobuhiro; Saito, Takehiko; Shobugawa, Yugo; Kondo, Hiroki; Hibino, Akinobu; Masaki, Hironori |
| EPI567843 | HA | Japan | 2014-Jan-25 | A/Nagasaki/13N057/2014 |  | Niigata University | Saito, Reiko; Takemae, Nobuhiro; Saito, Takehiko; Shobugawa, Yugo; Kondo, Hiroki; Hibino, Akinobu; Masaki, Hironori |
| EPI567844 | MP | Japan | 2014-Jan-25 | A/Nagasaki/13N057/2014 |  | Niigata University | Saito, Reiko; Takemae, Nobuhiro; Saito, Takehiko; Shobugawa, Yugo; Kondo, Hiroki; Hibino, Akinobu; Masaki, Hironori |
| EPI567847 | NS | Japan | 2014-Jan-25 | A/Nagasaki/13N057/2014 |  | Niigata University | Saito, Reiko; Takemae, Nobuhiro; Saito, Takehiko; Shobugawa, Yugo; Kondo, Hiroki; Hibino, Akinobu; Masaki, Hironori |
| EPI567846 | NA | Japan | 2014-Jan-25 | A/Nagasaki/13N057/2014 |  | Niigata University | Saito, Reiko; Takemae, Nobuhiro; Saito, Takehiko; Shobugawa, Yugo; Kondo, Hiroki; Hibino, Akinobu; Masaki, Hironori |
| EPI568492 | MP | United States | 2014-Sep-03 | A/Alaska/38/2014 | Alaska State Virology Lab | Centers for Disease Control and Prevention | NA |
| EPI568493 | PA | United States | 2014-Sep-03 | A/Alaska/38/2014 | Alaska State Virology Lab | Centers for Disease Control and Prevention | NA |
| EPI568494 | PB2 | United States | 2014-Sep-03 | A/Alaska/38/2014 | Alaska State Virology Lab | Centers for Disease Control and Prevention | NA |
| EPI568495 | NA | United States | 2014-Sep-03 | A/Alaska/38/2014 | Alaska State Virology Lab | Centers for Disease Control and Prevention | NA |
| EPI569247 | NP | United States | 2014-Sep-03 | A/Alaska/38/2014 | Alaska State Virology Lab | Centers for Disease Control and Prevention | NA |
| EPI569248 | NS | United States | 2014-Sep-03 | A/Alaska/38/2014 | Alaska State Virology Lab | Centers for Disease Control and Prevention | NA |
| EPI572284 | PB1 | United States | 2014-Sep-03 | A/Alaska/38/2014 | Alaska State Virology Lab | Centers for Disease Control and Prevention | NA |
| EPI577019 | HA | United States | 2014-Sep-03 | A/Alaska/38/2014 | Alaska State Virology Lab | Centers for Disease Control and Prevention | NA |
| EPI509419 | PA | United States | 2014-Jan-18 | A/Texas/09/2014 | Texas Department of State Health Services-Laboratory Services | Centers for Disease Control and Prevention | NA |
| EPI509420 | PB2 | United States | 2014-Jan-18 | A/Texas/09/2014 | Texas Department of State Health Services-Laboratory Services | Centers for Disease Control and Prevention | NA |
| EPI507532 | NP | United States | 2014-Jan-18 | A/Texas/09/2014 | Texas Department of State Health Services-Laboratory Services | Centers for Disease Control and Prevention | NA |
| EPI507533 | NS | United States | 2014-Jan-18 | A/Texas/09/2014 | Texas Department of State Health Services-Laboratory Services | Centers for Disease Control and Prevention | NA |
| EPI507534 | MP | United States | 2014-Jan-18 | A/Texas/09/2014 | Texas Department of State Health Services-Laboratory Services | Centers for Disease Control and Prevention | NA |
| EPI507535 | PB1 | United States | 2014-Jan-18 | A/Texas/09/2014 | Texas Department of State Health Services-Laboratory Services | Centers for Disease Control and Prevention | NA |
| EPI507536 | NA | United States | 2014-Jan-18 | A/Texas/09/2014 | Texas Department of State Health Services-Laboratory Services | Centers for Disease Control and Prevention | NA |
| EPI507537 | HA | United States | 2014-Jan-18 | A/Texas/09/2014 | Texas Department of State Health Services-Laboratory Services | Centers for Disease Control and Prevention | NA |
| EPI587764 | PB1 | Egypt | 2014-Oct-04 | A/Egypt/4372/2014 | U.S. Naval Medical Research Unit No.3 | Centers for Disease Control and Prevention | NA |
| EPI587759 | NP | Egypt | 2014-Oct-04 | A/Egypt/4372/2014 | U.S. Naval Medical Research Unit No.3 | Centers for Disease Control and Prevention | NA |
| EPI587760 | NS | Egypt | 2014-Oct-04 | A/Egypt/4372/2014 | U.S. Naval Medical Research Unit No.3 | Centers for Disease Control and Prevention | NA |
| EPI587761 | MP | Egypt | 2014-Oct-04 | A/Egypt/4372/2014 | U.S. Naval Medical Research Unit No.3 | Centers for Disease Control and Prevention | NA |
| EPI587762 | PA | Egypt | 2014-Oct-04 | A/Egypt/4372/2014 | U.S. Naval Medical Research Unit No.3 | Centers for Disease Control and Prevention | NA |
| EPI587763 | PB2 | Egypt | 2014-Oct-04 | A/Egypt/4372/2014 | U.S. Naval Medical Research Unit No.3 | Centers for Disease Control and Prevention | NA |
| EPI587765 | NA | Egypt | 2014-Oct-04 | A/Egypt/4372/2014 | U.S. Naval Medical Research Unit No.3 | Centers for Disease Control and Prevention | NA |
| EPI587766 | HA | Egypt | 2014-Oct-04 | A/Egypt/4372/2014 | U.S. Naval Medical Research Unit No.3 | Centers for Disease Control and Prevention | NA |
| EPI508612 | NP | United States | 2013-Dec-30 | A/California/59/2013 | California Department of Health Services | Centers for Disease Control and Prevention | NA |
| EPI508613 | NS | United States | 2013-Dec-30 | A/California/59/2013 | California Department of Health Services | Centers for Disease Control and Prevention | NA |
| EPI508614 | MP | United States | 2013-Dec-30 | A/California/59/2013 | California Department of Health Services | Centers for Disease Control and Prevention | NA |
| EPI508615 | PA | United States | 2013-Dec-30 | A/California/59/2013 | California Department of Health Services | Centers for Disease Control and Prevention | NA |
| EPI508616 | PB2 | United States | 2013-Dec-30 | A/California/59/2013 | California Department of Health Services | Centers for Disease Control and Prevention | NA |
| EPI508617 | PB1 | United States | 2013-Dec-30 | A/California/59/2013 | California Department of Health Services | Centers for Disease Control and Prevention | NA |
| EPI508618 | NA | United States | 2013-Dec-30 | A/California/59/2013 | California Department of Health Services | Centers for Disease Control and Prevention | NA |
| EPI508619 | HA | United States | 2013-Dec-30 | A/California/59/2013 | California Department of Health Services | Centers for Disease Control and Prevention | NA |
| EPI516709 | NP | United States | 2013-Dec-25 | A/Louisiana/41/2013 | New York State Department of Health | Centers for Disease Control and Prevention | NA |
| EPI516710 | MP | United States | 2013-Dec-25 | A/Louisiana/41/2013 | New York State Department of Health | Centers for Disease Control and Prevention | NA |
| EPI516711 | PA | United States | 2013-Dec-25 | A/Louisiana/41/2013 | New York State Department of Health | Centers for Disease Control and Prevention | NA |
| EPI516712 | PB2 | United States | 2013-Dec-25 | A/Louisiana/41/2013 | New York State Department of Health | Centers for Disease Control and Prevention | NA |
| EPI516713 | PB1 | United States | 2013-Dec-25 | A/Louisiana/41/2013 | New York State Department of Health | Centers for Disease Control and Prevention | NA |
| EPI516714 | NA | United States | 2013-Dec-25 | A/Louisiana/41/2013 | New York State Department of Health | Centers for Disease Control and Prevention | NA |
| EPI516715 | HA | United States | 2013-Dec-25 | A/Louisiana/41/2013 | New York State Department of Health | Centers for Disease Control and Prevention | NA |
| EPI534036 | NS | United States | 2013-Dec-25 | A/Louisiana/41/2013 | New York State Department of Health | Centers for Disease Control and Prevention | NA |
| EPI529393 | NP | Australia | 2013-Nov-11 | A/SYDNEY/82/2013 | Clinical Virology Unit, CDIM | WHO Collaborating Centre for Reference and Research on Influenza | Deng,Y-M.; Iannello,P.; Spirason,N.; Jelley,L.; Lau,H.; Komadina,N. |
| EPI529394 | NS | Australia | 2013-Nov-11 | A/SYDNEY/82/2013 | Clinical Virology Unit, CDIM | WHO Collaborating Centre for Reference and Research on Influenza | Deng,Y-M.; Iannello,P.; Spirason,N.; Jelley,L.; Lau,H.; Komadina,N. |
| EPI529395 | MP | Australia | 2013-Nov-11 | A/SYDNEY/82/2013 | Clinical Virology Unit, CDIM | WHO Collaborating Centre for Reference and Research on Influenza | Deng,Y-M.; Iannello,P.; Spirason,N.; Jelley,L.; Lau,H.; Komadina,N. |
| EPI529396 | PA | Australia | 2013-Nov-11 | A/SYDNEY/82/2013 | Clinical Virology Unit, CDIM | WHO Collaborating Centre for Reference and Research on Influenza | Deng,Y-M.; Iannello,P.; Spirason,N.; Jelley,L.; Lau,H.; Komadina,N. |
| EPI529397 | PB2 | Australia | 2013-Nov-11 | A/SYDNEY/82/2013 | Clinical Virology Unit, CDIM | WHO Collaborating Centre for Reference and Research on Influenza | Deng,Y-M.; Iannello,P.; Spirason,N.; Jelley,L.; Lau,H.; Komadina,N. |
| EPI529398 | PB1 | Australia | 2013-Nov-11 | A/SYDNEY/82/2013 | Clinical Virology Unit, CDIM | WHO Collaborating Centre for Reference and Research on Influenza | Deng,Y-M.; Iannello,P.; Spirason,N.; Jelley,L.; Lau,H.; Komadina,N. |
| EPI529399 | NA | Australia | 2013-Nov-11 | A/SYDNEY/82/2013 | Clinical Virology Unit, CDIM | WHO Collaborating Centre for Reference and Research on Influenza | Deng,Y-M.; Iannello,P.; Spirason,N.; Jelley,L.; Lau,H.; Komadina,N. |
| EPI529400 | HA | Australia | 2013-Nov-11 | A/SYDNEY/82/2013 | Clinical Virology Unit, CDIM | WHO Collaborating Centre for Reference and Research on Influenza | Deng,Y-M.; Iannello,P.; Spirason,N.; Jelley,L.; Lau,H.; Komadina,N. |
| EPI564992 | HA | Lao, People's Democratic Republic | 2014-Sep-15 | A/Laos/952/2014 | National Center for Laboratory and Epidemiology | Centers for Disease Control and Prevention |  |
| EPI564985 | NP | Lao, People's Democratic Republic | 2014-Sep-15 | A/Laos/952/2014 | National Center for Laboratory and Epidemiology | Centers for Disease Control and Prevention |  |
| EPI564986 | NS | Lao, People's Democratic Republic | 2014-Sep-15 | A/Laos/952/2014 | National Center for Laboratory and Epidemiology | Centers for Disease Control and Prevention |  |
| EPI564987 | MP | Lao, People's Democratic Republic | 2014-Sep-15 | A/Laos/952/2014 | National Center for Laboratory and Epidemiology | Centers for Disease Control and Prevention |  |
| EPI564988 | PA | Lao, People's Democratic Republic | 2014-Sep-15 | A/Laos/952/2014 | National Center for Laboratory and Epidemiology | Centers for Disease Control and Prevention |  |
| EPI564989 | PB2 | Lao, People's Democratic Republic | 2014-Sep-15 | A/Laos/952/2014 | National Center for Laboratory and Epidemiology | Centers for Disease Control and Prevention |  |
| EPI564990 | PB1 | Lao, People's Democratic Republic | 2014-Sep-15 | A/Laos/952/2014 | National Center for Laboratory and Epidemiology | Centers for Disease Control and Prevention |  |
| EPI564991 | NA | Lao, People's Democratic Republic | 2014-Sep-15 | A/Laos/952/2014 | National Center for Laboratory and Epidemiology | Centers for Disease Control and Prevention |  |
| EPI564993 | NP | Ethiopia | 2014-Nov-07 | A/Ethiopia/63/2014 | Ethiopian Health and Nutrition Research Institute (EHNRI) | Centers for Disease Control and Prevention |  |
| EPI564994 | NS | Ethiopia | 2014-Nov-07 | A/Ethiopia/63/2014 | Ethiopian Health and Nutrition Research Institute (EHNRI) | Centers for Disease Control and Prevention |  |
| EPI564995 | MP | Ethiopia | 2014-Nov-07 | A/Ethiopia/63/2014 | Ethiopian Health and Nutrition Research Institute (EHNRI) | Centers for Disease Control and Prevention |  |
| EPI564996 | PA | Ethiopia | 2014-Nov-07 | A/Ethiopia/63/2014 | Ethiopian Health and Nutrition Research Institute (EHNRI) | Centers for Disease Control and Prevention |  |
| EPI564997 | PB2 | Ethiopia | 2014-Nov-07 | A/Ethiopia/63/2014 | Ethiopian Health and Nutrition Research Institute (EHNRI) | Centers for Disease Control and Prevention |  |
| EPI564998 | PB1 | Ethiopia | 2014-Nov-07 | A/Ethiopia/63/2014 | Ethiopian Health and Nutrition Research Institute (EHNRI) | Centers for Disease Control and Prevention |  |
| EPI564999 | NA | Ethiopia | 2014-Nov-07 | A/Ethiopia/63/2014 | Ethiopian Health and Nutrition Research Institute (EHNRI) | Centers for Disease Control and Prevention |  |
| EPI565000 | HA | Ethiopia | 2014-Nov-07 | A/Ethiopia/63/2014 | Ethiopian Health and Nutrition Research Institute (EHNRI) | Centers for Disease Control and Prevention |  |
| EPI567686 | PB1 | Bahrain | 2014-Nov-18 | A/Bahrain/602/2014 | Ministry of Health Bahrain | Centers for Disease Control and Prevention |  |
| EPI567681 | NP | Bahrain | 2014-Nov-18 | A/Bahrain/602/2014 | Ministry of Health Bahrain | Centers for Disease Control and Prevention |  |
| EPI567682 | NS | Bahrain | 2014-Nov-18 | A/Bahrain/602/2014 | Ministry of Health Bahrain | Centers for Disease Control and Prevention |  |
| EPI567683 | MP | Bahrain | 2014-Nov-18 | A/Bahrain/602/2014 | Ministry of Health Bahrain | Centers for Disease Control and Prevention |  |
| EPI567684 | PA | Bahrain | 2014-Nov-18 | A/Bahrain/602/2014 | Ministry of Health Bahrain | Centers for Disease Control and Prevention |  |
| EPI567685 | PB2 | Bahrain | 2014-Nov-18 | A/Bahrain/602/2014 | Ministry of Health Bahrain | Centers for Disease Control and Prevention |  |
| EPI567687 | NA | Bahrain | 2014-Nov-18 | A/Bahrain/602/2014 | Ministry of Health Bahrain | Centers for Disease Control and Prevention | NA |
| EPI567688 | HA | Bahrain | 2014-Nov-18 | A/Bahrain/602/2014 | Ministry of Health Bahrain | Centers for Disease Control and Prevention | NA |
| EPI568042 | PA | Myanmar | 2014-Sep-05 | A/Myanmar/14M445/2014 |  | Niigata University | Saito, Reiko; Takemae, Nobuhiro; Saito, Takehiko; Shobugawa, Yugo; Kondo, Hiroki; Hibino, Akinobu; Yadanar, Kyaw; Yi Yi, Myint; Khin Yi, Oo; Htay Htay, Tin |
| EPI568043 | PB1 | Myanmar | 2014-Sep-05 | A/Myanmar/14M445/2014 |  | Niigata University | Saito, Reiko; Takemae, Nobuhiro; Saito, Takehiko; Shobugawa, Yugo; Kondo, Hiroki; Hibino, Akinobu; Yadanar, Kyaw; Yi Yi, Myint; Khin Yi, Oo; Htay Htay, Tin |
| EPI568048 | NS | Myanmar | 2014-Sep-05 | A/Myanmar/14M445/2014 |  | Niigata University | Saito, Reiko; Takemae, Nobuhiro; Saito, Takehiko; Shobugawa, Yugo; Kondo, Hiroki; Hibino, Akinobu; Yadanar, Kyaw; Yi Yi, Myint; Khin Yi, Oo; Htay Htay, Tin |
| EPI568044 | HA | Myanmar | 2014-Sep-05 | A/Myanmar/14M445/2014 |  | Niigata University | Saito, Reiko; Takemae, Nobuhiro; Saito, Takehiko; Shobugawa, Yugo; Kondo, Hiroki; Hibino, Akinobu; Yadanar, Kyaw; Yi Yi, Myint; Khin Yi, Oo; Htay Htay, Tin |
| EPI568046 | PB2 | Myanmar | 2014-Sep-05 | A/Myanmar/14M445/2014 |  | Niigata University | Saito, Reiko; Takemae, Nobuhiro; Saito, Takehiko; Shobugawa, Yugo; Kondo, Hiroki; Hibino, Akinobu; Yadanar, Kyaw; Yi Yi, Myint; Khin Yi, Oo; Htay Htay, Tin |
| EPI568045 | MP | Myanmar | 2014-Sep-05 | A/Myanmar/14M445/2014 |  | Niigata University | Saito, Reiko; Takemae, Nobuhiro; Saito, Takehiko; Shobugawa, Yugo; Kondo, Hiroki; Hibino, Akinobu; Yadanar, Kyaw; Yi Yi, Myint; Khin Yi, Oo; Htay Htay, Tin |
| EPI568047 | NA | Myanmar | 2014-Sep-05 | A/Myanmar/14M445/2014 |  | Niigata University | Saito, Reiko; Takemae, Nobuhiro; Saito, Takehiko; Shobugawa, Yugo; Kondo, Hiroki; Hibino, Akinobu; Yadanar, Kyaw; Yi Yi, Myint; Khin Yi, Oo; Htay Htay, Tin |
| EPI568049 | NP | Myanmar | 2014-Sep-05 | A/Myanmar/14M445/2014 |  | Niigata University | Saito, Reiko; Takemae, Nobuhiro; Saito, Takehiko; Shobugawa, Yugo; Kondo, Hiroki; Hibino, Akinobu; Yadanar, Kyaw; Yi Yi, Myint; Khin Yi, Oo; Htay Htay, Tin |
| EPI620332 | NP | Colombia | 2015-Feb-13 | A/Colombia/2537/2015 | Instituto Nacional de Salud de Columbia | Centers for Disease Control and Prevention |  |
| EPI620333 | NS | Colombia | 2015-Feb-13 | A/Colombia/2537/2015 | Instituto Nacional de Salud de Columbia | Centers for Disease Control and Prevention |  |
| EPI620334 | MP | Colombia | 2015-Feb-13 | A/Colombia/2537/2015 | Instituto Nacional de Salud de Columbia | Centers for Disease Control and Prevention |  |
| EPI620335 | PA | Colombia | 2015-Feb-13 | A/Colombia/2537/2015 | Instituto Nacional de Salud de Columbia | Centers for Disease Control and Prevention |  |
| EPI620338 | NA | Colombia | 2015-Feb-13 | A/Colombia/2537/2015 | Instituto Nacional de Salud de Columbia | Centers for Disease Control and Prevention |  |
| EPI620339 | HA | Colombia | 2015-Feb-13 | A/Colombia/2537/2015 | Instituto Nacional de Salud de Columbia | Centers for Disease Control and Prevention |  |
| EPI620336 | PB2 | Colombia | 2015-Feb-13 | A/Colombia/2537/2015 | Instituto Nacional de Salud de Columbia | Centers for Disease Control and Prevention |  |
| EPI620337 | PB1 | Colombia | 2015-Feb-13 | A/Colombia/2537/2015 | Instituto Nacional de Salud de Columbia | Centers for Disease Control and Prevention |  |
| EPI626069 | NP | United States | 2015-Mar-18 | A/California/80/2015 | California Department of Health Services | Centers for Disease Control and Prevention |  |
| EPI626070 | NS | United States | 2015-Mar-18 | A/California/80/2015 | California Department of Health Services | Centers for Disease Control and Prevention |  |
| EPI626071 | MP | United States | 2015-Mar-18 | A/California/80/2015 | California Department of Health Services | Centers for Disease Control and Prevention |  |
| EPI626072 | PA | United States | 2015-Mar-18 | A/California/80/2015 | California Department of Health Services | Centers for Disease Control and Prevention |  |
| EPI626075 | NA | United States | 2015-Mar-18 | A/California/80/2015 | California Department of Health Services | Centers for Disease Control and Prevention |  |
| EPI626076 | HA | United States | 2015-Mar-18 | A/California/80/2015 | California Department of Health Services | Centers for Disease Control and Prevention |  |
| EPI626073 | PB2 | United States | 2015-Mar-18 | A/California/80/2015 | California Department of Health Services | Centers for Disease Control and Prevention |  |
| EPI626074 | PB1 | United States | 2015-Mar-18 | A/California/80/2015 | California Department of Health Services | Centers for Disease Control and Prevention |  |
| EPI320002 | PB1 | United States | 2011-Feb-15 | A/California/17/2011 | Naval Health Research Center | Centers for Disease Control and Prevention |  |
| EPI316331 | NS | United States | 2011-Feb-15 | A/California/17/2011 | Naval Health Research Center | Centers for Disease Control and Prevention |  |
| EPI316330 | NP | United States | 2011-Feb-15 | A/California/17/2011 | Naval Health Research Center | Centers for Disease Control and Prevention |  |
| EPI316332 | PA | United States | 2011-Feb-15 | A/California/17/2011 | Naval Health Research Center | Centers for Disease Control and Prevention |  |
| EPI316333 | PB2 | United States | 2011-Feb-15 | A/California/17/2011 | Naval Health Research Center | Centers for Disease Control and Prevention |  |
| EPI316334 | NA | United States | 2011-Feb-15 | A/California/17/2011 | Naval Health Research Center | Centers for Disease Control and Prevention |  |
| EPI316335 | HA | United States | 2011-Feb-15 | A/California/17/2011 | Naval Health Research Center | Centers for Disease Control and Prevention |  |
| EPI320001 | MP | United States | 2011-Feb-15 | A/California/17/2011 | Naval Health Research Center | Centers for Disease Control and Prevention |  |
| EPI466830 | NP | Bolivia, Plurinationial State of | 2013-Jun-08 | A/Bolivia/559/2013 | Instituto Nacional de Laboratoriosde Salud (INLASA) | Centers for Disease Control and Prevention |  |
| EPI466831 | NS | Bolivia, Plurinationial State of | 2013-Jun-08 | A/Bolivia/559/2013 | Instituto Nacional de Laboratoriosde Salud (INLASA) | Centers for Disease Control and Prevention |  |
| EPI466832 | MP | Bolivia, Plurinationial State of | 2013-Jun-08 | A/Bolivia/559/2013 | Instituto Nacional de Laboratoriosde Salud (INLASA) | Centers for Disease Control and Prevention |  |
| EPI466833 | PA | Bolivia, Plurinationial State of | 2013-Jun-08 | A/Bolivia/559/2013 | Instituto Nacional de Laboratoriosde Salud (INLASA) | Centers for Disease Control and Prevention |  |
| EPI466834 | PB2 | Bolivia, Plurinationial State of | 2013-Jun-08 | A/Bolivia/559/2013 | Instituto Nacional de Laboratoriosde Salud (INLASA) | Centers for Disease Control and Prevention |  |
| EPI466835 | PB1 | Bolivia, Plurinationial State of | 2013-Jun-08 | A/Bolivia/559/2013 | Instituto Nacional de Laboratoriosde Salud (INLASA) | Centers for Disease Control and Prevention |  |
| EPI466836 | NA | Bolivia, Plurinationial State of | 2013-Jun-08 | A/Bolivia/559/2013 | Instituto Nacional de Laboratoriosde Salud (INLASA) | Centers for Disease Control and Prevention | NA |
| EPI466837 | HA | Bolivia, Plurinationial State of | 2013-Jun-08 | A/Bolivia/559/2013 | Instituto Nacional de Laboratoriosde Salud (INLASA) | Centers for Disease Control and Prevention | NA |
| EPI468848 | PA | India | 2013-Feb-18 | A/India/P132194/2013 |  | Other Database Import | Potdar,V.A.; Dakhave,M.R.; Patil,K.N.; Kadam,A.A.; Mullick,J.; Chadha,M.S. |
| EPI468829 | PB2 | India | 2013-Feb-18 | A/India/P132194/2013 |  | Other Database Import | Potdar,V.A.; Dakhave,M.R.; Patil,K.N.; Kadam,A.A.; Mullick,J.; Chadha,M.S. |
| EPI468830 | PB1 | India | 2013-Feb-18 | A/India/P132194/2013 |  | Other Database Import | Potdar,V.A.; Dakhave,M.R.; Patil,K.N.; Kadam,A.A.; Mullick,J.; Chadha,M.S. |
| EPI468832 | HA | India | 2013-Feb-18 | A/India/P132194/2013 |  | Other Database Import | Potdar,V.A.; Dakhave,M.R.; Patil,K.N.; Kadam,A.A.; Mullick,J.; Chadha,M.S. |
| EPI468833 | NP | India | 2013-Feb-18 | A/India/P132194/2013 |  | Other Database Import | Potdar,V.A.; Dakhave,M.R.; Patil,K.N.; Kadam,A.A.; Mullick,J.; Chadha,M.S. |
| EPI468834 | NA | India | 2013-Feb-18 | A/India/P132194/2013 |  | Other Database Import | Potdar,V.A.; Dakhave,M.R.; Patil,K.N.; Kadam,A.A.; Mullick,J.; Chadha,M.S. |
| EPI468835 | MP | India | 2013-Feb-18 | A/India/P132194/2013 |  | Other Database Import | Potdar,V.A.; Dakhave,M.R.; Patil,K.N.; Kadam,A.A.; Mullick,J.; Chadha,M.S. |
| EPI468836 | NS | India | 2013-Feb-18 | A/India/P132194/2013 |  | Other Database Import | Potdar,V.A.; Dakhave,M.R.; Patil,K.N.; Kadam,A.A.; Mullick,J.; Chadha,M.S. |
| EPI498106 | PB2 | India | 2013-Jan-27 | A/India/P131027/2013 |  | Other Database Import | Potdar,V.A.; Kadam,A.A.; Dakhave,M.R.; Chadha,M.S. |
| EPI498107 | PB1 | India | 2013-Jan-27 | A/India/P131027/2013 |  | Other Database Import | Potdar,V.A.; Kadam,A.A.; Dakhave,M.R.; Chadha,M.S. |
| EPI498109 | HA | India | 2013-Jan-27 | A/India/P131027/2013 |  | Other Database Import | Potdar,V.A.; Kadam,A.A.; Dakhave,M.R.; Chadha,M.S. |
| EPI498110 | NP | India | 2013-Jan-27 | A/India/P131027/2013 |  | Other Database Import | Potdar,V.A.; Kadam,A.A.; Dakhave,M.R.; Chadha,M.S. |
| EPI498111 | NA | India | 2013-Jan-27 | A/India/P131027/2013 |  | Other Database Import | Potdar,V.A.; Kadam,A.A.; Dakhave,M.R.; Chadha,M.S. |
| EPI498112 | MP | India | 2013-Jan-27 | A/India/P131027/2013 |  | Other Database Import | Potdar,V.A.; Kadam,A.A.; Dakhave,M.R.; Chadha,M.S. |
| EPI498113 | NS | India | 2013-Jan-27 | A/India/P131027/2013 |  | Other Database Import | Potdar,V.A.; Kadam,A.A.; Dakhave,M.R.; Chadha,M.S. |
| EPI498116 | PA | India | 2013-Jan-27 | A/India/P131027/2013 |  | Other Database Import | Potdar,V.A.; Kadam,A.A.; Dakhave,M.R.; Chadha,M.S. |
| EPI503390 | NP | United States | 2014-Jan-03 | A/Florida/01/2014 | Florida Department of Health-Jacksonville | Centers for Disease Control and Prevention |  |
| EPI503391 | NS | United States | 2014-Jan-03 | A/Florida/01/2014 | Florida Department of Health-Jacksonville | Centers for Disease Control and Prevention |  |
| EPI503392 | MP | United States | 2014-Jan-03 | A/Florida/01/2014 | Florida Department of Health-Jacksonville | Centers for Disease Control and Prevention |  |
| EPI503393 | PA | United States | 2014-Jan-03 | A/Florida/01/2014 | Florida Department of Health-Jacksonville | Centers for Disease Control and Prevention |  |
| EPI503396 | NA | United States | 2014-Jan-03 | A/Florida/01/2014 | Florida Department of Health-Jacksonville | Centers for Disease Control and Prevention |  |
| EPI503397 | HA | United States | 2014-Jan-03 | A/Florida/01/2014 | Florida Department of Health-Jacksonville | Centers for Disease Control and Prevention |  |
| EPI503394 | PB2 | United States | 2014-Jan-03 | A/Florida/01/2014 | Florida Department of Health-Jacksonville | Centers for Disease Control and Prevention |  |
| EPI503395 | PB1 | United States | 2014-Jan-03 | A/Florida/01/2014 | Florida Department of Health-Jacksonville | Centers for Disease Control and Prevention |  |
| EPI511893 | NP | United States | 2014-Feb-08 | A/Washington/05/2014 | Spokane Regional Health District | Centers for Disease Control and Prevention |  |
| EPI511894 | NS | United States | 2014-Feb-08 | A/Washington/05/2014 | Spokane Regional Health District | Centers for Disease Control and Prevention |  |
| EPI511895 | PA | United States | 2014-Feb-08 | A/Washington/05/2014 | Spokane Regional Health District | Centers for Disease Control and Prevention |  |
| EPI511896 | PB2 | United States | 2014-Feb-08 | A/Washington/05/2014 | Spokane Regional Health District | Centers for Disease Control and Prevention |  |
| EPI511897 | PB1 | United States | 2014-Feb-08 | A/Washington/05/2014 | Spokane Regional Health District | Centers for Disease Control and Prevention |  |
| EPI511898 | HA | United States | 2014-Feb-08 | A/Washington/05/2014 | Spokane Regional Health District | Centers for Disease Control and Prevention |  |
| EPI515739 | MP | United States | 2014-Feb-08 | A/Washington/05/2014 | Spokane Regional Health District | Centers for Disease Control and Prevention | NA |
| EPI516290 | NA | United States | 2014-Feb-08 | A/Washington/05/2014 | Spokane Regional Health District | Centers for Disease Control and Prevention | NA |
| EPI562718 | MP | Malaysia | 2014-Mar-02 | A/Malaysia/11/2014 | Institut Penyelidikan Perubatan | WHO Collaborating Centre for Reference and Research on Influenza | Deng,Y-M.; Iannello,P.; Spirason,N; Jelley,L.;Lau,H.; Komadina,N. |
| EPI562720 | HA | Malaysia | 2014-Mar-02 | A/Malaysia/11/2014 | Institut Penyelidikan Perubatan | WHO Collaborating Centre for Reference and Research on Influenza | Deng,Y-M.; Iannello,P.; Spirason,N; Jelley,L.;Lau,H.; Komadina,N. |
| EPI562719 | NA | Malaysia | 2014-Mar-02 | A/Malaysia/11/2014 | Institut Penyelidikan Perubatan | WHO Collaborating Centre for Reference and Research on Influenza | Deng,Y-M.; Iannello,P.; Spirason,N; Jelley,L.;Lau,H.; Komadina,N. |
| EPI561955 | NP | Malaysia | 2014-Mar-02 | A/Malaysia/11/2014 | Institut Penyelidikan Perubatan | WHO Collaborating Centre for Reference and Research on Influenza | Deng,Y-M.; Iannello,P.; Spirason,N; Jelley,L.;Lau,H.; Komadina,N. |
| EPI561956 | NS | Malaysia | 2014-Mar-02 | A/Malaysia/11/2014 | Institut Penyelidikan Perubatan | WHO Collaborating Centre for Reference and Research on Influenza | Deng,Y-M.; Iannello,P.; Spirason,N; Jelley,L.;Lau,H.; Komadina,N. |
| EPI561957 | PA | Malaysia | 2014-Mar-02 | A/Malaysia/11/2014 | Institut Penyelidikan Perubatan | WHO Collaborating Centre for Reference and Research on Influenza | Deng,Y-M.; Iannello,P.; Spirason,N; Jelley,L.;Lau,H.; Komadina,N. |
| EPI561958 | PB2 | Malaysia | 2014-Mar-02 | A/Malaysia/11/2014 | Institut Penyelidikan Perubatan | WHO Collaborating Centre for Reference and Research on Influenza | Deng,Y-M.; Iannello,P.; Spirason,N; Jelley,L.;Lau,H.; Komadina,N. |
| EPI561959 | PB1 | Malaysia | 2014-Mar-02 | A/Malaysia/11/2014 | Institut Penyelidikan Perubatan | WHO Collaborating Centre for Reference and Research on Influenza | Deng,Y-M.; Iannello,P.; Spirason,N; Jelley,L.;Lau,H.; Komadina,N. |
| EPI598943 | MP | Thailand | 2010-Sep-28 | A/Bangkok/SIMI511/2010 |  | Other Database Import | Horthongkham,N.; Athipanyasilp,N.; Kaewnapan,B.; Sirirtantikorn,S.; Kantakamalakul,W.; Sutthent,R. |
| EPI598921 | NP | Thailand | 2010-Sep-28 | A/Bangkok/SIMI511/2010 |  | Other Database Import | Horthongkham,N.; Athipanyasilp,N.; Kaewnapan,B.; Sirirtantikorn,S.; Kantakamalakul,W.; Sutthent,R. |
| EPI598932 | NS | Thailand | 2010-Sep-28 | A/Bangkok/SIMI511/2010 |  | Other Database Import | Horthongkham,N.; Athipanyasilp,N.; Kaewnapan,B.; Sirirtantikorn,S.; Kantakamalakul,W.; Sutthent,R. |
| EPI598877 | NA | Thailand | 2010-Sep-28 | A/Bangkok/SIMI511/2010 |  | Other Database Import | Horthongkham,N.; Athipanyasilp,N.; Kaewnapan,B.; Sirirtantikorn,S.; Kantakamalakul,W.; Sutthent,R. |
| EPI598306 | PA | Thailand | 2010-Sep-28 | A/Bangkok/SIMI511/2010 |  | Other Database Import | Horthongkham,N.; Athipanyasilp,N.; Kaewnapan,B.; Sirirtantikorn,S.; Kantakamalakul,W.; Sutthent,R. |
| EPI598888 | HA | Thailand | 2010-Sep-28 | A/Bangkok/SIMI511/2010 |  | Other Database Import | Horthongkham,N.; Athipanyasilp,N.; Kaewnapan,B.; Sirirtantikorn,S.; Kantakamalakul,W.; Sutthent,R. |
| EPI598899 | PB1 | Thailand | 2010-Sep-28 | A/Bangkok/SIMI511/2010 |  | Other Database Import | Horthongkham,N.; Athipanyasilp,N.; Kaewnapan,B.; Sirirtantikorn,S.; Kantakamalakul,W.; Sutthent,R. |
| EPI598910 | PB2 | Thailand | 2010-Sep-28 | A/Bangkok/SIMI511/2010 |  | Other Database Import | Horthongkham,N.; Athipanyasilp,N.; Kaewnapan,B.; Sirirtantikorn,S.; Kantakamalakul,W.; Sutthent,R. |
| EPI677947 | NP | United Kingdom | 2014-Feb-25 | A/England/41420357/2014 | Microbiology Services Colindale, Public Health England | Microbiology Services Colindale, Public Health England | Galiano,M. |
| EPI677948 | NS | United Kingdom | 2014-Feb-25 | A/England/41420357/2014 | Microbiology Services Colindale, Public Health England | Microbiology Services Colindale, Public Health England | Galiano,M. |
| EPI677949 | MP | United Kingdom | 2014-Feb-25 | A/England/41420357/2014 | Microbiology Services Colindale, Public Health England | Microbiology Services Colindale, Public Health England | Galiano,M. |
| EPI677950 | PA | United Kingdom | 2014-Feb-25 | A/England/41420357/2014 | Microbiology Services Colindale, Public Health England | Microbiology Services Colindale, Public Health England | Galiano,M. |
| EPI677951 | PB2 | United Kingdom | 2014-Feb-25 | A/England/41420357/2014 | Microbiology Services Colindale, Public Health England | Microbiology Services Colindale, Public Health England | Galiano,M. |
| EPI677952 | PB1 | United Kingdom | 2014-Feb-25 | A/England/41420357/2014 | Microbiology Services Colindale, Public Health England | Microbiology Services Colindale, Public Health England | Galiano,M. |
| EPI677953 | NA | United Kingdom | 2014-Feb-25 | A/England/41420357/2014 | Microbiology Services Colindale, Public Health England | Microbiology Services Colindale, Public Health England | Galiano,M. |
| EPI677954 | HA | United Kingdom | 2014-Feb-25 | A/England/41420357/2014 | Microbiology Services Colindale, Public Health England | Microbiology Services Colindale, Public Health England | Galiano,M. |
| EPI319573 | PA | Singapore | 2010-Dec-15 | A/Singapore/640/2010 | Ministry of Health, Singapore | WHO Collaborating Centre for Reference and Research on Influenza | NA |
| EPI319574 | PB2 | Singapore | 2010-Dec-15 | A/Singapore/640/2010 | Ministry of Health, Singapore | WHO Collaborating Centre for Reference and Research on Influenza | NA |
| EPI319571 | NS | Singapore | 2010-Dec-15 | A/Singapore/640/2010 | Ministry of Health, Singapore | WHO Collaborating Centre for Reference and Research on Influenza | NA |
| EPI319575 | PB1 | Singapore | 2010-Dec-15 | A/Singapore/640/2010 | Ministry of Health, Singapore | WHO Collaborating Centre for Reference and Research on Influenza | NA |
| EPI319578 | HA | Singapore | 2010-Dec-15 | A/Singapore/640/2010 | Ministry of Health, Singapore | WHO Collaborating Centre for Reference and Research on Influenza | NA |
| EPI319576 | MP | Singapore | 2010-Dec-15 | A/Singapore/640/2010 | Ministry of Health, Singapore | WHO Collaborating Centre for Reference and Research on Influenza | NA |
| EPI319572 | NP | Singapore | 2010-Dec-15 | A/Singapore/640/2010 | Ministry of Health, Singapore | WHO Collaborating Centre for Reference and Research on Influenza | NA |
| EPI319577 | NA | Singapore | 2010-Dec-15 | A/Singapore/640/2010 | Ministry of Health, Singapore | WHO Collaborating Centre for Reference and Research on Influenza | NA |
| EPI349213 | MP | Cambodia | 2011-Jul-18 | A/CAMBODIA/11/2011 | Institute Pasteur du Cambodia | WHO Collaborating Centre for Reference and Research on Influenza | Deng,Y-M.; Iannello,P.; Caldwell,N.; Leang,S-K; Komadina,N. |
| EPI349214 | NA | Cambodia | 2011-Jul-18 | A/CAMBODIA/11/2011 | Institute Pasteur du Cambodia | WHO Collaborating Centre for Reference and Research on Influenza | Deng,Y-M.; Iannello,P.; Caldwell,N.; Leang,S-K; Komadina,N. |
| EPI349215 | HA | Cambodia | 2011-Jul-18 | A/CAMBODIA/11/2011 | Institute Pasteur du Cambodia | WHO Collaborating Centre for Reference and Research on Influenza | Deng,Y-M.; Iannello,P.; Caldwell,N.; Leang,S-K; Komadina,N. |
| EPI370269 | NP | Cambodia | 2011-Jul-18 | A/CAMBODIA/11/2011 | Institute Pasteur du Cambodia | WHO Collaborating Centre for Reference and Research on Influenza | Deng,Y-M.; Iannello,P.; Caldwell,N.; Leang,S-K; Komadina,N. |
| EPI370270 | NS | Cambodia | 2011-Jul-18 | A/CAMBODIA/11/2011 | Institute Pasteur du Cambodia | WHO Collaborating Centre for Reference and Research on Influenza | Deng,Y-M.; Iannello,P.; Caldwell,N.; Leang,S-K; Komadina,N. |
| EPI370271 | PA | Cambodia | 2011-Jul-18 | A/CAMBODIA/11/2011 | Institute Pasteur du Cambodia | WHO Collaborating Centre for Reference and Research on Influenza | Deng,Y-M.; Iannello,P.; Caldwell,N.; Leang,S-K; Komadina,N. |
| EPI370272 | PB2 | Cambodia | 2011-Jul-18 | A/CAMBODIA/11/2011 | Institute Pasteur du Cambodia | WHO Collaborating Centre for Reference and Research on Influenza | Deng,Y-M.; Iannello,P.; Caldwell,N.; Leang,S-K; Komadina,N. |
| EPI370273 | PB1 | Cambodia | 2011-Jul-18 | A/CAMBODIA/11/2011 | Institute Pasteur du Cambodia | WHO Collaborating Centre for Reference and Research on Influenza | Deng,Y-M.; Iannello,P.; Caldwell,N.; Leang,S-K; Komadina,N. |
| EPI394204 | MP | Malaysia | 2012-Jan-01 | A/MALAYSIA/192/2012 | Institut Penyelidikan Perubatan | WHO Collaborating Centre for Reference and Research on Influenza | Deng,Y-M; Iannello,P; Caldwell,N; Jelley,L; Komadina,N |
| EPI394205 | NA | Malaysia | 2012-Jan-01 | A/MALAYSIA/192/2012 | Institut Penyelidikan Perubatan | WHO Collaborating Centre for Reference and Research on Influenza | Deng,Y-M; Iannello,P; Caldwell,N; Jelley,L; Komadina,N |
| EPI394206 | HA | Malaysia | 2012-Jan-01 | A/MALAYSIA/192/2012 | Institut Penyelidikan Perubatan | WHO Collaborating Centre for Reference and Research on Influenza | Deng,Y-M; Iannello,P; Caldwell,N; Jelley,L; Komadina,N |
| EPI450418 | NP | Malaysia | 2012-Jan-01 | A/MALAYSIA/192/2012 | Institut Penyelidikan Perubatan | WHO Collaborating Centre for Reference and Research on Influenza | Deng,Y-M; Iannello,P; Caldwell,N; Jelley,L; Komadina,N |
| EPI450419 | NS | Malaysia | 2012-Jan-01 | A/MALAYSIA/192/2012 | Institut Penyelidikan Perubatan | WHO Collaborating Centre for Reference and Research on Influenza | Deng,Y-M; Iannello,P; Caldwell,N; Jelley,L; Komadina,N |
| EPI450420 | PA | Malaysia | 2012-Jan-01 | A/MALAYSIA/192/2012 | Institut Penyelidikan Perubatan | WHO Collaborating Centre for Reference and Research on Influenza | Deng,Y-M; Iannello,P; Caldwell,N; Jelley,L; Komadina,N |
| EPI450421 | PB2 | Malaysia | 2012-Jan-01 | A/MALAYSIA/192/2012 | Institut Penyelidikan Perubatan | WHO Collaborating Centre for Reference and Research on Influenza | Deng,Y-M; Iannello,P; Caldwell,N; Jelley,L; Komadina,N |
| EPI450422 | PB1 | Malaysia | 2012-Jan-01 | A/MALAYSIA/192/2012 | Institut Penyelidikan Perubatan | WHO Collaborating Centre for Reference and Research on Influenza | Deng,Y-M; Iannello,P; Caldwell,N; Jelley,L; Komadina,N |
| EPI394290 | MP | Singapore | 2012-Jun-22 | A/SINGAPORE/12/2012 | Ministry of Health, Singapore | WHO Collaborating Centre for Reference and Research on Influenza | Deng,Y-M; Iannello,P; Caldwell,N; Jelley,L; Komadina,N |
| EPI394291 | NA | Singapore | 2012-Jun-22 | A/SINGAPORE/12/2012 | Ministry of Health, Singapore | WHO Collaborating Centre for Reference and Research on Influenza | Deng,Y-M; Iannello,P; Caldwell,N; Jelley,L; Komadina,N |
| EPI394292 | HA | Singapore | 2012-Jun-22 | A/SINGAPORE/12/2012 | Ministry of Health, Singapore | WHO Collaborating Centre for Reference and Research on Influenza | Deng,Y-M; Iannello,P; Caldwell,N; Jelley,L; Komadina,N |
| EPI450423 | NP | Singapore | 2012-Jun-22 | A/SINGAPORE/12/2012 | Ministry of Health, Singapore | WHO Collaborating Centre for Reference and Research on Influenza | Deng,Y-M; Iannello,P; Caldwell,N; Jelley,L; Komadina,N |
| EPI450424 | NS | Singapore | 2012-Jun-22 | A/SINGAPORE/12/2012 | Ministry of Health, Singapore | WHO Collaborating Centre for Reference and Research on Influenza | Deng,Y-M; Iannello,P; Caldwell,N; Jelley,L; Komadina,N |
| EPI450425 | PA | Singapore | 2012-Jun-22 | A/SINGAPORE/12/2012 | Ministry of Health, Singapore | WHO Collaborating Centre for Reference and Research on Influenza | Deng,Y-M; Iannello,P; Caldwell,N; Jelley,L; Komadina,N |
| EPI450426 | PB2 | Singapore | 2012-Jun-22 | A/SINGAPORE/12/2012 | Ministry of Health, Singapore | WHO Collaborating Centre for Reference and Research on Influenza | Deng,Y-M; Iannello,P; Caldwell,N; Jelley,L; Komadina,N |
| EPI450427 | PB1 | Singapore | 2012-Jun-22 | A/SINGAPORE/12/2012 | Ministry of Health, Singapore | WHO Collaborating Centre for Reference and Research on Influenza | Deng,Y-M; Iannello,P; Caldwell,N; Jelley,L; Komadina,N |
| EPI460817 | NP | Peru | 2013-Apr-19 | A/Peru/106/2013 | US NAMRU-6 | Centers for Disease Control and Prevention | NA |
| EPI460818 | MP | Peru | 2013-Apr-19 | A/Peru/106/2013 | US NAMRU-6 | Centers for Disease Control and Prevention | NA |
| EPI460819 | PA | Peru | 2013-Apr-19 | A/Peru/106/2013 | US NAMRU-6 | Centers for Disease Control and Prevention | NA |
| EPI460820 | PB2 | Peru | 2013-Apr-19 | A/Peru/106/2013 | US NAMRU-6 | Centers for Disease Control and Prevention | NA |
| EPI460821 | PB1 | Peru | 2013-Apr-19 | A/Peru/106/2013 | US NAMRU-6 | Centers for Disease Control and Prevention | NA |
| EPI460822 | NA | Peru | 2013-Apr-19 | A/Peru/106/2013 | US NAMRU-6 | Centers for Disease Control and Prevention | NA |
| EPI460823 | HA | Peru | 2013-Apr-19 | A/Peru/106/2013 | US NAMRU-6 | Centers for Disease Control and Prevention | NA |
| EPI464411 | NS | Peru | 2013-Apr-19 | A/Peru/106/2013 | US NAMRU-6 | Centers for Disease Control and Prevention | NA |
| EPI468886 | PB2 | India | 2011-Sep-23 | A/India/P1114854/2011 |  | Other Database Import | Potdar,V.A.; Dakhave,M.R.; Patil,K.N.; Kadam,A.A.; Mullick,J.; Chadha,M.S. |
| EPI468887 | PB1 | India | 2011-Sep-23 | A/India/P1114854/2011 |  | Other Database Import | Potdar,V.A.; Dakhave,M.R.; Patil,K.N.; Kadam,A.A.; Mullick,J.; Chadha,M.S. |
| EPI468889 | HA | India | 2011-Sep-23 | A/India/P1114854/2011 |  | Other Database Import | Potdar,V.A.; Dakhave,M.R.; Patil,K.N.; Kadam,A.A.; Mullick,J.; Chadha,M.S. |
| EPI468890 | NP | India | 2011-Sep-23 | A/India/P1114854/2011 |  | Other Database Import | Potdar,V.A.; Dakhave,M.R.; Patil,K.N.; Kadam,A.A.; Mullick,J.; Chadha,M.S. |
| EPI468891 | NA | India | 2011-Sep-23 | A/India/P1114854/2011 |  | Other Database Import | Potdar,V.A.; Dakhave,M.R.; Patil,K.N.; Kadam,A.A.; Mullick,J.; Chadha,M.S. |
| EPI468892 | MP | India | 2011-Sep-23 | A/India/P1114854/2011 |  | Other Database Import | Potdar,V.A.; Dakhave,M.R.; Patil,K.N.; Kadam,A.A.; Mullick,J.; Chadha,M.S. |
| EPI468893 | NS | India | 2011-Sep-23 | A/India/P1114854/2011 |  | Other Database Import | Potdar,V.A.; Dakhave,M.R.; Patil,K.N.; Kadam,A.A.; Mullick,J.; Chadha,M.S. |
| EPI468934 | PA | India | 2011-Sep-23 | A/India/P1114854/2011 |  | Other Database Import | Potdar,V.A.; Dakhave,M.R.; Patil,K.N.; Kadam,A.A.; Mullick,J.; Chadha,M.S. |
| EPI580157 | NP | Uganda | 2014-Jul-07 | A/Uganda/1625/2014 | Uganda Virus Research Institute (UVRI), National Influenza Center | Centers for Disease Control and Prevention | NA |
| EPI580158 | NS | Uganda | 2014-Jul-07 | A/Uganda/1625/2014 | Uganda Virus Research Institute (UVRI), National Influenza Center | Centers for Disease Control and Prevention | NA |
| EPI580159 | MP | Uganda | 2014-Jul-07 | A/Uganda/1625/2014 | Uganda Virus Research Institute (UVRI), National Influenza Center | Centers for Disease Control and Prevention | NA |
| EPI580160 | PA | Uganda | 2014-Jul-07 | A/Uganda/1625/2014 | Uganda Virus Research Institute (UVRI), National Influenza Center | Centers for Disease Control and Prevention | NA |
| EPI580161 | PB2 | Uganda | 2014-Jul-07 | A/Uganda/1625/2014 | Uganda Virus Research Institute (UVRI), National Influenza Center | Centers for Disease Control and Prevention | NA |
| EPI580162 | PB1 | Uganda | 2014-Jul-07 | A/Uganda/1625/2014 | Uganda Virus Research Institute (UVRI), National Influenza Center | Centers for Disease Control and Prevention | NA |
| EPI580163 | NA | Uganda | 2014-Jul-07 | A/Uganda/1625/2014 | Uganda Virus Research Institute (UVRI), National Influenza Center | Centers for Disease Control and Prevention | NA |
| EPI580164 | HA | Uganda | 2014-Jul-07 | A/Uganda/1625/2014 | Uganda Virus Research Institute (UVRI), National Influenza Center | Centers for Disease Control and Prevention | NA |
| EPI277233 | PB2 | Thailand | 2010-Feb-23 | A/Bangkok/INS424/2010 |  | Other Database Import | The NIAID Influenza Genome Sequencing Consortium |
| EPI277234 | PB1 | Thailand | 2010-Feb-23 | A/Bangkok/INS424/2010 |  | Other Database Import | The NIAID Influenza Genome Sequencing Consortium |
| EPI277235 | PA | Thailand | 2010-Feb-23 | A/Bangkok/INS424/2010 |  | Other Database Import | The NIAID Influenza Genome Sequencing Consortium |
| EPI277236 | HA | Thailand | 2010-Feb-23 | A/Bangkok/INS424/2010 |  | Other Database Import | The NIAID Influenza Genome Sequencing Consortium |
| EPI277237 | NP | Thailand | 2010-Feb-23 | A/Bangkok/INS424/2010 |  | Other Database Import | The NIAID Influenza Genome Sequencing Consortium |
| EPI277238 | NA | Thailand | 2010-Feb-23 | A/Bangkok/INS424/2010 |  | Other Database Import | The NIAID Influenza Genome Sequencing Consortium |
| EPI277239 | MP | Thailand | 2010-Feb-23 | A/Bangkok/INS424/2010 |  | Other Database Import | The NIAID Influenza Genome Sequencing Consortium |
| EPI277240 | NS | Thailand | 2010-Feb-23 | A/Bangkok/INS424/2010 |  | Other Database Import | The NIAID Influenza Genome Sequencing Consortium |
| EPI353907 | HA | Mexico | 2011-Dec-12 | A/Mexico/3723/2011 | Laboratorio de Virus Respiratorio | Centers for Disease Control and Prevention | NA |
| EPI353431 | NS | Mexico | 2011-Dec-12 | A/Mexico/3723/2011 | Laboratorio de Virus Respiratorio | Centers for Disease Control and Prevention |  |
| EPI353432 | MP | Mexico | 2011-Dec-12 | A/Mexico/3723/2011 | Laboratorio de Virus Respiratorio | Centers for Disease Control and Prevention | NA |
| EPI353433 | PA | Mexico | 2011-Dec-12 | A/Mexico/3723/2011 | Laboratorio de Virus Respiratorio | Centers for Disease Control and Prevention | NA |
| EPI353434 | PB2 | Mexico | 2011-Dec-12 | A/Mexico/3723/2011 | Laboratorio de Virus Respiratorio | Centers for Disease Control and Prevention | NA |
| EPI353435 | PB1 | Mexico | 2011-Dec-12 | A/Mexico/3723/2011 | Laboratorio de Virus Respiratorio | Centers for Disease Control and Prevention | NA |
| EPI353436 | NA | Mexico | 2011-Dec-12 | A/Mexico/3723/2011 | Laboratorio de Virus Respiratorio | Centers for Disease Control and Prevention | NA |
| EPI353430 | NP | Mexico | 2011-Dec-12 | A/Mexico/3723/2011 | Laboratorio de Virus Respiratorio | Centers for Disease Control and Prevention | NA |
| EPI357518 | MP | Kenya | 2011-Nov-10 | A/Kenya/196/2011 | US Army Medical Research Unit - Kenya (USAMRU-K), GEIS Human Influenza Program | USAMRU-K | Bulimo,W.D; Achilla,R.A; Majanja,J.M; Wadegu,M.O; Osuna,F.A; Mukunzi,S.O; Mwangi, J.K; Mwangi,J.W; Opot, B.H; Kibet, K.M; Muthoni, J.N; Njiri,J.O.; Ocholla,S; Wurapa,K.E |
| EPI357519 | NS | Kenya | 2011-Nov-10 | A/Kenya/196/2011 | US Army Medical Research Unit - Kenya (USAMRU-K), GEIS Human Influenza Program | USAMRU-K | Bulimo,W.D; Achilla,R.A; Majanja,J.M; Wadegu,M.O; Osuna,F.A; Mukunzi,S.O; Mwangi, J.K; Mwangi,J.W; Opot, B.H; Kibet, K.M; Muthoni, J.N; Njiri,J.O.; Ocholla,S; Wurapa,K.E |
| EPI357515 | PB1 | Kenya | 2011-Nov-10 | A/Kenya/196/2011 | US Army Medical Research Unit - Kenya (USAMRU-K), GEIS Human Influenza Program | USAMRU-K | Bulimo,W.D; Achilla,R.A; Majanja,J.M; Wadegu,M.O; Osuna,F.A; Mukunzi,S.O; Mwangi, J.K; Mwangi,J.W; Opot, B.H; Kibet, K.M; Muthoni, J.N; Njiri,J.O.; Ocholla,S; Wurapa,K.E |
| EPI357517 | PA | Kenya | 2011-Nov-10 | A/Kenya/196/2011 | US Army Medical Research Unit - Kenya (USAMRU-K), GEIS Human Influenza Program | USAMRU-K | Bulimo,W.D; Achilla,R.A; Majanja,J.M; Wadegu,M.O; Osuna,F.A; Mukunzi,S.O; Mwangi, J.K; Mwangi,J.W; Opot, B.H; Kibet, K.M; Muthoni, J.N; Njiri,J.O.; Ocholla,S; Wurapa,K.E |
| EPI357516 | PB2 | Kenya | 2011-Nov-10 | A/Kenya/196/2011 | US Army Medical Research Unit - Kenya (USAMRU-K), GEIS Human Influenza Program | USAMRU-K | Bulimo,W.D; Achilla,R.A; Majanja,J.M; Wadegu,M.O; Osuna,F.A; Mukunzi,S.O; Mwangi, J.K; Mwangi,J.W; Opot, B.H; Kibet, K.M; Muthoni, J.N; Njiri,J.O.; Ocholla,S; Wurapa,K.E |
| EPI357514 | NA | Kenya | 2011-Nov-10 | A/Kenya/196/2011 | US Army Medical Research Unit - Kenya (USAMRU-K), GEIS Human Influenza Program | USAMRU-K | Bulimo,W.D; Achilla,R.A; Majanja,J.M; Wadegu,M.O; Osuna,F.A; Mukunzi,S.O; Mwangi, J.K; Mwangi,J.W; Opot, B.H; Kibet, K.M; Muthoni, J.N; Njiri,J.O.; Ocholla,S; Wurapa,K.E |
| EPI357520 | NP | Kenya | 2011-Nov-10 | A/Kenya/196/2011 | US Army Medical Research Unit - Kenya (USAMRU-K), GEIS Human Influenza Program | USAMRU-K | Bulimo,W.D; Achilla,R.A; Majanja,J.M; Wadegu,M.O; Osuna,F.A; Mukunzi,S.O; Mwangi, J.K; Mwangi,J.W; Opot, B.H; Kibet, K.M; Muthoni, J.N; Njiri,J.O.; Ocholla,S; Wurapa,K.E |
| EPI357513 | HA | Kenya | 2011-Nov-10 | A/Kenya/196/2011 | US Army Medical Research Unit - Kenya (USAMRU-K), GEIS Human Influenza Program | USAMRU-K | Bulimo,W.D; Achilla,R.A; Majanja,J.M; Wadegu,M.O; Osuna,F.A; Mukunzi,S.O; Mwangi, J.K; Mwangi,J.W; Opot, B.H; Kibet, K.M; Muthoni, J.N; Njiri,J.O.; Ocholla,S; Wurapa,K.E |
| EPI368644 | NP | United States | 2012-Jan-26 | A/Texas/23/2012 | Texas Department of State Health Services-Laboratory Services | Centers for Disease Control and Prevention | NA |
| EPI368645 | NS | United States | 2012-Jan-26 | A/Texas/23/2012 | Texas Department of State Health Services-Laboratory Services | Centers for Disease Control and Prevention | NA |
| EPI368646 | PA | United States | 2012-Jan-26 | A/Texas/23/2012 | Texas Department of State Health Services-Laboratory Services | Centers for Disease Control and Prevention | NA |
| EPI368647 | PB2 | United States | 2012-Jan-26 | A/Texas/23/2012 | Texas Department of State Health Services-Laboratory Services | Centers for Disease Control and Prevention | NA |
| EPI368648 | PB1 | United States | 2012-Jan-26 | A/Texas/23/2012 | Texas Department of State Health Services-Laboratory Services | Centers for Disease Control and Prevention | NA |
| EPI366330 | MP | United States | 2012-Jan-26 | A/Texas/23/2012 | Texas Department of State Health Services-Laboratory Services | Centers for Disease Control and Prevention | NA |
| EPI366331 | NA | United States | 2012-Jan-26 | A/Texas/23/2012 | Texas Department of State Health Services-Laboratory Services | Centers for Disease Control and Prevention | NA |
| EPI366332 | HA | United States | 2012-Jan-26 | A/Texas/23/2012 | Texas Department of State Health Services-Laboratory Services | Centers for Disease Control and Prevention | NA |
| EPI393764 | MP | United States | 2012-Jun-17 | A/Washington/24/2012 | Washington State Public Health Laboratory | Centers for Disease Control and Prevention | NA |
| EPI393765 | NA | United States | 2012-Jun-17 | A/Washington/24/2012 | Washington State Public Health Laboratory | Centers for Disease Control and Prevention | NA |
| EPI394884 | HA | United States | 2012-Jun-17 | A/Washington/24/2012 | Washington State Public Health Laboratory | Centers for Disease Control and Prevention | NA |
| EPI418238 | NP | United States | 2012-Jun-17 | A/Washington/24/2012 | Washington State Public Health Laboratory | Centers for Disease Control and Prevention | NA |
| EPI418239 | NS | United States | 2012-Jun-17 | A/Washington/24/2012 | Washington State Public Health Laboratory | Centers for Disease Control and Prevention | NA |
| EPI418240 | PA | United States | 2012-Jun-17 | A/Washington/24/2012 | Washington State Public Health Laboratory | Centers for Disease Control and Prevention | NA |
| EPI418241 | PB2 | United States | 2012-Jun-17 | A/Washington/24/2012 | Washington State Public Health Laboratory | Centers for Disease Control and Prevention | NA |
| EPI418242 | PB1 | United States | 2012-Jun-17 | A/Washington/24/2012 | Washington State Public Health Laboratory | Centers for Disease Control and Prevention | NA |
| EPI468805 | PB2 | India | 2012-Mar-07 | A/India/VD122268/2012 |  | Other Database Import | Potdar,V.A.; Dakhave,M.R.; Patil,K.N.; Kadam,A.A.; Mullick,J.; Chadha,M.S. |
| EPI468806 | PB1 | India | 2012-Mar-07 | A/India/VD122268/2012 |  | Other Database Import | Potdar,V.A.; Dakhave,M.R.; Patil,K.N.; Kadam,A.A.; Mullick,J.; Chadha,M.S. |
| EPI468808 | HA | India | 2012-Mar-07 | A/India/VD122268/2012 |  | Other Database Import | Potdar,V.A.; Dakhave,M.R.; Patil,K.N.; Kadam,A.A.; Mullick,J.; Chadha,M.S. |
| EPI468809 | NP | India | 2012-Mar-07 | A/India/VD122268/2012 |  | Other Database Import | Potdar,V.A.; Dakhave,M.R.; Patil,K.N.; Kadam,A.A.; Mullick,J.; Chadha,M.S. |
| EPI468810 | NA | India | 2012-Mar-07 | A/India/VD122268/2012 |  | Other Database Import | Potdar,V.A.; Dakhave,M.R.; Patil,K.N.; Kadam,A.A.; Mullick,J.; Chadha,M.S. |
| EPI468811 | MP | India | 2012-Mar-07 | A/India/VD122268/2012 |  | Other Database Import | Potdar,V.A.; Dakhave,M.R.; Patil,K.N.; Kadam,A.A.; Mullick,J.; Chadha,M.S. |
| EPI468812 | NS | India | 2012-Mar-07 | A/India/VD122268/2012 |  | Other Database Import | Potdar,V.A.; Dakhave,M.R.; Patil,K.N.; Kadam,A.A.; Mullick,J.; Chadha,M.S. |
| EPI468839 | PA | India | 2012-Mar-07 | A/India/VD122268/2012 |  | Other Database Import | Potdar,V.A.; Dakhave,M.R.; Patil,K.N.; Kadam,A.A.; Mullick,J.; Chadha,M.S. |
| EPI468894 | PB2 | India | 2012-Jan-19 | A/India/Nsk12388/2012 |  | Other Database Import | Potdar,V.A.; Dakhave,M.R.; Patil,K.N.; Kadam,A.A.; Mullick,J.; Chadha,M.S. |
| EPI468895 | PB1 | India | 2012-Jan-19 | A/India/Nsk12388/2012 |  | Other Database Import | Potdar,V.A.; Dakhave,M.R.; Patil,K.N.; Kadam,A.A.; Mullick,J.; Chadha,M.S. |
| EPI468897 | HA | India | 2012-Jan-19 | A/India/Nsk12388/2012 |  | Other Database Import | Potdar,V.A.; Dakhave,M.R.; Patil,K.N.; Kadam,A.A.; Mullick,J.; Chadha,M.S. |
| EPI468898 | NP | India | 2012-Jan-19 | A/India/Nsk12388/2012 |  | Other Database Import | Potdar,V.A.; Dakhave,M.R.; Patil,K.N.; Kadam,A.A.; Mullick,J.; Chadha,M.S. |
| EPI468899 | NA | India | 2012-Jan-19 | A/India/Nsk12388/2012 |  | Other Database Import | Potdar,V.A.; Dakhave,M.R.; Patil,K.N.; Kadam,A.A.; Mullick,J.; Chadha,M.S. |
| EPI468900 | MP | India | 2012-Jan-19 | A/India/Nsk12388/2012 |  | Other Database Import | Potdar,V.A.; Dakhave,M.R.; Patil,K.N.; Kadam,A.A.; Mullick,J.; Chadha,M.S. |
| EPI468901 | NS | India | 2012-Jan-19 | A/India/Nsk12388/2012 |  | Other Database Import | Potdar,V.A.; Dakhave,M.R.; Patil,K.N.; Kadam,A.A.; Mullick,J.; Chadha,M.S. |
| EPI468937 | PA | India | 2012-Jan-19 | A/India/Nsk12388/2012 |  | Other Database Import | Potdar,V.A.; Dakhave,M.R.; Patil,K.N.; Kadam,A.A.; Mullick,J.; Chadha,M.S. |
| EPI468948 | PA | India | 2012-Feb-24 | A/India/P121773/2012 |  | Other Database Import | Potdar,V.A.; Dakhave,M.R.; Patil,K.N.; Kadam,A.A.; Mullick,J.; Chadha,M.S. |
| EPI468918 | PB2 | India | 2012-Feb-24 | A/India/P121773/2012 |  | Other Database Import | Potdar,V.A.; Dakhave,M.R.; Patil,K.N.; Kadam,A.A.; Mullick,J.; Chadha,M.S. |
| EPI468919 | PB1 | India | 2012-Feb-24 | A/India/P121773/2012 |  | Other Database Import | Potdar,V.A.; Dakhave,M.R.; Patil,K.N.; Kadam,A.A.; Mullick,J.; Chadha,M.S. |
| EPI468921 | HA | India | 2012-Feb-24 | A/India/P121773/2012 |  | Other Database Import | Potdar,V.A.; Dakhave,M.R.; Patil,K.N.; Kadam,A.A.; Mullick,J.; Chadha,M.S. |
| EPI468922 | NP | India | 2012-Feb-24 | A/India/P121773/2012 |  | Other Database Import | Potdar,V.A.; Dakhave,M.R.; Patil,K.N.; Kadam,A.A.; Mullick,J.; Chadha,M.S. |
| EPI468923 | NA | India | 2012-Feb-24 | A/India/P121773/2012 |  | Other Database Import | Potdar,V.A.; Dakhave,M.R.; Patil,K.N.; Kadam,A.A.; Mullick,J.; Chadha,M.S. |
| EPI468924 | MP | India | 2012-Feb-24 | A/India/P121773/2012 |  | Other Database Import | Potdar,V.A.; Dakhave,M.R.; Patil,K.N.; Kadam,A.A.; Mullick,J.; Chadha,M.S. |
| EPI468925 | NS | India | 2012-Feb-24 | A/India/P121773/2012 |  | Other Database Import | Potdar,V.A.; Dakhave,M.R.; Patil,K.N.; Kadam,A.A.; Mullick,J.; Chadha,M.S. |
| EPI536168 | PA | France | 2012-Mar-21 | A/Paris/1119/2012 |  | Institut Pasteur | Barbezange, Cyril |
| EPI536172 | MP | France | 2012-Mar-21 | A/Paris/1119/2012 |  | Institut Pasteur | Barbezange, Cyril |
| EPI536170 | NP | France | 2012-Mar-21 | A/Paris/1119/2012 |  | Institut Pasteur | Barbezange, Cyril |
| EPI536171 | NA | France | 2012-Mar-21 | A/Paris/1119/2012 |  | Institut Pasteur | Barbezange, Cyril |
| EPI536166 | PB2 | France | 2012-Mar-21 | A/Paris/1119/2012 |  | Institut Pasteur | Barbezange, Cyril |
| EPI536167 | PB1 | France | 2012-Mar-21 | A/Paris/1119/2012 |  | Institut Pasteur | Barbezange, Cyril |
| EPI536169 | HA | France | 2012-Mar-21 | A/Paris/1119/2012 |  | Institut Pasteur | Barbezange, Cyril |
| EPI536173 | NS | France | 2012-Mar-21 | A/Paris/1119/2012 |  | Institut Pasteur | Barbezange, Cyril |
| EPI543228 | MP | Spain | 2011-Feb-08 | A/Madrid/RR7495/2011 | Instituto de Salud Carlos III | Instituto de Salud Carlos III | Pozo,F; Ledesma,J; Calderon,A.; Gonzalez -Esguevillas,M.; Molinero,M.; Casas,I. |
| EPI543231 | PA | Spain | 2011-Feb-08 | A/Madrid/RR7495/2011 | Instituto de Salud Carlos III | Instituto de Salud Carlos III | Pozo,F; Ledesma,J; Calderon,A.; Gonzalez -Esguevillas,M.; Molinero,M.; Casas,I. |
| EPI543232 | PB1 | Spain | 2011-Feb-08 | A/Madrid/RR7495/2011 | Instituto de Salud Carlos III | Instituto de Salud Carlos III | Pozo,F; Ledesma,J; Calderon,A.; Gonzalez -Esguevillas,M.; Molinero,M.; Casas,I. |
| EPI543233 | PB2 | Spain | 2011-Feb-08 | A/Madrid/RR7495/2011 | Instituto de Salud Carlos III | Instituto de Salud Carlos III | Pozo,F; Ledesma,J; Calderon,A.; Gonzalez -Esguevillas,M.; Molinero,M.; Casas,I. |
| EPI543158 | NA | Spain | 2011-Feb-08 | A/Madrid/RR7495/2011 | Instituto de Salud Carlos III | Instituto de Salud Carlos III | Pozo,F; Ledesma,J; Calderon,A.; Gonzalez -Esguevillas,M.; Molinero,M.; Casas,I. |
| EPI543079 | HA | Spain | 2011-Feb-08 | A/Madrid/RR7495/2011 | Instituto de Salud Carlos III | Instituto de Salud Carlos III | Pozo,F; Ledesma,J; Calderon,A.; Gonzalez -Esguevillas,M.; Molinero,M.; Casas,I. |
| EPI543229 | NP | Spain | 2011-Feb-08 | A/Madrid/RR7495/2011 | Instituto de Salud Carlos III | Instituto de Salud Carlos III | Pozo,F; Ledesma,J; Calderon,A.; Gonzalez -Esguevillas,M.; Molinero,M.; Casas,I. |
| EPI543230 | NS | Spain | 2011-Feb-08 | A/Madrid/RR7495/2011 | Instituto de Salud Carlos III | Instituto de Salud Carlos III | Pozo,F; Ledesma,J; Calderon,A.; Gonzalez -Esguevillas,M.; Molinero,M.; Casas,I. |
| EPI324788 | PA | Russian Federation | 2011-Jan-22 | A/Moscow/IIV-83/2011 |  | Other Database Import | Poglazov,A.B.; Shlyapnikova,O.V.; Prilipov,A.G. |
| EPI324789 | PB2 | Russian Federation | 2011-Jan-22 | A/Moscow/IIV-83/2011 |  | Other Database Import | Poglazov,A.B.; Shlyapnikova,O.V.; Prilipov,A.G. |
| EPI324790 | PB1 | Russian Federation | 2011-Jan-22 | A/Moscow/IIV-83/2011 |  | Other Database Import | Poglazov,A.B.; Shlyapnikova,O.V.; Prilipov,A.G. |
| EPI324786 | NA | Russian Federation | 2011-Jan-22 | A/Moscow/IIV-83/2011 |  | Other Database Import | Poglazov,A.B.; Shlyapnikova,O.V.; Prilipov,A.G. |
| EPI324787 | NP | Russian Federation | 2011-Jan-22 | A/Moscow/IIV-83/2011 |  | Other Database Import | Poglazov,A.B.; Shlyapnikova,O.V.; Prilipov,A.G. |
| EPI324791 | MP | Russian Federation | 2011-Jan-22 | A/Moscow/IIV-83/2011 |  | Other Database Import | Poglazov,A.B.; Shlyapnikova,O.V.; Prilipov,A.G. |
| EPI324792 | NS | Russian Federation | 2011-Jan-22 | A/Moscow/IIV-83/2011 |  | Other Database Import | Poglazov,A.B.; Shlyapnikova,O.V.; Prilipov,A.G. |
| EPI324793 | HA | Russian Federation | 2011-Jan-22 | A/Moscow/IIV-83/2011 |  | Other Database Import | Poglazov,A.B.; Shlyapnikova,O.V.; Prilipov,A.G. |
| EPI356423 | NP | Germany | 2011-Jan-01 | A/Germany-MV/R26/2011 |  | Friedrich-Loeffler-Institut | Goller, Katja; Starick, Elke |
| EPI356424 | NS | Germany | 2011-Jan-01 | A/Germany-MV/R26/2011 |  | Friedrich-Loeffler-Institut | Goller, Katja; Starick, Elke |
| EPI356425 | MP | Germany | 2011-Jan-01 | A/Germany-MV/R26/2011 |  | Friedrich-Loeffler-Institut | Goller, Katja; Starick, Elke |
| EPI356426 | PA | Germany | 2011-Jan-01 | A/Germany-MV/R26/2011 |  | Friedrich-Loeffler-Institut | Goller, Katja; Starick, Elke |
| EPI356427 | PB2 | Germany | 2011-Jan-01 | A/Germany-MV/R26/2011 |  | Friedrich-Loeffler-Institut | Goller, Katja; Starick, Elke |
| EPI356428 | PB1 | Germany | 2011-Jan-01 | A/Germany-MV/R26/2011 |  | Friedrich-Loeffler-Institut | Goller, Katja; Starick, Elke |
| EPI356429 | NA | Germany | 2011-Jan-01 | A/Germany-MV/R26/2011 |  | Friedrich-Loeffler-Institut | Goller, Katja; Starick, Elke |
| EPI356430 | HA | Germany | 2011-Jan-01 | A/Germany-MV/R26/2011 |  | Friedrich-Loeffler-Institut | Goller, Katja; Starick, Elke |
| EPI370303 | PA | Australia | 2011-Aug-23 | A/PERTH/299/2011 | Pathwest QE II Medical Centre | WHO Collaborating Centre for Reference and Research on Influenza | Deng,Y-M; Iannello,P; Caldwell,N; Komadina,N. |
| EPI370304 | PB2 | Australia | 2011-Aug-23 | A/PERTH/299/2011 | Pathwest QE II Medical Centre | WHO Collaborating Centre for Reference and Research on Influenza | Deng,Y-M; Iannello,P; Caldwell,N; Komadina,N. |
| EPI370305 | PB1 | Australia | 2011-Aug-23 | A/PERTH/299/2011 | Pathwest QE II Medical Centre | WHO Collaborating Centre for Reference and Research on Influenza | Deng,Y-M; Iannello,P; Caldwell,N; Komadina,N. |
| EPI370300 | NP | Australia | 2011-Aug-23 | A/PERTH/299/2011 | Pathwest QE II Medical Centre | WHO Collaborating Centre for Reference and Research on Influenza | Deng,Y-M; Iannello,P; Caldwell,N; Komadina,N. |
| EPI370301 | NS | Australia | 2011-Aug-23 | A/PERTH/299/2011 | Pathwest QE II Medical Centre | WHO Collaborating Centre for Reference and Research on Influenza | Deng,Y-M; Iannello,P; Caldwell,N; Komadina,N. |
| EPI370302 | MP | Australia | 2011-Aug-23 | A/PERTH/299/2011 | Pathwest QE II Medical Centre | WHO Collaborating Centre for Reference and Research on Influenza | Deng,Y-M; Iannello,P; Caldwell,N; Komadina,N. |
| EPI370306 | NA | Australia | 2011-Aug-23 | A/PERTH/299/2011 | Pathwest QE II Medical Centre | WHO Collaborating Centre for Reference and Research on Influenza | Deng,Y-M; Iannello,P; Caldwell,N; Komadina,N. |
| EPI370307 | HA | Australia | 2011-Aug-23 | A/PERTH/299/2011 | Pathwest QE II Medical Centre | WHO Collaborating Centre for Reference and Research on Influenza | Deng,Y-M; Iannello,P; Caldwell,N; Komadina,N. |
| EPI394281 | MP | Fiji | 2012-Jul-16 | A/FIJI/4/2012 | National Centre for Scientific Services for Virology and Vector Borne Diseases | WHO Collaborating Centre for Reference and Research on Influenza | Deng,Y-M; Iannello,P; Caldwell,N; Jelley,L; Komadina,N |
| EPI394282 | NA | Fiji | 2012-Jul-16 | A/FIJI/4/2012 | National Centre for Scientific Services for Virology and Vector Borne Diseases | WHO Collaborating Centre for Reference and Research on Influenza | Deng,Y-M; Iannello,P; Caldwell,N; Jelley,L; Komadina,N |
| EPI394283 | HA | Fiji | 2012-Jul-16 | A/FIJI/4/2012 | National Centre for Scientific Services for Virology and Vector Borne Diseases | WHO Collaborating Centre for Reference and Research on Influenza | Deng,Y-M; Iannello,P; Caldwell,N; Jelley,L; Komadina,N |
| EPI450413 | NP | Fiji | 2012-Jul-16 | A/FIJI/4/2012 | National Centre for Scientific Services for Virology and Vector Borne Diseases | WHO Collaborating Centre for Reference and Research on Influenza | Deng,Y-M; Iannello,P; Caldwell,N; Jelley,L; Komadina,N |
| EPI450414 | NS | Fiji | 2012-Jul-16 | A/FIJI/4/2012 | National Centre for Scientific Services for Virology and Vector Borne Diseases | WHO Collaborating Centre for Reference and Research on Influenza | Deng,Y-M; Iannello,P; Caldwell,N; Jelley,L; Komadina,N |
| EPI450415 | PA | Fiji | 2012-Jul-16 | A/FIJI/4/2012 | National Centre for Scientific Services for Virology and Vector Borne Diseases | WHO Collaborating Centre for Reference and Research on Influenza | Deng,Y-M; Iannello,P; Caldwell,N; Jelley,L; Komadina,N |
| EPI450416 | PB2 | Fiji | 2012-Jul-16 | A/FIJI/4/2012 | National Centre for Scientific Services for Virology and Vector Borne Diseases | WHO Collaborating Centre for Reference and Research on Influenza | Deng,Y-M; Iannello,P; Caldwell,N; Jelley,L; Komadina,N |
| EPI450417 | PB1 | Fiji | 2012-Jul-16 | A/FIJI/4/2012 | National Centre for Scientific Services for Virology and Vector Borne Diseases | WHO Collaborating Centre for Reference and Research on Influenza | Deng,Y-M; Iannello,P; Caldwell,N; Jelley,L; Komadina,N |
| EPI507595 | NP | Portugal | 2013-Mar-01 | A/Lisbon/137/2013 | National Institute for Medical Research | Centers for Disease Control and Prevention | NA |
| EPI507596 | NS | Portugal | 2013-Mar-01 | A/Lisbon/137/2013 | National Institute for Medical Research | Centers for Disease Control and Prevention | NA |
| EPI507597 | MP | Portugal | 2013-Mar-01 | A/Lisbon/137/2013 | National Institute for Medical Research | Centers for Disease Control and Prevention | NA |
| EPI507598 | PA | Portugal | 2013-Mar-01 | A/Lisbon/137/2013 | National Institute for Medical Research | Centers for Disease Control and Prevention | NA |
| EPI507599 | PB2 | Portugal | 2013-Mar-01 | A/Lisbon/137/2013 | National Institute for Medical Research | Centers for Disease Control and Prevention | NA |
| EPI507600 | PB1 | Portugal | 2013-Mar-01 | A/Lisbon/137/2013 | National Institute for Medical Research | Centers for Disease Control and Prevention | NA |
| EPI507601 | NA | Portugal | 2013-Mar-01 | A/Lisbon/137/2013 | National Institute for Medical Research | Centers for Disease Control and Prevention | NA |
| EPI507602 | HA | Portugal | 2013-Mar-01 | A/Lisbon/137/2013 | National Institute for Medical Research | Centers for Disease Control and Prevention | NA |
| EPI614470 | PB1 | Vietnam | 2013-Feb-28 | A/Vietnam/3050/2013 | National Institute of Hygiene and Epidemiology | Centers for Disease Control and Prevention | NA |
| EPI614463 | NP | Vietnam | 2013-Feb-28 | A/Vietnam/3050/2013 | National Institute of Hygiene and Epidemiology | Centers for Disease Control and Prevention | NA |
| EPI614464 | NS | Vietnam | 2013-Feb-28 | A/Vietnam/3050/2013 | National Institute of Hygiene and Epidemiology | Centers for Disease Control and Prevention | NA |
| EPI614465 | MP | Vietnam | 2013-Feb-28 | A/Vietnam/3050/2013 | National Institute of Hygiene and Epidemiology | Centers for Disease Control and Prevention | NA |
| EPI614467 | PA | Vietnam | 2013-Feb-28 | A/Vietnam/3050/2013 | National Institute of Hygiene and Epidemiology | Centers for Disease Control and Prevention | NA |
| EPI614468 | PB2 | Vietnam | 2013-Feb-28 | A/Vietnam/3050/2013 | National Institute of Hygiene and Epidemiology | Centers for Disease Control and Prevention | NA |
| EPI614471 | NA | Vietnam | 2013-Feb-28 | A/Vietnam/3050/2013 | National Institute of Hygiene and Epidemiology | Centers for Disease Control and Prevention | NA |
| EPI614472 | HA | Vietnam | 2013-Feb-28 | A/Vietnam/3050/2013 | National Institute of Hygiene and Epidemiology | Centers for Disease Control and Prevention | NA |
| EPI177294 | HA | United States | 2009-Apr-09 | A/California/07/2009 | Naval Health Research Center | Centers for Disease Control and Prevention | NA |
| EPI176615 | PB1 | United States | 2009-Apr-09 | A/California/07/2009 | Naval Health Research Center | Centers for Disease Control and Prevention | NA |
| EPI176616 | PB2 | United States | 2009-Apr-09 | A/California/07/2009 | Naval Health Research Center | Centers for Disease Control and Prevention | NA |
| EPI176617 | PA | United States | 2009-Apr-09 | A/California/07/2009 | Naval Health Research Center | Centers for Disease Control and Prevention | NA |
| EPI176618 | NS | United States | 2009-Apr-09 | A/California/07/2009 | Naval Health Research Center | Centers for Disease Control and Prevention | NA |
| EPI176619 | MP | United States | 2009-Apr-09 | A/California/07/2009 | Naval Health Research Center | Centers for Disease Control and Prevention | NA |
| EPI185379 | NA | United States | 2009-Apr-09 | A/California/07/2009 | Naval Health Research Center | Centers for Disease Control and Prevention | NA |
| EPI184298 | NP | United States | 2009-Apr-09 | A/California/07/2009 | Naval Health Research Center | Centers for Disease Control and Prevention | NA |
| EPI263455 | PB2 | Singapore | 2010-Jan-27 | A/Singapore/KK066/2010 |  | Other Database Import | Phuah,S.P.; Meng,S.; Teo,M.L.; Yong,T.Y.; Loh,P.L.; Quek,D.; Ho,Y.L.; Tan,Y.L.; Cui,L. |
| EPI263456 | PB1 | Singapore | 2010-Jan-27 | A/Singapore/KK066/2010 |  | Other Database Import | Phuah,S.P.; Meng,S.; Teo,M.L.; Yong,T.Y.; Loh,P.L.; Quek,D.; Ho,Y.L.; Tan,Y.L.; Cui,L. |
| EPI263457 | PA | Singapore | 2010-Jan-27 | A/Singapore/KK066/2010 |  | Other Database Import | Phuah,S.P.; Meng,S.; Teo,M.L.; Yong,T.Y.; Loh,P.L.; Quek,D.; Ho,Y.L.; Tan,Y.L.; Cui,L. |
| EPI263458 | HA | Singapore | 2010-Jan-27 | A/Singapore/KK066/2010 |  | Other Database Import | Phuah,S.P.; Meng,S.; Teo,M.L.; Yong,T.Y.; Loh,P.L.; Quek,D.; Ho,Y.L.; Tan,Y.L.; Cui,L. |
| EPI263459 | NP | Singapore | 2010-Jan-27 | A/Singapore/KK066/2010 |  | Other Database Import | Phuah,S.P.; Meng,S.; Teo,M.L.; Yong,T.Y.; Loh,P.L.; Quek,D.; Ho,Y.L.; Tan,Y.L.; Cui,L. |
| EPI263460 | NA | Singapore | 2010-Jan-27 | A/Singapore/KK066/2010 |  | Other Database Import | Phuah,S.P.; Meng,S.; Teo,M.L.; Yong,T.Y.; Loh,P.L.; Quek,D.; Ho,Y.L.; Tan,Y.L.; Cui,L. |
| EPI263461 | MP | Singapore | 2010-Jan-27 | A/Singapore/KK066/2010 |  | Other Database Import | Phuah,S.P.; Meng,S.; Teo,M.L.; Yong,T.Y.; Loh,P.L.; Quek,D.; Ho,Y.L.; Tan,Y.L.; Cui,L. |
| EPI263462 | NS | Singapore | 2010-Jan-27 | A/Singapore/KK066/2010 |  | Other Database Import | Phuah,S.P.; Meng,S.; Teo,M.L.; Yong,T.Y.; Loh,P.L.; Quek,D.; Ho,Y.L.; Tan,Y.L.; Cui,L. |
| EPI309986 | PA | United States | 2011-Feb-25 | A/Maryland/07/2011 | Maryland Department of Health and Mental Hygiene | Centers for Disease Control and Prevention | NA |
| EPI309987 | PB2 | United States | 2011-Feb-25 | A/Maryland/07/2011 | Maryland Department of Health and Mental Hygiene | Centers for Disease Control and Prevention | NA |
| EPI309988 | PB1 | United States | 2011-Feb-25 | A/Maryland/07/2011 | Maryland Department of Health and Mental Hygiene | Centers for Disease Control and Prevention | NA |
| EPI309985 | MP | United States | 2011-Feb-25 | A/Maryland/07/2011 | Maryland Department of Health and Mental Hygiene | Centers for Disease Control and Prevention | NA |
| EPI309989 | NA | United States | 2011-Feb-25 | A/Maryland/07/2011 | Maryland Department of Health and Mental Hygiene | Centers for Disease Control and Prevention | NA |
| EPI309990 | HA | United States | 2011-Feb-25 | A/Maryland/07/2011 | Maryland Department of Health and Mental Hygiene | Centers for Disease Control and Prevention | NA |
| EPI316372 | NP | United States | 2011-Feb-25 | A/Maryland/07/2011 | Maryland Department of Health and Mental Hygiene | Centers for Disease Control and Prevention | NA |
| EPI316373 | NS | United States | 2011-Feb-25 | A/Maryland/07/2011 | Maryland Department of Health and Mental Hygiene | Centers for Disease Control and Prevention | NA |
| EPI330325 | NP | China | 2011-Mar-07 | A/Guangdong-Xinxing/SWL198/2011 | WHO Chinese National Influenza Center | WHO Chinese National Influenza Center | Lan,Yu;Wang,Dayan;Li,Xiyan;Huang,Weijuan;Zhao,Xiang;Chen,Yaoyao;Yang,Lei;Shu,Yuelong |
| EPI330326 | NS | China | 2011-Mar-07 | A/Guangdong-Xinxing/SWL198/2011 | WHO Chinese National Influenza Center | WHO Chinese National Influenza Center | Lan,Yu;Wang,Dayan;Li,Xiyan;Huang,Weijuan;Zhao,Xiang;Chen,Yaoyao;Yang,Lei;Shu,Yuelong |
| EPI330327 | MP | China | 2011-Mar-07 | A/Guangdong-Xinxing/SWL198/2011 | WHO Chinese National Influenza Center | WHO Chinese National Influenza Center | Lan,Yu;Wang,Dayan;Li,Xiyan;Huang,Weijuan;Zhao,Xiang;Chen,Yaoyao;Yang,Lei;Shu,Yuelong |
| EPI330330 | PB1 | China | 2011-Mar-07 | A/Guangdong-Xinxing/SWL198/2011 | WHO Chinese National Influenza Center | WHO Chinese National Influenza Center | Lan,Yu;Wang,Dayan;Li,Xiyan;Huang,Weijuan;Zhao,Xiang;Chen,Yaoyao;Yang,Lei;Shu,Yuelong |
| EPI330331 | NA | China | 2011-Mar-07 | A/Guangdong-Xinxing/SWL198/2011 | WHO Chinese National Influenza Center | WHO Chinese National Influenza Center | Lan,Yu;Wang,Dayan;Li,Xiyan;Huang,Weijuan;Zhao,Xiang;Chen,Yaoyao;Yang,Lei;Shu,Yuelong |
| EPI330332 | HA | China | 2011-Mar-07 | A/Guangdong-Xinxing/SWL198/2011 | WHO Chinese National Influenza Center | WHO Chinese National Influenza Center | Lan,Yu;Wang,Dayan;Li,Xiyan;Huang,Weijuan;Zhao,Xiang;Chen,Yaoyao;Yang,Lei;Shu,Yuelong |
| EPI330328 | PA | China | 2011-Mar-07 | A/Guangdong-Xinxing/SWL198/2011 | WHO Chinese National Influenza Center | WHO Chinese National Influenza Center | Lan,Yu;Wang,Dayan;Li,Xiyan;Huang,Weijuan;Zhao,Xiang;Chen,Yaoyao;Yang,Lei;Shu,Yuelong |
| EPI330329 | PB2 | China | 2011-Mar-07 | A/Guangdong-Xinxing/SWL198/2011 | WHO Chinese National Influenza Center | WHO Chinese National Influenza Center | Lan,Yu;Wang,Dayan;Li,Xiyan;Huang,Weijuan;Zhao,Xiang;Chen,Yaoyao;Yang,Lei;Shu,Yuelong |
| EPI349572 | PB2 | Taiwan | 2011-Feb-08 | A/Taiwan/4611/2011 |  | Other Database Import | Yang,J.R.; Huang,Y.P.; Chang,F.Y.; Hsu,L.C.; Lin,Y.C.; Su,C.H.; Chen,P.J.; Wu,H.S.; Liu,M.T.; Yang,J.-R.; Wu,H.-S.; Liu,M.-T. |
| EPI349659 | PA | Taiwan | 2011-Feb-08 | A/Taiwan/4611/2011 |  | Other Database Import | Yang,J.R.; Huang,Y.P.; Chang,F.Y.; Hsu,L.C.; Lin,Y.C.; Su,C.H.; Chen,P.J.; Wu,H.S.; Liu,M.T.; Yang,J.-R.; Wu,H.-S.; Liu,M.-T. |
| EPI349717 | MP | Taiwan | 2011-Feb-08 | A/Taiwan/4611/2011 |  | Other Database Import | Yang,J.R.; Huang,Y.P.; Chang,F.Y.; Hsu,L.C.; Lin,Y.C.; Su,C.H.; Chen,P.J.; Wu,H.S.; Liu,M.T.; Yang,J.-R.; Wu,H.-S.; Liu,M.-T. |
| EPI349543 | HA | Taiwan | 2011-Feb-08 | A/Taiwan/4611/2011 |  | Other Database Import | Yang,J.R.; Huang,Y.P.; Chang,F.Y.; Hsu,L.C.; Lin,Y.C.; Su,C.H.; Chen,P.J.; Wu,H.S.; Liu,M.T.; Yang,J.-R.; Wu,H.-S.; Liu,M.-T. |
| EPI349688 | NP | Taiwan | 2011-Feb-08 | A/Taiwan/4611/2011 |  | Other Database Import | Yang,J.R.; Huang,Y.P.; Chang,F.Y.; Hsu,L.C.; Lin,Y.C.; Su,C.H.; Chen,P.J.; Wu,H.S.; Liu,M.T.; Yang,J.-R.; Wu,H.-S.; Liu,M.-T. |
| EPI349746 | NS | Taiwan | 2011-Feb-08 | A/Taiwan/4611/2011 |  | Other Database Import | Yang,J.R.; Huang,Y.P.; Chang,F.Y.; Hsu,L.C.; Lin,Y.C.; Su,C.H.; Chen,P.J.; Wu,H.S.; Liu,M.T.; Yang,J.-R.; Wu,H.-S.; Liu,M.-T. |
| EPI349601 | NA | Taiwan | 2011-Feb-08 | A/Taiwan/4611/2011 |  | Other Database Import | Yang,J.R.; Huang,Y.P.; Chang,F.Y.; Hsu,L.C.; Lin,Y.C.; Su,C.H.; Chen,P.J.; Wu,H.S.; Liu,M.T.; Yang,J.-R.; Wu,H.-S.; Liu,M.-T. |
| EPI349630 | PB1 | Taiwan | 2011-Feb-08 | A/Taiwan/4611/2011 |  | Other Database Import | Yang,J.R.; Huang,Y.P.; Chang,F.Y.; Hsu,L.C.; Lin,Y.C.; Su,C.H.; Chen,P.J.; Wu,H.S.; Liu,M.T.; Yang,J.-R.; Wu,H.-S.; Liu,M.-T. |
| EPI354377 | PA | India | 2010-Sep-09 | A/India/GWL_DSC/2010 |  | Other Database Import | Sharma,S.; Parida,M.M.; Shukla,J.; Joshi,G.; Dash,P.; Rao,P.V.L. |
| EPI354373 | NA | India | 2010-Sep-09 | A/India/GWL_DSC/2010 |  | Other Database Import | Sharma,S.; Parida,M.M.; Shukla,J.; Joshi,G.; Dash,P.; Rao,P.V.L. |
| EPI354375 | NP | India | 2010-Sep-09 | A/India/GWL_DSC/2010 |  | Other Database Import | Sharma,S.; Parida,M.M.; Shukla,J.; Joshi,G.; Dash,P.; Rao,P.V.L. |
| EPI354379 | HA | India | 2010-Sep-09 | A/India/GWL_DSC/2010 |  | Other Database Import | Sharma,S.; Parida,M.M.; Shukla,J.; Joshi,G.; Dash,P.; Rao,P.V.L. |
| EPI354394 | NS | India | 2010-Sep-09 | A/India/GWL_DSC/2010 |  | Other Database Import | Sharma,S.; Parida,M.M.; Shukla,J.; Joshi,G.; Dash,P.; Rao,P.V.L. |
| EPI354397 | PB1 | India | 2010-Sep-09 | A/India/GWL_DSC/2010 |  | Other Database Import | Sharma,S.; Parida,M.M.; Shukla,J.; Joshi,G.; Dash,P.; Rao,P.V.L. |
| EPI354378 | PB2 | India | 2010-Sep-09 | A/India/GWL_DSC/2010 |  | Other Database Import | Sharma,S.; Parida,M.M.; Shukla,J.; Joshi,G.; Dash,P.; Rao,P.V.L. |
| EPI354392 | MP | India | 2010-Sep-09 | A/India/GWL_DSC/2010 |  | Other Database Import | Sharma,S.; Parida,M.M.; Shukla,J.; Joshi,G.; Dash,P.; Rao,P.V.L. |
| EPI354380 | PA | India | 2010-Mar-22 | A/India/Blore/2010 |  | Other Database Import | Sharma,S.; Parida,M.M.; Shukla,J.; Joshi,G.; Dash,P.; Rao,P.V.L. |
| EPI354381 | PB2 | India | 2010-Mar-22 | A/India/Blore/2010 |  | Other Database Import | Sharma,S.; Parida,M.M.; Shukla,J.; Joshi,G.; Dash,P.; Rao,P.V.L. |
| EPI354374 | NA | India | 2010-Mar-22 | A/India/Blore/2010 |  | Other Database Import | Sharma,S.; Parida,M.M.; Shukla,J.; Joshi,G.; Dash,P.; Rao,P.V.L. |
| EPI354382 | HA | India | 2010-Mar-22 | A/India/Blore/2010 |  | Other Database Import | Sharma,S.; Parida,M.M.; Shukla,J.; Joshi,G.; Dash,P.; Rao,P.V.L. |
| EPI354393 | MP | India | 2010-Mar-22 | A/India/Blore/2010 |  | Other Database Import | Sharma,S.; Parida,M.M.; Shukla,J.; Joshi,G.; Dash,P.; Rao,P.V.L. |
| EPI354396 | NS | India | 2010-Mar-22 | A/India/Blore/2010 |  | Other Database Import | Sharma,S.; Parida,M.M.; Shukla,J.; Joshi,G.; Dash,P.; Rao,P.V.L. |
| EPI354398 | PB1 | India | 2010-Mar-22 | A/India/Blore/2010 |  | Other Database Import | Sharma,S.; Parida,M.M.; Shukla,J.; Joshi,G.; Dash,P.; Rao,P.V.L. |
| EPI354376 | NP | India | 2010-Mar-22 | A/India/Blore/2010 |  | Other Database Import | Sharma,S.; Parida,M.M.; Shukla,J.; Joshi,G.; Dash,P.; Rao,P.V.L. |
| EPI536159 | PB1 | France | 2011-Jan-28 | A/Paris/1230/2011 |  | Institut Pasteur | Barbezange, Cyril |
| EPI536163 | NA | France | 2011-Jan-28 | A/Paris/1230/2011 |  | Institut Pasteur | Barbezange, Cyril |
| EPI536162 | NP | France | 2011-Jan-28 | A/Paris/1230/2011 |  | Institut Pasteur | Barbezange, Cyril |
| EPI536161 | HA | France | 2011-Jan-28 | A/Paris/1230/2011 |  | Institut Pasteur | Barbezange, Cyril |
| EPI536164 | MP | France | 2011-Jan-28 | A/Paris/1230/2011 |  | Institut Pasteur | Barbezange, Cyril |
| EPI536158 | PB2 | France | 2011-Jan-28 | A/Paris/1230/2011 |  | Institut Pasteur | Barbezange, Cyril |
| EPI536160 | PA | France | 2011-Jan-28 | A/Paris/1230/2011 |  | Institut Pasteur | Barbezange, Cyril |
| EPI536165 | NS | France | 2011-Jan-28 | A/Paris/1230/2011 |  | Institut Pasteur | Barbezange, Cyril |
| EPI650018 | NP | French Guiana | 2015-Jun-15 | A/French Guiana/6135/2015 | National Influenza Center French Guiana and French Indies | Centers for Disease Control and Prevention | NA |
| EPI650019 | NS | French Guiana | 2015-Jun-15 | A/French Guiana/6135/2015 | National Influenza Center French Guiana and French Indies | Centers for Disease Control and Prevention | NA |
| EPI650020 | MP | French Guiana | 2015-Jun-15 | A/French Guiana/6135/2015 | National Influenza Center French Guiana and French Indies | Centers for Disease Control and Prevention | NA |
| EPI650021 | PA | French Guiana | 2015-Jun-15 | A/French Guiana/6135/2015 | National Influenza Center French Guiana and French Indies | Centers for Disease Control and Prevention | NA |
| EPI650024 | NA | French Guiana | 2015-Jun-15 | A/French Guiana/6135/2015 | National Influenza Center French Guiana and French Indies | Centers for Disease Control and Prevention | NA |
| EPI650025 | HA | French Guiana | 2015-Jun-15 | A/French Guiana/6135/2015 | National Influenza Center French Guiana and French Indies | Centers for Disease Control and Prevention | NA |
| EPI650022 | PB2 | French Guiana | 2015-Jun-15 | A/French Guiana/6135/2015 | National Influenza Center French Guiana and French Indies | Centers for Disease Control and Prevention | NA |
| EPI650023 | PB1 | French Guiana | 2015-Jun-15 | A/French Guiana/6135/2015 | National Influenza Center French Guiana and French Indies | Centers for Disease Control and Prevention | NA |
| EPI675799 | NP | Hong Kong (SAR) | 2015-Oct-08 | A/Hong Kong/15587/2015 | Government Virus Unit | Centers for Disease Control and Prevention | NA |
| EPI675800 | NS | Hong Kong (SAR) | 2015-Oct-08 | A/Hong Kong/15587/2015 | Government Virus Unit | Centers for Disease Control and Prevention | NA |
| EPI675801 | MP | Hong Kong (SAR) | 2015-Oct-08 | A/Hong Kong/15587/2015 | Government Virus Unit | Centers for Disease Control and Prevention | NA |
| EPI675802 | PA | Hong Kong (SAR) | 2015-Oct-08 | A/Hong Kong/15587/2015 | Government Virus Unit | Centers for Disease Control and Prevention | NA |
| EPI675803 | PB2 | Hong Kong (SAR) | 2015-Oct-08 | A/Hong Kong/15587/2015 | Government Virus Unit | Centers for Disease Control and Prevention | NA |
| EPI675804 | PB1 | Hong Kong (SAR) | 2015-Oct-08 | A/Hong Kong/15587/2015 | Government Virus Unit | Centers for Disease Control and Prevention | NA |
| EPI675805 | NA | Hong Kong (SAR) | 2015-Oct-08 | A/Hong Kong/15587/2015 | Government Virus Unit | Centers for Disease Control and Prevention | NA |
| EPI675806 | HA | Hong Kong (SAR) | 2015-Oct-08 | A/Hong Kong/15587/2015 | Government Virus Unit | Centers for Disease Control and Prevention | NA |
| EPI690865 | PA | Costa Rica | 2015-Oct-26 | A/Costa Rica/0373/2015 | Laboratorio Nacional de Influenza | Centers for Disease Control and Prevention | NA |
| EPI690866 | PB2 | Costa Rica | 2015-Oct-26 | A/Costa Rica/0373/2015 | Laboratorio Nacional de Influenza | Centers for Disease Control and Prevention | NA |
| EPI690867 | PB1 | Costa Rica | 2015-Oct-26 | A/Costa Rica/0373/2015 | Laboratorio Nacional de Influenza | Centers for Disease Control and Prevention | NA |
| EPI690862 | NP | Costa Rica | 2015-Oct-26 | A/Costa Rica/0373/2015 | Laboratorio Nacional de Influenza | Centers for Disease Control and Prevention | NA |
| EPI690863 | NS | Costa Rica | 2015-Oct-26 | A/Costa Rica/0373/2015 | Laboratorio Nacional de Influenza | Centers for Disease Control and Prevention | NA |
| EPI690864 | MP | Costa Rica | 2015-Oct-26 | A/Costa Rica/0373/2015 | Laboratorio Nacional de Influenza | Centers for Disease Control and Prevention | NA |
| EPI690868 | NA | Costa Rica | 2015-Oct-26 | A/Costa Rica/0373/2015 | Laboratorio Nacional de Influenza | Centers for Disease Control and Prevention | NA |
| EPI690869 | HA | Costa Rica | 2015-Oct-26 | A/Costa Rica/0373/2015 | Laboratorio Nacional de Influenza | Centers for Disease Control and Prevention | NA |
| EPI650072 | HA | Australia | 2015-Mar-12 | A/South Australia/22/2015 | WHO Collaborating Centre for Reference and Research on Influenza | Centers for Disease Control and Prevention | NA |
| EPI650065 | NP | Australia | 2015-Mar-12 | A/South Australia/22/2015 | WHO Collaborating Centre for Reference and Research on Influenza | Centers for Disease Control and Prevention | NA |
| EPI650066 | NS | Australia | 2015-Mar-12 | A/South Australia/22/2015 | WHO Collaborating Centre for Reference and Research on Influenza | Centers for Disease Control and Prevention | NA |
| EPI650067 | MP | Australia | 2015-Mar-12 | A/South Australia/22/2015 | WHO Collaborating Centre for Reference and Research on Influenza | Centers for Disease Control and Prevention | NA |
| EPI650068 | PA | Australia | 2015-Mar-12 | A/South Australia/22/2015 | WHO Collaborating Centre for Reference and Research on Influenza | Centers for Disease Control and Prevention | NA |
| EPI650069 | PB2 | Australia | 2015-Mar-12 | A/South Australia/22/2015 | WHO Collaborating Centre for Reference and Research on Influenza | Centers for Disease Control and Prevention | NA |
| EPI650070 | PB1 | Australia | 2015-Mar-12 | A/South Australia/22/2015 | WHO Collaborating Centre for Reference and Research on Influenza | Centers for Disease Control and Prevention | NA |
| EPI650071 | NA | Australia | 2015-Mar-12 | A/South Australia/22/2015 | WHO Collaborating Centre for Reference and Research on Influenza | Centers for Disease Control and Prevention | NA |
| EPI695507 | NP | South Africa | 2015-Aug-26 | A/South Africa/5325/2015 | National Institute for Communicable Disease | Centers for Disease Control and Prevention | NA |
| EPI695508 | NS | South Africa | 2015-Aug-26 | A/South Africa/5325/2015 | National Institute for Communicable Disease | Centers for Disease Control and Prevention | NA |
| EPI695509 | MP | South Africa | 2015-Aug-26 | A/South Africa/5325/2015 | National Institute for Communicable Disease | Centers for Disease Control and Prevention | NA |
| EPI695510 | PA | South Africa | 2015-Aug-26 | A/South Africa/5325/2015 | National Institute for Communicable Disease | Centers for Disease Control and Prevention | NA |
| EPI695511 | PB2 | South Africa | 2015-Aug-26 | A/South Africa/5325/2015 | National Institute for Communicable Disease | Centers for Disease Control and Prevention | NA |
| EPI695512 | PB1 | South Africa | 2015-Aug-26 | A/South Africa/5325/2015 | National Institute for Communicable Disease | Centers for Disease Control and Prevention | NA |
| EPI695513 | NA | South Africa | 2015-Aug-26 | A/South Africa/5325/2015 | National Institute for Communicable Disease | Centers for Disease Control and Prevention | NA |
| EPI695514 | HA | South Africa | 2015-Aug-26 | A/South Africa/5325/2015 | National Institute for Communicable Disease | Centers for Disease Control and Prevention | NA |
| EPI691354 | PA | Mali | 2015-Sep-28 | A/Mali/239 CI/2015 | NIC Lab CVD-MALI | Centers for Disease Control and Prevention | NA |
| EPI691355 | PB2 | Mali | 2015-Sep-28 | A/Mali/239 CI/2015 | NIC Lab CVD-MALI | Centers for Disease Control and Prevention | NA |
| EPI691356 | PB1 | Mali | 2015-Sep-28 | A/Mali/239 CI/2015 | NIC Lab CVD-MALI | Centers for Disease Control and Prevention | NA |
| EPI691351 | NP | Mali | 2015-Sep-28 | A/Mali/239 CI/2015 | NIC Lab CVD-MALI | Centers for Disease Control and Prevention | NA |
| EPI691352 | NS | Mali | 2015-Sep-28 | A/Mali/239 CI/2015 | NIC Lab CVD-MALI | Centers for Disease Control and Prevention | NA |
| EPI691353 | MP | Mali | 2015-Sep-28 | A/Mali/239 CI/2015 | NIC Lab CVD-MALI | Centers for Disease Control and Prevention | NA |
| EPI691357 | NA | Mali | 2015-Sep-28 | A/Mali/239 CI/2015 | NIC Lab CVD-MALI | Centers for Disease Control and Prevention | NA |
| EPI691358 | HA | Mali | 2015-Sep-28 | A/Mali/239 CI/2015 | NIC Lab CVD-MALI | Centers for Disease Control and Prevention | NA |
| EPI565041 | NP | Hong Kong (SAR) | 2014-Oct-13 | A/Hong Kong/7561/2014 | Government Virus Unit | Centers for Disease Control and Prevention | NA |
| EPI565042 | NS | Hong Kong (SAR) | 2014-Oct-13 | A/Hong Kong/7561/2014 | Government Virus Unit | Centers for Disease Control and Prevention | NA |
| EPI565043 | MP | Hong Kong (SAR) | 2014-Oct-13 | A/Hong Kong/7561/2014 | Government Virus Unit | Centers for Disease Control and Prevention | NA |
| EPI565044 | PA | Hong Kong (SAR) | 2014-Oct-13 | A/Hong Kong/7561/2014 | Government Virus Unit | Centers for Disease Control and Prevention | NA |
| EPI565045 | PB2 | Hong Kong (SAR) | 2014-Oct-13 | A/Hong Kong/7561/2014 | Government Virus Unit | Centers for Disease Control and Prevention | NA |
| EPI565046 | PB1 | Hong Kong (SAR) | 2014-Oct-13 | A/Hong Kong/7561/2014 | Government Virus Unit | Centers for Disease Control and Prevention | NA |
| EPI565047 | NA | Hong Kong (SAR) | 2014-Oct-13 | A/Hong Kong/7561/2014 | Government Virus Unit | Centers for Disease Control and Prevention | NA |
| EPI565048 | HA | Hong Kong (SAR) | 2014-Oct-13 | A/Hong Kong/7561/2014 | Government Virus Unit | Centers for Disease Control and Prevention | NA |
| EPI638389 | PB2 | Dominican Republic | 2015-May-11 | A/Dominican Republic/9251/2015 | Laboratorio de Investigacion / Centro de Educacion Medica y Amistad Dominico Japones (CEMADOJA) | Centers for Disease Control and Prevention | NA |
| EPI638390 | PB1 | Dominican Republic | 2015-May-11 | A/Dominican Republic/9251/2015 | Laboratorio de Investigacion / Centro de Educacion Medica y Amistad Dominico Japones (CEMADOJA) | Centers for Disease Control and Prevention | NA |
| EPI638385 | NP | Dominican Republic | 2015-May-11 | A/Dominican Republic/9251/2015 | Laboratorio de Investigacion / Centro de Educacion Medica y Amistad Dominico Japones (CEMADOJA) | Centers for Disease Control and Prevention | NA |
| EPI638386 | NS | Dominican Republic | 2015-May-11 | A/Dominican Republic/9251/2015 | Laboratorio de Investigacion / Centro de Educacion Medica y Amistad Dominico Japones (CEMADOJA) | Centers for Disease Control and Prevention | NA |
| EPI638387 | MP | Dominican Republic | 2015-May-11 | A/Dominican Republic/9251/2015 | Laboratorio de Investigacion / Centro de Educacion Medica y Amistad Dominico Japones (CEMADOJA) | Centers for Disease Control and Prevention | NA |
| EPI638388 | PA | Dominican Republic | 2015-May-11 | A/Dominican Republic/9251/2015 | Laboratorio de Investigacion / Centro de Educacion Medica y Amistad Dominico Japones (CEMADOJA) | Centers for Disease Control and Prevention | NA |
| EPI638391 | NA | Dominican Republic | 2015-May-11 | A/Dominican Republic/9251/2015 | Laboratorio de Investigacion / Centro de Educacion Medica y Amistad Dominico Japones (CEMADOJA) | Centers for Disease Control and Prevention | NA |
| EPI638392 | HA | Dominican Republic | 2015-May-11 | A/Dominican Republic/9251/2015 | Laboratorio de Investigacion / Centro de Educacion Medica y Amistad Dominico Japones (CEMADOJA) | Centers for Disease Control and Prevention | NA |
| EPI529411 | HA | Wallis and Futuna | 2013-Nov-19 | A/NEW CALEDONIA/58/2013 | Institut Pasteur New Caledonia | WHO Collaborating Centre for Reference and Research on Influenza | Deng,Y-M.; Iannello,P.; Spirason,N.; Jelley,L.; Lau,H.; Komadina,N. |
| EPI529404 | NP | Wallis and Futuna | 2013-Nov-19 | A/NEW CALEDONIA/58/2013 | Institut Pasteur New Caledonia | WHO Collaborating Centre for Reference and Research on Influenza | Deng,Y-M.; Iannello,P.; Spirason,N.; Jelley,L.; Lau,H.; Komadina,N. |
| EPI529405 | NS | Wallis and Futuna | 2013-Nov-19 | A/NEW CALEDONIA/58/2013 | Institut Pasteur New Caledonia | WHO Collaborating Centre for Reference and Research on Influenza | Deng,Y-M.; Iannello,P.; Spirason,N.; Jelley,L.; Lau,H.; Komadina,N. |
| EPI529406 | MP | Wallis and Futuna | 2013-Nov-19 | A/NEW CALEDONIA/58/2013 | Institut Pasteur New Caledonia | WHO Collaborating Centre for Reference and Research on Influenza | Deng,Y-M.; Iannello,P.; Spirason,N.; Jelley,L.; Lau,H.; Komadina,N. |
| EPI529407 | PA | Wallis and Futuna | 2013-Nov-19 | A/NEW CALEDONIA/58/2013 | Institut Pasteur New Caledonia | WHO Collaborating Centre for Reference and Research on Influenza | Deng,Y-M.; Iannello,P.; Spirason,N.; Jelley,L.; Lau,H.; Komadina,N. |
| EPI529408 | PB2 | Wallis and Futuna | 2013-Nov-19 | A/NEW CALEDONIA/58/2013 | Institut Pasteur New Caledonia | WHO Collaborating Centre for Reference and Research on Influenza | Deng,Y-M.; Iannello,P.; Spirason,N.; Jelley,L.; Lau,H.; Komadina,N. |
| EPI529409 | PB1 | Wallis and Futuna | 2013-Nov-19 | A/NEW CALEDONIA/58/2013 | Institut Pasteur New Caledonia | WHO Collaborating Centre for Reference and Research on Influenza | Deng,Y-M.; Iannello,P.; Spirason,N.; Jelley,L.; Lau,H.; Komadina,N. |
| EPI529410 | NA | Wallis and Futuna | 2013-Nov-19 | A/NEW CALEDONIA/58/2013 | Institut Pasteur New Caledonia | WHO Collaborating Centre for Reference and Research on Influenza | Deng,Y-M.; Iannello,P.; Spirason,N.; Jelley,L.; Lau,H.; Komadina,N. |
| EPI540916 | MP | Sri Lanka | 2014-May-03 | A/SRI LANKA/31/2014 | Medical Research Institute | WHO Collaborating Centre for Reference and Research on Influenza | Deng,Y-M.; Iannello,P.; Spirason,N.; Jelley,L.; Lau,H.; Komadina,N. |
| EPI540917 | NA | Sri Lanka | 2014-May-03 | A/SRI LANKA/31/2014 | Medical Research Institute | WHO Collaborating Centre for Reference and Research on Influenza | Deng,Y-M.; Iannello,P.; Spirason,N.; Jelley,L.; Lau,H.; Komadina,N. |
| EPI540918 | HA | Sri Lanka | 2014-May-03 | A/SRI LANKA/31/2014 | Medical Research Institute | WHO Collaborating Centre for Reference and Research on Influenza | Deng,Y-M.; Iannello,P.; Spirason,N.; Jelley,L.; Lau,H.; Komadina,N. |
| EPI561970 | NP | Sri Lanka | 2014-May-03 | A/SRI LANKA/31/2014 | Medical Research Institute | WHO Collaborating Centre for Reference and Research on Influenza | Deng,Y-M.; Iannello,P.; Spirason,N.; Jelley,L.; Lau,H.; Komadina,N. |
| EPI561971 | NS | Sri Lanka | 2014-May-03 | A/SRI LANKA/31/2014 | Medical Research Institute | WHO Collaborating Centre for Reference and Research on Influenza | Deng,Y-M.; Iannello,P.; Spirason,N.; Jelley,L.; Lau,H.; Komadina,N. |
| EPI561972 | PA | Sri Lanka | 2014-May-03 | A/SRI LANKA/31/2014 | Medical Research Institute | WHO Collaborating Centre for Reference and Research on Influenza | Deng,Y-M.; Iannello,P.; Spirason,N.; Jelley,L.; Lau,H.; Komadina,N. |
| EPI561973 | PB2 | Sri Lanka | 2014-May-03 | A/SRI LANKA/31/2014 | Medical Research Institute | WHO Collaborating Centre for Reference and Research on Influenza | Deng,Y-M.; Iannello,P.; Spirason,N.; Jelley,L.; Lau,H.; Komadina,N. |
| EPI561974 | PB1 | Sri Lanka | 2014-May-03 | A/SRI LANKA/31/2014 | Medical Research Institute | WHO Collaborating Centre for Reference and Research on Influenza | Deng,Y-M.; Iannello,P.; Spirason,N.; Jelley,L.; Lau,H.; Komadina,N. |
| EPI614644 | NS | Puerto Rico | 2014-Nov-14 | A/Puerto Rico/30/2014 | Puerto Rico Department of Health | Centers for Disease Control and Prevention | NA |
| EPI575879 | NP | Puerto Rico | 2014-Nov-14 | A/Puerto Rico/30/2014 | Puerto Rico Department of Health | Centers for Disease Control and Prevention | NA |
| EPI575880 | MP | Puerto Rico | 2014-Nov-14 | A/Puerto Rico/30/2014 | Puerto Rico Department of Health | Centers for Disease Control and Prevention | NA |
| EPI575881 | PA | Puerto Rico | 2014-Nov-14 | A/Puerto Rico/30/2014 | Puerto Rico Department of Health | Centers for Disease Control and Prevention | NA |
| EPI575882 | PB2 | Puerto Rico | 2014-Nov-14 | A/Puerto Rico/30/2014 | Puerto Rico Department of Health | Centers for Disease Control and Prevention | NA |
| EPI575883 | PB1 | Puerto Rico | 2014-Nov-14 | A/Puerto Rico/30/2014 | Puerto Rico Department of Health | Centers for Disease Control and Prevention | NA |
| EPI575884 | NA | Puerto Rico | 2014-Nov-14 | A/Puerto Rico/30/2014 | Puerto Rico Department of Health | Centers for Disease Control and Prevention | NA |
| EPI575885 | HA | Puerto Rico | 2014-Nov-14 | A/Puerto Rico/30/2014 | Puerto Rico Department of Health | Centers for Disease Control and Prevention | NA |
| EPI614496 | PB2 | South Africa | 2013-Jun-06 | A/South Africa/3626/2013 | National Institute for Medical Research | Centers for Disease Control and Prevention | NA |
| EPI614497 | PB1 | South Africa | 2013-Jun-06 | A/South Africa/3626/2013 | National Institute for Medical Research | Centers for Disease Control and Prevention | NA |
| EPI614492 | NP | South Africa | 2013-Jun-06 | A/South Africa/3626/2013 | National Institute for Medical Research | Centers for Disease Control and Prevention | NA |
| EPI614493 | NS | South Africa | 2013-Jun-06 | A/South Africa/3626/2013 | National Institute for Medical Research | Centers for Disease Control and Prevention | NA |
| EPI614494 | MP | South Africa | 2013-Jun-06 | A/South Africa/3626/2013 | National Institute for Medical Research | Centers for Disease Control and Prevention | NA |
| EPI614495 | PA | South Africa | 2013-Jun-06 | A/South Africa/3626/2013 | National Institute for Medical Research | Centers for Disease Control and Prevention | NA |
| EPI577030 | NA | South Africa | 2013-Jun-06 | A/South Africa/3626/2013 | National Institute for Medical Research | Centers for Disease Control and Prevention | NA |
| EPI577031 | HA | South Africa | 2013-Jun-06 | A/South Africa/3626/2013 | National Institute for Medical Research | Centers for Disease Control and Prevention | NA |
| EPI360008 | HA | Czech Republic | 2011-Oct-25 | A/Czech Republic/112/2011 |  | Other Database Import | Nagy,A.; Jirincova,H.; Havlickova,M. |
| EPI360009 | NP | Czech Republic | 2011-Oct-25 | A/Czech Republic/112/2011 |  | Other Database Import | Nagy,A.; Jirincova,H.; Havlickova,M. |
| EPI360015 | NS | Czech Republic | 2011-Oct-25 | A/Czech Republic/112/2011 |  | Other Database Import | Nagy,A.; Jirincova,H.; Havlickova,M. |
| EPI360011 | NA | Czech Republic | 2011-Oct-25 | A/Czech Republic/112/2011 |  | Other Database Import | Nagy,A.; Jirincova,H.; Havlickova,M. |
| EPI360013 | MP | Czech Republic | 2011-Oct-25 | A/Czech Republic/112/2011 |  | Other Database Import | Nagy,A.; Jirincova,H.; Havlickova,M. |
| EPI360004 | PB2 | Czech Republic | 2011-Oct-25 | A/Czech Republic/112/2011 |  | Other Database Import | Nagy,A.; Jirincova,H.; Havlickova,M. |
| EPI360005 | PB1 | Czech Republic | 2011-Oct-25 | A/Czech Republic/112/2011 |  | Other Database Import | Nagy,A.; Jirincova,H.; Havlickova,M. |
| EPI360006 | PA | Czech Republic | 2011-Oct-25 | A/Czech Republic/112/2011 |  | Other Database Import | Nagy,A.; Jirincova,H.; Havlickova,M. |
| EPI393767 | NP | United States | 2012-Apr-17 | A/New York/34/2012 | New York State Department of Health | Centers for Disease Control and Prevention | NA |
| EPI393768 | NS | United States | 2012-Apr-17 | A/New York/34/2012 | New York State Department of Health | Centers for Disease Control and Prevention | NA |
| EPI393769 | MP | United States | 2012-Apr-17 | A/New York/34/2012 | New York State Department of Health | Centers for Disease Control and Prevention | NA |
| EPI393770 | PA | United States | 2012-Apr-17 | A/New York/34/2012 | New York State Department of Health | Centers for Disease Control and Prevention | NA |
| EPI393771 | PB2 | United States | 2012-Apr-17 | A/New York/34/2012 | New York State Department of Health | Centers for Disease Control and Prevention | NA |
| EPI393772 | PB1 | United States | 2012-Apr-17 | A/New York/34/2012 | New York State Department of Health | Centers for Disease Control and Prevention | NA |
| EPI393773 | NA | United States | 2012-Apr-17 | A/New York/34/2012 | New York State Department of Health | Centers for Disease Control and Prevention | NA |
| EPI393774 | HA | United States | 2012-Apr-17 | A/New York/34/2012 | New York State Department of Health | Centers for Disease Control and Prevention | NA |
| EPI330850 | NP | China | 2011-Apr-02 | A/Fujian-Xinluo/SWL1141/2011 | WHO Chinese National Influenza Center | WHO Chinese National Influenza Center | Lan,Yu;Wang,Dayan;Li,Xiyan;Huang,Weijuan;Zhao,Xiang;Chen,Yaoyao;Yang,Lei;Shu,Yuelong |
| EPI330851 | NS | China | 2011-Apr-02 | A/Fujian-Xinluo/SWL1141/2011 | WHO Chinese National Influenza Center | WHO Chinese National Influenza Center | Lan,Yu;Wang,Dayan;Li,Xiyan;Huang,Weijuan;Zhao,Xiang;Chen,Yaoyao;Yang,Lei;Shu,Yuelong |
| EPI330852 | MP | China | 2011-Apr-02 | A/Fujian-Xinluo/SWL1141/2011 | WHO Chinese National Influenza Center | WHO Chinese National Influenza Center | Lan,Yu;Wang,Dayan;Li,Xiyan;Huang,Weijuan;Zhao,Xiang;Chen,Yaoyao;Yang,Lei;Shu,Yuelong |
| EPI330855 | PB1 | China | 2011-Apr-02 | A/Fujian-Xinluo/SWL1141/2011 | WHO Chinese National Influenza Center | WHO Chinese National Influenza Center | Lan,Yu;Wang,Dayan;Li,Xiyan;Huang,Weijuan;Zhao,Xiang;Chen,Yaoyao;Yang,Lei;Shu,Yuelong |
| EPI330856 | NA | China | 2011-Apr-02 | A/Fujian-Xinluo/SWL1141/2011 | WHO Chinese National Influenza Center | WHO Chinese National Influenza Center | Lan,Yu;Wang,Dayan;Li,Xiyan;Huang,Weijuan;Zhao,Xiang;Chen,Yaoyao;Yang,Lei;Shu,Yuelong |
| EPI330857 | HA | China | 2011-Apr-02 | A/Fujian-Xinluo/SWL1141/2011 | WHO Chinese National Influenza Center | WHO Chinese National Influenza Center | Lan,Yu;Wang,Dayan;Li,Xiyan;Huang,Weijuan;Zhao,Xiang;Chen,Yaoyao;Yang,Lei;Shu,Yuelong |
| EPI330853 | PA | China | 2011-Apr-02 | A/Fujian-Xinluo/SWL1141/2011 | WHO Chinese National Influenza Center | WHO Chinese National Influenza Center | Lan,Yu;Wang,Dayan;Li,Xiyan;Huang,Weijuan;Zhao,Xiang;Chen,Yaoyao;Yang,Lei;Shu,Yuelong |
| EPI330854 | PB2 | China | 2011-Apr-02 | A/Fujian-Xinluo/SWL1141/2011 | WHO Chinese National Influenza Center | WHO Chinese National Influenza Center | Lan,Yu;Wang,Dayan;Li,Xiyan;Huang,Weijuan;Zhao,Xiang;Chen,Yaoyao;Yang,Lei;Shu,Yuelong |
| EPI567850 | PB1 | Vietnam | 2013-Aug-03 | A/Vietnam/13V H1-1/2013 |  | Niigata University | Saito, Reiko; Takemae, Nobuhiro; Saito, Takehiko; Shobugawa, Yugo; Kondo, Hiroki; Hibino, Akinobu |
| EPI567852 | MP | Vietnam | 2013-Aug-03 | A/Vietnam/13V H1-1/2013 |  | Niigata University | Saito, Reiko; Takemae, Nobuhiro; Saito, Takehiko; Shobugawa, Yugo; Kondo, Hiroki; Hibino, Akinobu |
| EPI567851 | HA | Vietnam | 2013-Aug-03 | A/Vietnam/13V H1-1/2013 |  | Niigata University | Saito, Reiko; Takemae, Nobuhiro; Saito, Takehiko; Shobugawa, Yugo; Kondo, Hiroki; Hibino, Akinobu |
| EPI567856 | NP | Vietnam | 2013-Aug-03 | A/Vietnam/13V H1-1/2013 |  | Niigata University | Saito, Reiko; Takemae, Nobuhiro; Saito, Takehiko; Shobugawa, Yugo; Kondo, Hiroki; Hibino, Akinobu |
| EPI567853 | PB2 | Vietnam | 2013-Aug-03 | A/Vietnam/13V H1-1/2013 |  | Niigata University | Saito, Reiko; Takemae, Nobuhiro; Saito, Takehiko; Shobugawa, Yugo; Kondo, Hiroki; Hibino, Akinobu |
| EPI567849 | PA | Vietnam | 2013-Aug-03 | A/Vietnam/13V H1-1/2013 |  | Niigata University | Saito, Reiko; Takemae, Nobuhiro; Saito, Takehiko; Shobugawa, Yugo; Kondo, Hiroki; Hibino, Akinobu |
| EPI567854 | NA | Vietnam | 2013-Aug-03 | A/Vietnam/13V H1-1/2013 |  | Niigata University | Saito, Reiko; Takemae, Nobuhiro; Saito, Takehiko; Shobugawa, Yugo; Kondo, Hiroki; Hibino, Akinobu |
| EPI567855 | NS | Vietnam | 2013-Aug-03 | A/Vietnam/13V H1-1/2013 |  | Niigata University | Saito, Reiko; Takemae, Nobuhiro; Saito, Takehiko; Shobugawa, Yugo; Kondo, Hiroki; Hibino, Akinobu |
| EPI247059 | PB2 | India | 2009-May-01 | A/Hyd/NIV51/2009 |  | Other Database Import | Mishra,A.; Potdar,V.; Chadha,M.; Jadhav,S.; Mullick,J.; Cherian,S. |
| EPI247060 | PB1 | India | 2009-May-01 | A/Hyd/NIV51/2009 |  | Other Database Import | Mishra,A.; Potdar,V.; Chadha,M.; Jadhav,S.; Mullick,J.; Cherian,S. |
| EPI247061 | PA | India | 2009-May-01 | A/Hyd/NIV51/2009 |  | Other Database Import | Mishra,A.; Potdar,V.; Chadha,M.; Jadhav,S.; Mullick,J.; Cherian,S. |
| EPI247062 | HA | India | 2009-May-01 | A/Hyd/NIV51/2009 |  | Other Database Import | Mishra,A.; Potdar,V.; Chadha,M.; Jadhav,S.; Mullick,J.; Cherian,S. |
| EPI247063 | NP | India | 2009-May-01 | A/Hyd/NIV51/2009 |  | Other Database Import | Mishra,A.; Potdar,V.; Chadha,M.; Jadhav,S.; Mullick,J.; Cherian,S. |
| EPI247064 | NA | India | 2009-May-01 | A/Hyd/NIV51/2009 |  | Other Database Import | Mishra,A.; Potdar,V.; Chadha,M.; Jadhav,S.; Mullick,J.; Cherian,S. |
| EPI247065 | MP | India | 2009-May-01 | A/Hyd/NIV51/2009 |  | Other Database Import | Mishra,A.; Potdar,V.; Chadha,M.; Jadhav,S.; Mullick,J.; Cherian,S. |
| EPI247066 | NS | India | 2009-May-01 | A/Hyd/NIV51/2009 |  | Other Database Import | Mishra,A.; Potdar,V.; Chadha,M.; Jadhav,S.; Mullick,J.; Cherian,S. |
| EPI278889 | NP | United States | 2010-Jun-01 | A/Texas/03/2010 | Texas Department of State Health Services-Laboratory Services | Centers for Disease Control and Prevention | NA |
| EPI278890 | NS | United States | 2010-Jun-01 | A/Texas/03/2010 | Texas Department of State Health Services-Laboratory Services | Centers for Disease Control and Prevention | NA |
| EPI278891 | MP | United States | 2010-Jun-01 | A/Texas/03/2010 | Texas Department of State Health Services-Laboratory Services | Centers for Disease Control and Prevention | NA |
| EPI278892 | PA | United States | 2010-Jun-01 | A/Texas/03/2010 | Texas Department of State Health Services-Laboratory Services | Centers for Disease Control and Prevention | NA |
| EPI278893 | PB2 | United States | 2010-Jun-01 | A/Texas/03/2010 | Texas Department of State Health Services-Laboratory Services | Centers for Disease Control and Prevention | NA |
| EPI278894 | PB1 | United States | 2010-Jun-01 | A/Texas/03/2010 | Texas Department of State Health Services-Laboratory Services | Centers for Disease Control and Prevention | NA |
| EPI278895 | NA | United States | 2010-Jun-01 | A/Texas/03/2010 | Texas Department of State Health Services-Laboratory Services | Centers for Disease Control and Prevention | NA |
| EPI278896 | HA | United States | 2010-Jun-01 | A/Texas/03/2010 | Texas Department of State Health Services-Laboratory Services | Centers for Disease Control and Prevention | NA |
| EPI390385 | HA | Russian Federation | 2011-Aug-01 | A/St. Petersburg/100/2011 |  | Other Database Import | Wentworth,D.E.; Dugan,V.; Halpin,R.; Lin,X.; Bera,J.; Ghedin,E.; Fedorova,N.; Overton,L.; Tsitrin,T.; Stockwell,T.; Amedeo,P.; Bishop,B.; Chen,H.; Edworthy,P.; Gupta,N.; Katzel,D.; Li,K.; Schobel,S.; Shrivastava,S.; Thovarai,V.; Wang,S.; Ramanunninair,M.; Silverman,J.; Devis,R.; Phan,L.; Le,J.; Pokorny,B.A.; Onodera,S.; Fulvini,A.A.; He,Y.; Kilbourne,E.D.; Bucher,D.; Bao,Y.; Sanders,R.; Dernovoy,D.; Kiryutin,B.; Lipman,D.J.; Tatusova,T. |
| EPI390386 | MP | Russian Federation | 2011-Aug-01 | A/St. Petersburg/100/2011 |  | Other Database Import | Wentworth,D.E.; Dugan,V.; Halpin,R.; Lin,X.; Bera,J.; Ghedin,E.; Fedorova,N.; Overton,L.; Tsitrin,T.; Stockwell,T.; Amedeo,P.; Bishop,B.; Chen,H.; Edworthy,P.; Gupta,N.; Katzel,D.; Li,K.; Schobel,S.; Shrivastava,S.; Thovarai,V.; Wang,S.; Ramanunninair,M.; Silverman,J.; Devis,R.; Phan,L.; Le,J.; Pokorny,B.A.; Onodera,S.; Fulvini,A.A.; He,Y.; Kilbourne,E.D.; Bucher,D.; Bao,Y.; Sanders,R.; Dernovoy,D.; Kiryutin,B.; Lipman,D.J.; Tatusova,T. |
| EPI390387 | NA | Russian Federation | 2011-Aug-01 | A/St. Petersburg/100/2011 |  | Other Database Import | Wentworth,D.E.; Dugan,V.; Halpin,R.; Lin,X.; Bera,J.; Ghedin,E.; Fedorova,N.; Overton,L.; Tsitrin,T.; Stockwell,T.; Amedeo,P.; Bishop,B.; Chen,H.; Edworthy,P.; Gupta,N.; Katzel,D.; Li,K.; Schobel,S.; Shrivastava,S.; Thovarai,V.; Wang,S.; Ramanunninair,M.; Silverman,J.; Devis,R.; Phan,L.; Le,J.; Pokorny,B.A.; Onodera,S.; Fulvini,A.A.; He,Y.; Kilbourne,E.D.; Bucher,D.; Bao,Y.; Sanders,R.; Dernovoy,D.; Kiryutin,B.; Lipman,D.J.; Tatusova,T. |
| EPI390388 | NP | Russian Federation | 2011-Aug-01 | A/St. Petersburg/100/2011 |  | Other Database Import | Wentworth,D.E.; Dugan,V.; Halpin,R.; Lin,X.; Bera,J.; Ghedin,E.; Fedorova,N.; Overton,L.; Tsitrin,T.; Stockwell,T.; Amedeo,P.; Bishop,B.; Chen,H.; Edworthy,P.; Gupta,N.; Katzel,D.; Li,K.; Schobel,S.; Shrivastava,S.; Thovarai,V.; Wang,S.; Ramanunninair,M.; Silverman,J.; Devis,R.; Phan,L.; Le,J.; Pokorny,B.A.; Onodera,S.; Fulvini,A.A.; He,Y.; Kilbourne,E.D.; Bucher,D.; Bao,Y.; Sanders,R.; Dernovoy,D.; Kiryutin,B.; Lipman,D.J.; Tatusova,T. |
| EPI390389 | NS | Russian Federation | 2011-Aug-01 | A/St. Petersburg/100/2011 |  | Other Database Import | Wentworth,D.E.; Dugan,V.; Halpin,R.; Lin,X.; Bera,J.; Ghedin,E.; Fedorova,N.; Overton,L.; Tsitrin,T.; Stockwell,T.; Amedeo,P.; Bishop,B.; Chen,H.; Edworthy,P.; Gupta,N.; Katzel,D.; Li,K.; Schobel,S.; Shrivastava,S.; Thovarai,V.; Wang,S.; Ramanunninair,M.; Silverman,J.; Devis,R.; Phan,L.; Le,J.; Pokorny,B.A.; Onodera,S.; Fulvini,A.A.; He,Y.; Kilbourne,E.D.; Bucher,D.; Bao,Y.; Sanders,R.; Dernovoy,D.; Kiryutin,B.; Lipman,D.J.; Tatusova,T. |
| EPI390390 | PA | Russian Federation | 2011-Aug-01 | A/St. Petersburg/100/2011 |  | Other Database Import | Wentworth,D.E.; Dugan,V.; Halpin,R.; Lin,X.; Bera,J.; Ghedin,E.; Fedorova,N.; Overton,L.; Tsitrin,T.; Stockwell,T.; Amedeo,P.; Bishop,B.; Chen,H.; Edworthy,P.; Gupta,N.; Katzel,D.; Li,K.; Schobel,S.; Shrivastava,S.; Thovarai,V.; Wang,S.; Ramanunninair,M.; Silverman,J.; Devis,R.; Phan,L.; Le,J.; Pokorny,B.A.; Onodera,S.; Fulvini,A.A.; He,Y.; Kilbourne,E.D.; Bucher,D.; Bao,Y.; Sanders,R.; Dernovoy,D.; Kiryutin,B.; Lipman,D.J.; Tatusova,T. |
| EPI390391 | PB1 | Russian Federation | 2011-Aug-01 | A/St. Petersburg/100/2011 |  | Other Database Import | Wentworth,D.E.; Dugan,V.; Halpin,R.; Lin,X.; Bera,J.; Ghedin,E.; Fedorova,N.; Overton,L.; Tsitrin,T.; Stockwell,T.; Amedeo,P.; Bishop,B.; Chen,H.; Edworthy,P.; Gupta,N.; Katzel,D.; Li,K.; Schobel,S.; Shrivastava,S.; Thovarai,V.; Wang,S.; Ramanunninair,M.; Silverman,J.; Devis,R.; Phan,L.; Le,J.; Pokorny,B.A.; Onodera,S.; Fulvini,A.A.; He,Y.; Kilbourne,E.D.; Bucher,D.; Bao,Y.; Sanders,R.; Dernovoy,D.; Kiryutin,B.; Lipman,D.J.; Tatusova,T. |
| EPI390392 | PB2 | Russian Federation | 2011-Aug-01 | A/St. Petersburg/100/2011 |  | Other Database Import | Wentworth,D.E.; Dugan,V.; Halpin,R.; Lin,X.; Bera,J.; Ghedin,E.; Fedorova,N.; Overton,L.; Tsitrin,T.; Stockwell,T.; Amedeo,P.; Bishop,B.; Chen,H.; Edworthy,P.; Gupta,N.; Katzel,D.; Li,K.; Schobel,S.; Shrivastava,S.; Thovarai,V.; Wang,S.; Ramanunninair,M.; Silverman,J.; Devis,R.; Phan,L.; Le,J.; Pokorny,B.A.; Onodera,S.; Fulvini,A.A.; He,Y.; Kilbourne,E.D.; Bucher,D.; Bao,Y.; Sanders,R.; Dernovoy,D.; Kiryutin,B.; Lipman,D.J.; Tatusova,T. |
|  |  |  |  |  |  |  | NA |
|  |  |  |  |  |  |  | NA |
| all 119 strains included except brunei/01, A_India_Gwl-01_2011 and PB1 of wisconsin 88 |  |  |  |  |  |  | NA |
|  |  |  |  |  |  |  | NA |
|  |  |  |  |  |  |  | NA |
| Complete genome from MCC Tree |  |  |  |  |  |  | NA |
|  |  |  |  |  |  |  | NA |
|  |  |  |  |  |  |  | NA |
| EPI630416 | NA | India | 2015-Mar-30 | A/India/P158900/2015 | National Institute of Virology | National Institute of Infectious Diseases (NIID) | Takashita,Emi; Fujisaki,Seiichiro; Shirakura,Masayuki; Watanabe,Shinji; Odagiri,Takato |
| EPI630417 | HA | India | 2015-Mar-30 | A/India/P158900/2015 | National Institute of Virology | National Institute of Infectious Diseases (NIID) | Takashita,Emi; Fujisaki,Seiichiro; Shirakura,Masayuki; Watanabe,Shinji; Odagiri,Takato |
| EPI635511 | NP | India | 2015-Mar-30 | A/India/P158900/2015 | National Institute of Virology | National Institute of Infectious Diseases (NIID) | Takashita,Emi; Fujisaki,Seiichiro; Shirakura,Masayuki; Watanabe,Shinji; Odagiri,Takato |
| EPI635512 | NS | India | 2015-Mar-30 | A/India/P158900/2015 | National Institute of Virology | National Institute of Infectious Diseases (NIID) | Takashita,Emi; Fujisaki,Seiichiro; Shirakura,Masayuki; Watanabe,Shinji; Odagiri,Takato |
| EPI635513 | MP | India | 2015-Mar-30 | A/India/P158900/2015 | National Institute of Virology | National Institute of Infectious Diseases (NIID) | Takashita,Emi; Fujisaki,Seiichiro; Shirakura,Masayuki; Watanabe,Shinji; Odagiri,Takato |
| EPI635514 | PA | India | 2015-Mar-30 | A/India/P158900/2015 | National Institute of Virology | National Institute of Infectious Diseases (NIID) | Takashita,Emi; Fujisaki,Seiichiro; Shirakura,Masayuki; Watanabe,Shinji; Odagiri,Takato |
| EPI635515 | PB2 | India | 2015-Mar-30 | A/India/P158900/2015 | National Institute of Virology | National Institute of Infectious Diseases (NIID) | Takashita,Emi; Fujisaki,Seiichiro; Shirakura,Masayuki; Watanabe,Shinji; Odagiri,Takato |
| EPI635516 | PB1 | India | 2015-Mar-30 | A/India/P158900/2015 | National Institute of Virology | National Institute of Infectious Diseases (NIID) | Takashita,Emi; Fujisaki,Seiichiro; Shirakura,Masayuki; Watanabe,Shinji; Odagiri,Takato |
|  |  |  |  |  |  |  | NA |
| EPI626137 | PB2 | Bangladesh | 2015-May-10 | A/Bangladesh/01/2015 | Institute of Epidemiology Disease Control and Research (IEDCR) & Bangladesh National Influenza Centre (NIC) | Centers for Disease Control and Prevention | NA |
| EPI626138 | PB1 | Bangladesh | 2015-May-10 | A/Bangladesh/01/2015 | Institute of Epidemiology Disease Control and Research (IEDCR) & Bangladesh National Influenza Centre (NIC) | Centers for Disease Control and Prevention | NA |
| EPI626133 | NP | Bangladesh | 2015-May-10 | A/Bangladesh/01/2015 | Institute of Epidemiology Disease Control and Research (IEDCR) & Bangladesh National Influenza Centre (NIC) | Centers for Disease Control and Prevention | NA |
| EPI626134 | NS | Bangladesh | 2015-May-10 | A/Bangladesh/01/2015 | Institute of Epidemiology Disease Control and Research (IEDCR) & Bangladesh National Influenza Centre (NIC) | Centers for Disease Control and Prevention | NA |
| EPI626135 | MP | Bangladesh | 2015-May-10 | A/Bangladesh/01/2015 | Institute of Epidemiology Disease Control and Research (IEDCR) & Bangladesh National Influenza Centre (NIC) | Centers for Disease Control and Prevention | NA |
| EPI626136 | PA | Bangladesh | 2015-May-10 | A/Bangladesh/01/2015 | Institute of Epidemiology Disease Control and Research (IEDCR) & Bangladesh National Influenza Centre (NIC) | Centers for Disease Control and Prevention | NA |
| EPI626139 | NA | Bangladesh | 2015-May-10 | A/Bangladesh/01/2015 | Institute of Epidemiology Disease Control and Research (IEDCR) & Bangladesh National Influenza Centre (NIC) | Centers for Disease Control and Prevention | NA |
| EPI626140 | HA | Bangladesh | 2015-May-10 | A/Bangladesh/01/2015 | Institute of Epidemiology Disease Control and Research (IEDCR) & Bangladesh National Influenza Centre (NIC) | Centers for Disease Control and Prevention | NA |
|  |  |  |  |  |  |  | NA |
| EPI638389 | PB2 | Dominican Republic | 2015-May-11 | A/Dominican Republic/9251/2015 | Laboratorio de Investigacion / Centro de Educacion Medica y Amistad Dominico Japones (CEMADOJA) | Centers for Disease Control and Prevention | NA |
| EPI638390 | PB1 | Dominican Republic | 2015-May-11 | A/Dominican Republic/9251/2015 | Laboratorio de Investigacion / Centro de Educacion Medica y Amistad Dominico Japones (CEMADOJA) | Centers for Disease Control and Prevention | NA |
| EPI638385 | NP | Dominican Republic | 2015-May-11 | A/Dominican Republic/9251/2015 | Laboratorio de Investigacion / Centro de Educacion Medica y Amistad Dominico Japones (CEMADOJA) | Centers for Disease Control and Prevention | NA |
| EPI638386 | NS | Dominican Republic | 2015-May-11 | A/Dominican Republic/9251/2015 | Laboratorio de Investigacion / Centro de Educacion Medica y Amistad Dominico Japones (CEMADOJA) | Centers for Disease Control and Prevention | NA |
| EPI638387 | MP | Dominican Republic | 2015-May-11 | A/Dominican Republic/9251/2015 | Laboratorio de Investigacion / Centro de Educacion Medica y Amistad Dominico Japones (CEMADOJA) | Centers for Disease Control and Prevention | NA |
| EPI638388 | PA | Dominican Republic | 2015-May-11 | A/Dominican Republic/9251/2015 | Laboratorio de Investigacion / Centro de Educacion Medica y Amistad Dominico Japones (CEMADOJA) | Centers for Disease Control and Prevention | NA |
| EPI638391 | NA | Dominican Republic | 2015-May-11 | A/Dominican Republic/9251/2015 | Laboratorio de Investigacion / Centro de Educacion Medica y Amistad Dominico Japones (CEMADOJA) | Centers for Disease Control and Prevention | NA |
| EPI638392 | HA | Dominican Republic | 2015-May-11 | A/Dominican Republic/9251/2015 | Laboratorio de Investigacion / Centro de Educacion Medica y Amistad Dominico Japones (CEMADOJA) | Centers for Disease Control and Prevention | NA |
| EPI695457 | PA | Brazil | 2015-Sep-04 | A/Brazil/69221/2015 | Instituto Adolfo Lutz | Centers for Disease Control and Prevention | NA |
| EPI695458 | PB2 | Brazil | 2015-Sep-04 | A/Brazil/69221/2015 | Instituto Adolfo Lutz | Centers for Disease Control and Prevention | NA |
| EPI695459 | PB1 | Brazil | 2015-Sep-04 | A/Brazil/69221/2015 | Instituto Adolfo Lutz | Centers for Disease Control and Prevention | NA |
| EPI695454 | NP | Brazil | 2015-Sep-04 | A/Brazil/69221/2015 | Instituto Adolfo Lutz | Centers for Disease Control and Prevention | NA |
| EPI695455 | NS | Brazil | 2015-Sep-04 | A/Brazil/69221/2015 | Instituto Adolfo Lutz | Centers for Disease Control and Prevention | NA |
| EPI695456 | MP | Brazil | 2015-Sep-04 | A/Brazil/69221/2015 | Instituto Adolfo Lutz | Centers for Disease Control and Prevention | NA |
| EPI695460 | NA | Brazil | 2015-Sep-04 | A/Brazil/69221/2015 | Instituto Adolfo Lutz | Centers for Disease Control and Prevention | NA |
| EPI695461 | HA | Brazil | 2015-Sep-04 | A/Brazil/69221/2015 | Instituto Adolfo Lutz | Centers for Disease Control and Prevention | NA |
| EPI680459 | PB2 | Pakistan | 2015-Nov-14 | A/Pakistan/336/2015 | National Institute of Health | Centers for Disease Control and Prevention | NA |
| EPI680460 | PB1 | Pakistan | 2015-Nov-14 | A/Pakistan/336/2015 | National Institute of Health | Centers for Disease Control and Prevention | NA |
| EPI680455 | NP | Pakistan | 2015-Nov-14 | A/Pakistan/336/2015 | National Institute of Health | Centers for Disease Control and Prevention | NA |
| EPI680456 | NS | Pakistan | 2015-Nov-14 | A/Pakistan/336/2015 | National Institute of Health | Centers for Disease Control and Prevention | NA |
| EPI680457 | MP | Pakistan | 2015-Nov-14 | A/Pakistan/336/2015 | National Institute of Health | Centers for Disease Control and Prevention | NA |
| EPI680458 | PA | Pakistan | 2015-Nov-14 | A/Pakistan/336/2015 | National Institute of Health | Centers for Disease Control and Prevention | NA |
| EPI680461 | NA | Pakistan | 2015-Nov-14 | A/Pakistan/336/2015 | National Institute of Health | Centers for Disease Control and Prevention | NA |
| EPI680462 | HA | Pakistan | 2015-Nov-14 | A/Pakistan/336/2015 | National Institute of Health | Centers for Disease Control and Prevention | NA |
| EPI691359 | NP | Taiwan | 2015-Sep-05 | A/Taiwan/1036/2015 | ADImmune Corporation | Centers for Disease Control and Prevention | NA |
| EPI691360 | NS | Taiwan | 2015-Sep-05 | A/Taiwan/1036/2015 | ADImmune Corporation | Centers for Disease Control and Prevention | NA |
| EPI691361 | MP | Taiwan | 2015-Sep-05 | A/Taiwan/1036/2015 | ADImmune Corporation | Centers for Disease Control and Prevention | NA |
| EPI691362 | PA | Taiwan | 2015-Sep-05 | A/Taiwan/1036/2015 | ADImmune Corporation | Centers for Disease Control and Prevention | NA |
| EPI691363 | PB2 | Taiwan | 2015-Sep-05 | A/Taiwan/1036/2015 | ADImmune Corporation | Centers for Disease Control and Prevention | NA |
| EPI691364 | PB1 | Taiwan | 2015-Sep-05 | A/Taiwan/1036/2015 | ADImmune Corporation | Centers for Disease Control and Prevention | NA |
| EPI691365 | NA | Taiwan | 2015-Sep-05 | A/Taiwan/1036/2015 | ADImmune Corporation | Centers for Disease Control and Prevention | NA |
| EPI691366 | HA | Taiwan | 2015-Sep-05 | A/Taiwan/1036/2015 | ADImmune Corporation | Centers for Disease Control and Prevention | NA |
| EPI672206 | PB2 | United States | 2015-Sep-12 | A/California/90/2015 | California Department of Health Services | Centers for Disease Control and Prevention | NA |
| EPI672207 | PB1 | United States | 2015-Sep-12 | A/California/90/2015 | California Department of Health Services | Centers for Disease Control and Prevention | NA |
| EPI672202 | NP | United States | 2015-Sep-12 | A/California/90/2015 | California Department of Health Services | Centers for Disease Control and Prevention | NA |
| EPI672203 | NS | United States | 2015-Sep-12 | A/California/90/2015 | California Department of Health Services | Centers for Disease Control and Prevention | NA |
| EPI672204 | MP | United States | 2015-Sep-12 | A/California/90/2015 | California Department of Health Services | Centers for Disease Control and Prevention | NA |
| EPI672205 | PA | United States | 2015-Sep-12 | A/California/90/2015 | California Department of Health Services | Centers for Disease Control and Prevention | NA |
| EPI672208 | NA | United States | 2015-Sep-12 | A/California/90/2015 | California Department of Health Services | Centers for Disease Control and Prevention | NA |
| EPI672209 | HA | United States | 2015-Sep-12 | A/California/90/2015 | California Department of Health Services | Centers for Disease Control and Prevention | NA |
| EPI691214 | NP | United States | 2015-Dec-19 | A/Florida/99/2015 | Florida Department of Health-Tampa | Centers for Disease Control and Prevention | NA |
| EPI691215 | NS | United States | 2015-Dec-19 | A/Florida/99/2015 | Florida Department of Health-Tampa | Centers for Disease Control and Prevention | NA |
| EPI691216 | MP | United States | 2015-Dec-19 | A/Florida/99/2015 | Florida Department of Health-Tampa | Centers for Disease Control and Prevention | NA |
| EPI691217 | PA | United States | 2015-Dec-19 | A/Florida/99/2015 | Florida Department of Health-Tampa | Centers for Disease Control and Prevention | NA |
| EPI691218 | PB2 | United States | 2015-Dec-19 | A/Florida/99/2015 | Florida Department of Health-Tampa | Centers for Disease Control and Prevention | NA |
| EPI691219 | PB1 | United States | 2015-Dec-19 | A/Florida/99/2015 | Florida Department of Health-Tampa | Centers for Disease Control and Prevention | NA |
| EPI691220 | NA | United States | 2015-Dec-19 | A/Florida/99/2015 | Florida Department of Health-Tampa | Centers for Disease Control and Prevention | NA |
| EPI691221 | HA | United States | 2015-Dec-19 | A/Florida/99/2015 | Florida Department of Health-Tampa | Centers for Disease Control and Prevention | NA |
| EPI693486 | PA | Nepal | 2015-Sep-09 | A/Nepal/3375/2015 | National Public Health Laboratory | Centers for Disease Control and Prevention | NA |
| EPI693483 | NP | Nepal | 2015-Sep-09 | A/Nepal/3375/2015 | National Public Health Laboratory | Centers for Disease Control and Prevention | NA |
| EPI693484 | NS | Nepal | 2015-Sep-09 | A/Nepal/3375/2015 | National Public Health Laboratory | Centers for Disease Control and Prevention | NA |
| EPI693485 | MP | Nepal | 2015-Sep-09 | A/Nepal/3375/2015 | National Public Health Laboratory | Centers for Disease Control and Prevention | NA |
| EPI693487 | PB2 | Nepal | 2015-Sep-09 | A/Nepal/3375/2015 | National Public Health Laboratory | Centers for Disease Control and Prevention | NA |
| EPI693488 | PB1 | Nepal | 2015-Sep-09 | A/Nepal/3375/2015 | National Public Health Laboratory | Centers for Disease Control and Prevention | NA |
| EPI693489 | NA | Nepal | 2015-Sep-09 | A/Nepal/3375/2015 | National Public Health Laboratory | Centers for Disease Control and Prevention | NA |
| EPI693490 | HA | Nepal | 2015-Sep-09 | A/Nepal/3375/2015 | National Public Health Laboratory | Centers for Disease Control and Prevention | NA |
| EPI687126 | NP | Japan | 2015-Mar-16 | A/Fukuoka/SDC1/2015 | Virus Research Center, Sendai Medical Center | National Institute of Infectious Diseases (NIID) | Takashita,Emi; Fujisaki,Seiichiro; Shirakura,Masayuki; Watanabe,Shinji; Odagiri,Takato |
| EPI687127 | NS | Japan | 2015-Mar-16 | A/Fukuoka/SDC1/2015 | Virus Research Center, Sendai Medical Center | National Institute of Infectious Diseases (NIID) | Takashita,Emi; Fujisaki,Seiichiro; Shirakura,Masayuki; Watanabe,Shinji; Odagiri,Takato |
| EPI687128 | MP | Japan | 2015-Mar-16 | A/Fukuoka/SDC1/2015 | Virus Research Center, Sendai Medical Center | National Institute of Infectious Diseases (NIID) | Takashita,Emi; Fujisaki,Seiichiro; Shirakura,Masayuki; Watanabe,Shinji; Odagiri,Takato |
| EPI687129 | PA | Japan | 2015-Mar-16 | A/Fukuoka/SDC1/2015 | Virus Research Center, Sendai Medical Center | National Institute of Infectious Diseases (NIID) | Takashita,Emi; Fujisaki,Seiichiro; Shirakura,Masayuki; Watanabe,Shinji; Odagiri,Takato |
| EPI687130 | PB2 | Japan | 2015-Mar-16 | A/Fukuoka/SDC1/2015 | Virus Research Center, Sendai Medical Center | National Institute of Infectious Diseases (NIID) | Takashita,Emi; Fujisaki,Seiichiro; Shirakura,Masayuki; Watanabe,Shinji; Odagiri,Takato |
| EPI687131 | PB1 | Japan | 2015-Mar-16 | A/Fukuoka/SDC1/2015 | Virus Research Center, Sendai Medical Center | National Institute of Infectious Diseases (NIID) | Takashita,Emi; Fujisaki,Seiichiro; Shirakura,Masayuki; Watanabe,Shinji; Odagiri,Takato |
| EPI586589 | NA | Japan | 2015-Mar-16 | A/Fukuoka/SDC1/2015 | Virus Research Center, Sendai Medical Center | National Institute of Infectious Diseases (NIID) | Takashita,Emi; Fujisaki,Seiichiro; Shirakura,Masayuki; Watanabe,Shinji; Odagiri,Takato |
| EPI586590 | HA | Japan | 2015-Mar-16 | A/Fukuoka/SDC1/2015 | Virus Research Center, Sendai Medical Center | National Institute of Infectious Diseases (NIID) | Takashita,Emi; Fujisaki,Seiichiro; Shirakura,Masayuki; Watanabe,Shinji; Odagiri,Takato |
| EPI584985 | NP | India | 2015-Jan-27 | A/India/1268/2015 | National Institute of Virology | Centers for Disease Control and Prevention |  |
| EPI584987 | NS | India | 2015-Jan-27 | A/India/1268/2015 | National Institute of Virology | Centers for Disease Control and Prevention | NA |
| EPI584988 | MP | India | 2015-Jan-27 | A/India/1268/2015 | National Institute of Virology | Centers for Disease Control and Prevention | NA |
| EPI584989 | PA | India | 2015-Jan-27 | A/India/1268/2015 | National Institute of Virology | Centers for Disease Control and Prevention | NA |
| EPI584990 | PB2 | India | 2015-Jan-27 | A/India/1268/2015 | National Institute of Virology | Centers for Disease Control and Prevention | NA |
| EPI584991 | PB1 | India | 2015-Jan-27 | A/India/1268/2015 | National Institute of Virology | Centers for Disease Control and Prevention | NA |
| EPI584992 | NA | India | 2015-Jan-27 | A/India/1268/2015 | National Institute of Virology | Centers for Disease Control and Prevention | NA |
