## Supplementary material for "Next-Gen sequencing of novel pandemic swine flu [A(H1N1)pdm09] virus in India revealed novel mutations across the genome": Spacial diffusion of A(H1N1)pdm09 viruses between geographical territories with bayes factor

**Table S2:	Spacial diffusion of A(H1N1)pdm09 viruses between geographical territories with bayes factor**

| **Sl No** | **Bayes Factor** | **Transition between geographical territories with their longitude and altitude** |
| --- | --- | --- |
| 1 | 36555.5343265314 | between Africa (long: 34.508522; lat: -8.783195) and Europe (long: 15.255119; lat: 54.525963) |
| 2 | 1635.417293719311 | between Central Asia (long: 68.8319; lat: 45.450687) and Middle East (long: 42.55096; lat: 29.298529) |
| 3 | 1466.0939575567506 | between South America (long: -55.491478; lat: -8.783195) and South Asia (long: 76.45631; lat: 25.03764) |
| 4 | 1304.3207582213108 | between Africa (long: 34.508522; lat: -8.783195) and North America (long: -105.25512; lat: 54.525963) |
| 5 | 719.5898817099219 | between Caribbean (long: -75.81834; lat: 14.525556) and North America (long: -105.25512; lat: 54.525963) |
| 6 | 394.2296542882096 | between Europe (long: 15.255119; lat: 54.525963) and South Asia (long: 76.45631; lat: 25.03764) |
| 7 | 275.169855862822 | between Central Asia (long: 68.8319; lat: 45.450687) and South America (long: -55.491478; lat: -8.783195) |
| 8 | 242.58395188106397 | between Africa (long: 34.508522; lat: -8.783195) and Middle East (long: 42.55096; lat: 29.298529) |
| 9 | 215.1360893201474 | between Central Asia (long: 68.8319; lat: 45.450687) and North Asia (long: 128.60144; lat: 35.871433) |
| 10 | 210.2262491135267 | between Middle East (long: 42.55096; lat: 29.298529) and South America (long: -55.491478; lat: -8.783195) |
| 11 | 197.5575786097743 | between Europe (long: 15.255119; lat: 54.525963) and Middle East (long: 42.55096; lat: 29.298529) |
| 12 | 175.13511087787467 | between Middle East (long: 42.55096; lat: 29.298529) and North America (long: -105.25512; lat: 54.525963) |
| 13 | 154.70390808282224 | between Caribbean (long: -75.81834; lat: 14.525556) and Middle East (long: 42.55096; lat: 29.298529) |
| 14 | 153.9414093857853 | between Central Asia (long: 68.8319; lat: 45.450687) and South Asia (long: 76.45631; lat: 25.03764) |
| 15 | 149.75592120854537 | between Caribbean (long: -75.81834; lat: 14.525556) and South America (long: -55.491478; lat: -8.783195) |
| 16 | 148.85810729494116 | between Africa (long: 34.508522; lat: -8.783195) and Caribbean (long: -75.81834; lat: 14.525556) |
| 17 | 141.7288507486571 | between Oceania (long: 140.01877; lat: -22.73591) and South Asia (long: 76.45631; lat: 25.03764) |
| 18 | 122.99363732018385 | between Africa (long: 34.508522; lat: -8.783195) and South America (long: -55.491478; lat: -8.783195) |
| 19 | 108.96046044668044 | between Africa (long: 34.508522; lat: -8.783195) and South Asia (long: 76.45631; lat: 25.03764) |
| 20 | 93.95724953344795 | between Middle East (long: 42.55096; lat: 29.298529) and South East Asia (long: 115.66283; lat: -2.21797) |
| 21 | 89.77169188956839 | between Middle East (long: 42.55096; lat: 29.298529) and North Asia (long: 128.60144; lat: 35.871433) |
| 22 | 79.73665991873779 | between Middle East (long: 42.55096; lat: 29.298529) and Oceania (long: 140.01877; lat: -22.73591) |
| 23 | 79.34475578878079 | between Central Asia (long: 68.8319; lat: 45.450687) and North America (long: -105.25512; lat: 54.525963) |
| 24 | 69.36887618230472 | between Europe (long: 15.255119; lat: 54.525963) and South America (long: -55.491478; lat: -8.783195) |
| 25 | 68.30881586223244 | between North America (long: -105.25512; lat: 54.525963) and South America (long: -55.491478; lat: -8.783195) |
| 26 | 65.66861355850432 | between Caribbean (long: -75.81834; lat: 14.525556) and Europe (long: 15.255119; lat: 54.525963) |
| 27 | 58.798278757864665 | between North America (long: -105.25512; lat: 54.525963) and Oceania (long: 140.01877; lat: -22.73591) |
| 28 | 57.920159733205224 | between Caribbean (long: -75.81834; lat: 14.525556) and Central Asia (long: 68.8319; lat: 45.450687) |
| 29 | 54.72909826757337 | between Africa (long: 34.508522; lat: -8.783195) and Central Asia (long: 68.8319; lat: 45.450687) |
| 30 | 48.409207198045934 | between Caribbean (long: -75.81834; lat: 14.525556) and Oceania (long: 140.01877; lat: -22.73591) |
| 31 | 47.676032823984976 | between Europe (long: 15.255119; lat: 54.525963) and North Asia (long: 128.60144; lat: 35.871433) |
| 32 | 43.669076886379976 | between South America (long: -55.491478; lat: -8.783195) and South East Asia (long: 115.66283; lat: -2.21797) |
| 33 | 41.75069653855484 | between Africa (long: 34.508522; lat: -8.783195) and South East Asia (long: 115.66283; lat: -2.21797) |
| 34 | 37.61418225410232 | between Central Asia (long: 68.8319; lat: 45.450687) and South East Asia (long: 115.66283; lat: -2.21797) |
| 35 | 35.41858620604572 | between North Asia (long: 128.60144; lat: 35.871433) and South East Asia (long: 115.66283; lat: -2.21797) |
| 36 | 32.056559123090025 | between North America (long: -105.25512; lat: 54.525963) and South Asia (long: 76.45631; lat: 25.03764) |
| 37 | 30.68412462268806 | between North America (long: -105.25512; lat: 54.525963) and North Asia (long: 128.60144; lat: 35.871433) |
| 38 | 25.738035527097743 | between Central Asia (long: 68.8319; lat: 45.450687) and Europe (long: 15.255119; lat: 54.525963) |
| 39 | 23.252220729011807 | between South Asia (long: 76.45631; lat: 25.03764) and South East Asia (long: 115.66283; lat: -2.21797) |
