## Supplementary material for "Next-Gen sequencing of novel pandemic swine flu [A(H1N1)pdm09] virus in India revealed novel mutations across the genome": Bayesian test statistics

| 0 | pk=0.9998866652741779 | BF=36555.5343265314 | between Africa (long: 34.508522; lat: -8.783195) and Europe (long: 15.255119; lat: 54.525963) |
| --- | --- | --- | --- |
| 1 | pk=0.9974728103256655 | BF=1635.417293719311 | between Central Asia (long: 68.8319; lat: 45.450687) and Middle East (long: 42.55096; lat: 29.298529) |
| 2 | pk=0.9971817606111538 | BF=1466.0939575567506 | between South America (long: -55.491478; lat: -8.783195) and South Asia (long: 76.45631; lat: 25.03764) |
| 3 | pk=0.996833324949911 | BF=1304.3207582213108 | between Africa (long: 34.508522; lat: -8.783195) and North America (long: -105.25512; lat: 54.525963) |
| 4 | pk=0.9942748513836553 | BF=719.5898817099219 | between Caribbean (long: -75.81834; lat: 14.525556) and North America (long: -105.25512; lat: 54.525963) |
| 5 | pk=0.989598994779503 | BF=394.2296542882096 | between Europe (long: 15.255119; lat: 54.525963) and South Asia (long: 76.45631; lat: 25.03764) |
| 6 | pk=0.985165473635778 | BF=275.169855862822 | between Central Asia (long: 68.8319; lat: 45.450687) and South America (long: -55.491478; lat: -8.783195) |
| 7 | pk=0.983206240947478 | BF=242.58395188106397 | between Africa (long: 34.508522; lat: -8.783195) and Middle East (long: 42.55096; lat: 29.298529) |
| 8 | pk=0.9811041172085694 | BF=215.1360893201474 | between Central Asia (long: 68.8319; lat: 45.450687) and North Asia (long: 128.60144; lat: 35.871433) |
| 9 | pk=0.9806713333216202 | BF=210.2262491135267 | between Middle East (long: 42.55096; lat: 29.298529) and South America (long: -55.491478; lat: -8.783195) |
| 10 | pk=0.9794573163510322 | BF=197.5575786097743 | between Europe (long: 15.255119; lat: 54.525963) and Middle East (long: 42.55096; lat: 29.298529) |
| 11 | pk=0.9768880321087506 | BF=175.13511087787467 | between Middle East (long: 42.55096; lat: 29.298529) and North America (long: -105.25512; lat: 54.525963) |
| 12 | pk=0.9739153341907956 | BF=154.70390808282224 | between Caribbean (long: -75.81834; lat: 14.525556) and Middle East (long: 42.55096; lat: 29.298529) |
| 13 | pk=0.973789518727749 | BF=153.9414093857853 | between Central Asia (long: 68.8319; lat: 45.450687) and South Asia (long: 76.45631; lat: 25.03764) |
| 14 | pk=0.9730766916610589 | BF=149.75592120854537 | between Caribbean (long: -75.81834; lat: 14.525556) and South America (long: -55.491478; lat: -8.783195) |
| 15 | pk=0.9729187055833536 | BF=148.85810729494116 | between Africa (long: 34.508522; lat: -8.783195) and Caribbean (long: -75.81834; lat: 14.525556) |
| 16 | pk=0.9715951544129071 | BF=141.7288507486571 | between Oceania (long: 140.01877; lat: -22.73591) and South Asia (long: 76.45631; lat: 25.03764) |
| 17 | pk=0.9674093520976172 | BF=122.99363732018385 | between Africa (long: 34.508522; lat: -8.783195) and South America (long: -55.491478; lat: -8.783195) |
